# Extracting abundance information from DNA-based data

**DOI:** 10.1101/2022.01.06.475221

**Authors:** Mingjie Luo, Yinqiu Ji, David Warton, Douglas W. Yu

## Abstract

The accurate extraction of species-abundance information from DNA-based data (metabarcoding, metagenomics) could contribute usefully to diet analysis and food-web reconstruction, the inference of species interactions, the modelling of population dynamics and species distributions, the biomonitoring of environmental state and change, and the inference of false positives and negatives. However, multiple sources of bias and noise in sampling and processing combine to inject error into DNA-based datasets. We focus here on the laboratory and bioinformatic processes of generating DNA-based data, since sampling bias and noise are addressed extensively in the ecological literature. To extract abundance information, it is useful to distinguish two concepts. (1) *Within-sample across-species* quantification describes relative species abundances within one sample. (2) *Across-sample within-species* quantification describes how the abundance of each individual species varies from sample to sample, as in a time series, an environmental gradient, or experimental treatments. First, we review the literature on methods to recover (1) *across-species* abundance information (which is achieved by removing what we call ‘species pipeline biases’) and (2) *within-species* abundance information (by removing what we call ‘pipeline noise’). We argue that many ecological questions can be answered by extracting only within-species quantification, and we therefore demonstrate how to use a ‘DNA spike-in’ to correct for pipeline noise and recover within*-species* abundance information. We also introduce a model-based estimator that can be employed on datasets without a physical spike-in to approximately estimate and correct for pipeline noise.

## Introduction

The accurate extraction of species-abundance information from DNA-based data could contribute usefully to the reconstruction of diets and quantitative food webs, the inference of species interactions, the modelling of population dynamics and species distributions, the biomonitoring of environmental state and change, and more prosaically, the inference of false positives and negatives (Thomas et al. 2016; Deagle et al. 2019; Peel et al. 2019; Carraro et al. 2020, 2021; Abrego et al. 2021; Rojahn et al. 2021). *Here we use the term abundance to mean any estimate of biomass or count of individuals*.

However, there are four general obstacles to the extraction of abundance information from DNA-based data (see Shelton et al. 2016; Griffin et al. 2020 for more formal treatments), which we will call here: (1) *species capture biases*, (2) *capture noise*, (3) *species pipeline biases*, and (4) *pipeline noise*.

1. *Species capture biases*. – Different species are more or less likely to be captured by a given sampling method or via non-random sampling designs. For instance, Malaise traps preferentially capture Diptera (deWaard et al. 2019), and different fish species, body sizes, and physiological conditions vary in their eDNA shedding rates (Thalinger et al. 2021; Yates et al. 2021b).
2. *Capture noise*. – Steinke et al. (2021) have shown that Malaise traps separated by only 3 m fail to capture the same species compositions, from which we infer that abundances vary stochastically across traps. Levi et al. (2019) showed that counts of salmon could be estimated via quantitative PCR of aquatic environmental DNA, but only after correcting for temporal fluctuations in streamflow. Other sources of capture noise include environmental variation in eDNA degradation rates, food availability, PCR inhibitors, and transport rates (reviewed in Yates et al. 2021a).
3. *Species pipeline biases*. – Species differ in body size (biomass bias), genome size, mitochondrial copy number, DNA extraction efficiency, and PCR amplification efficiency (primer bias) (Amend et al. 2010; Yu et al. 2012; Elbrecht and Leese 2015; Piñol et al. 2015, 2019; Tang et al. 2015; Bell et al. 2017; Krehenwinkel et al. 2017; McLaren et al. 2019; Pauvert et al. 2019; Garrido-Sanz et al. 2021; Yang et al. 2021; Iwaszkiewicz-Eggebrecht et al. 2022). Species can even differ in their propensity to survive a bioinformatic pipeline, such as when closely related species are clustered into one operational taxonomic unit (Pauvert et al. 2019).
4. *Pipeline noise*. – There is considerable noise in DNA-based pipelines, which breaks the relationship between starting sample biomasses and final numbers of reads per sample (Ji et al. 2020), caused in part by PCR stochasticity and the passing and pooling of small aliquots of liquid along wet-lab pipelines. In particular, it is common practice *to deliberately equalise the amount of data per sample* by “pooling samples in equimolar concentration” just before sequencing.

We do not consider species capture biases or capture noise further, referring the reader to the literature on eDNA occupancy correction (e.g. Dorazio and Erickson 2018; Doi et al. 2019; Erickson 2019; Griffin et al. 2020; Lyet et al. 2021; Stauffer et al. 2021) and the review by Yates et al. (2021a). Instead, our purpose is to review methods for the extraction of abundance information from already-collected samples, because even if species capture biases and capture noise can be corrected, the combination of species pipeline biases and pipeline noise still *causes the number of DNA sequences assigned to a species in a sample to be an error-prone measure of the abundance of that species in that sample* (McLaren et al. 2019).

To start, we illustrate in a simplified way how pipeline noise and species pipeline biases (hereafter, species biases) combine to inject error into DNA-based datasets. We start with a notionally true sample X species table or OTU table (Figure 1), where OTU stands for Operational Taxonomic Unit, i.e. a species hypothesis. Let each cell represent the true abundance (biomass or count) of that OTU in that sample.

**Figure 1.**
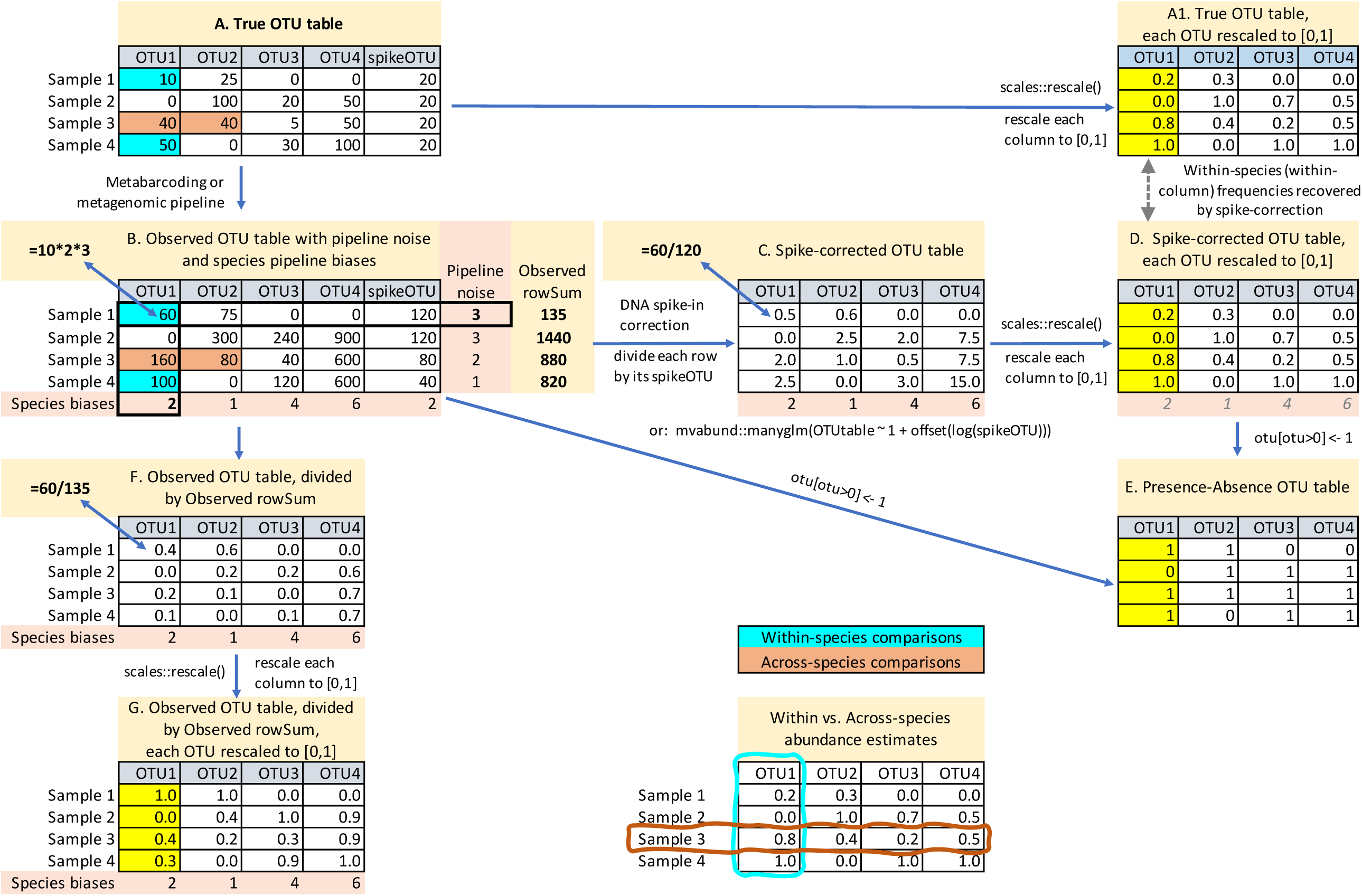
Pipeline noise versus species bias in OTU tables. **A**. The true OTU table, with cell numbers representing the true abundance of DNA for each OTU (column) in each sample (row). The spikeOTU column shows that the same amount of DNA spike-in has been added to each sample. **A1**. The true OTU table after rescaling each OTU column to the interval [0,1]. **B**. The observed OTU table after amplicon sequencing, showing the combined effects of pipeline noise and species biases. Each cell in Table A is multiplied by the Pipeline noise and Species bias values in that cell’s row and column. For instance, OTU1’s true abundance in Sample 1 is 10 but appears as 60 (=10*2*3). Pipeline noise thus causes the original 10:50 ratio of OTU1 in Samples 1 and 4 (blue cells) to appear as 60:100, while species bias causes the original 40:40 ratio of OTU1 and OTU2 (orange cells) to appear as 160:80. **C**. The observed OTU table after dividing each row by its observed spike-in reads, which removes pipeline noise. Note that species biases remain uncorrected. In statistical modelling, the observed spike-in values are an index of sampling (sequencing) effort and can be included as offset values. **D**. Table C after rescaling each column to the interval [0,1], to allow direct comparison with the rescaled true-OTU Table A1. Spike-in correction successfully recovers *within-species* abundance change from sample to sample. Species biases have not been removed but have now been ignored via rescaling. **E**. If spike-in reads are not available, or if it is suspected that capture noise is uncorrectable and high, the observed OTU table can be transformed to presence/absence. However, this method loses ecological information (Figure 2C). **F.** Pipeline noise cannot be reliably removed by using the total reads per sample as a proxy for sampling effort (Observed rowSum) because the observed rowSum is confounded by species composition. **G**. Table F after rescaling each OTU column to the interval [0,1], to contrast with Tables A1 and D. Line graphs of the OTU tables are in the spreadsheet version of this table, in Supplementary Information. Code syntax from the R language (R Core Team 2021), including the {mvabund} (Wang et al. 2012) and {scales} packages (https://scales.r-lib.org, accessed 16 Dec 2021). (**374 words**)

**Table 1.** Summary of reviewed methods for extracting abundance information from DNA-based data. Each method is scored for whether it can achieve within-species or across-species quantification or both.

| Method | Description | Within-species abundance | Across-species abundance |
| --- | --- | --- | --- |
| Multiplexed individual barcoding | DNA-barcode every individual in every sample | ✓ | ✓ |
| Presence-absence in multiple sub-samples | Take multiple subsamples and count presences | ✓ | ? |
| Design less biased PCR primers | Self-explanatory |  | ✓ |
| Quantitative/Digital-Droplet PCR | Quantify a species' DNA concentration per sample | ✓ | ✓ (with extra work) |
| Spike-in DNA | Add a fixed amount of external DNA to each sample to measure pipeline noise | ✓ |  |
| Model-based pipeline-noise estimation | Estimate the effect of pipeline noise by removing the effect of environmental predictors | ✓ |  |
| Unique Molecular Identifiers (UMIs) | Estimate the amount of starting DNA per sample and per species | ✓ | ? |
| Estimate and eliminate PCR bias | Use calibration samples and/or PCR time series to estimate species-specific PCR biases |  | ✓ |
| Forward and reverse metagenomics | Map and count shotgun reads to reference sequences |  | ✓ |

Pipeline noise affects the *rows* (samples) of an OTU table. Thus, even though in the true table, OTU1 is six times as abundant in sample 4 versus sample 1 (green cells in Figure 1 A), in the observed table, OTU1 is only *two* times as abundant in sample 4 (green cells in Figure 1 B). *Pipeline noise thus obscures how the abundance of each individual species varies across samples, where the samples could be a time series, an environmental gradient, or different experimental treatments*.

Species bias affects the *columns* (OTUs) of an OTU table. Thus, even though in the true table, OTU2 and OTU1 are equally abundant in sample 3 (orange cells in Figure 1 A), in the observed OTU table, OTU2 is two times as abundant as OTU1 in sample 3 (orange cells in Figure 1 B). Species bias thus obscures *relative species abundances*, which is important for diet analysis (Deagle et al. 2019) and when relative abundance within a sample provides information on species contribution to ecosystem functioning or services (e.g. relative fish species biomasses).

So how can we extract abundance information from DNA-based data? It is helpful to distinguish between two concepts from Ji et al. (2020; see also Garrido-Sanz et al. 2021):

1. *Within*-species quantification: E.g. “Species A is more abundant in this sample than it is in that sample (e.g. two points on a time series).” This is achieved by removing pipeline noise (Figure 2 A1, D).
2. *Across*-species quantification: E.g. “Species A is more abundant than Species B *in this sample* (i.e. relative species abundance).” This is achieved by removing species biases.

We can state this mathematically as:

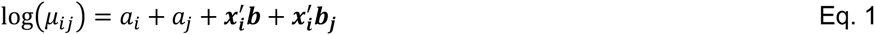

where *μ_ij_* is the abundance of species *j* in sample *i*, *α_i_* is a measure of the overall abundance of a sample, *α_j_* is a measure of how abundant species *j* is across samples, and we assume a vector of environmental variables ***x_i_*** (whose transpose is ***x′_i_***) have an effect on total abundance (via ***b***) as well as having a compositional effect, *i.e.* affecting different species in different ways (via, ***b_j_***). The responses to environmental variables (ssume a vector of environmental varia ***b*** and, ***b_j_***) are typically the main quantities of biological interest, being used to model and monitor species distributions. Pipeline noise biases our estimate of *α_i_*, which would be zero for identical replicates in the absence of stochasticity, which in turn biases estimates of effects of environmental variables (***b*** and, ***b_j_***). Species pipeline biases affect our estimate of *α_i_*, affecting across-species quantification.

**Figure 2.**
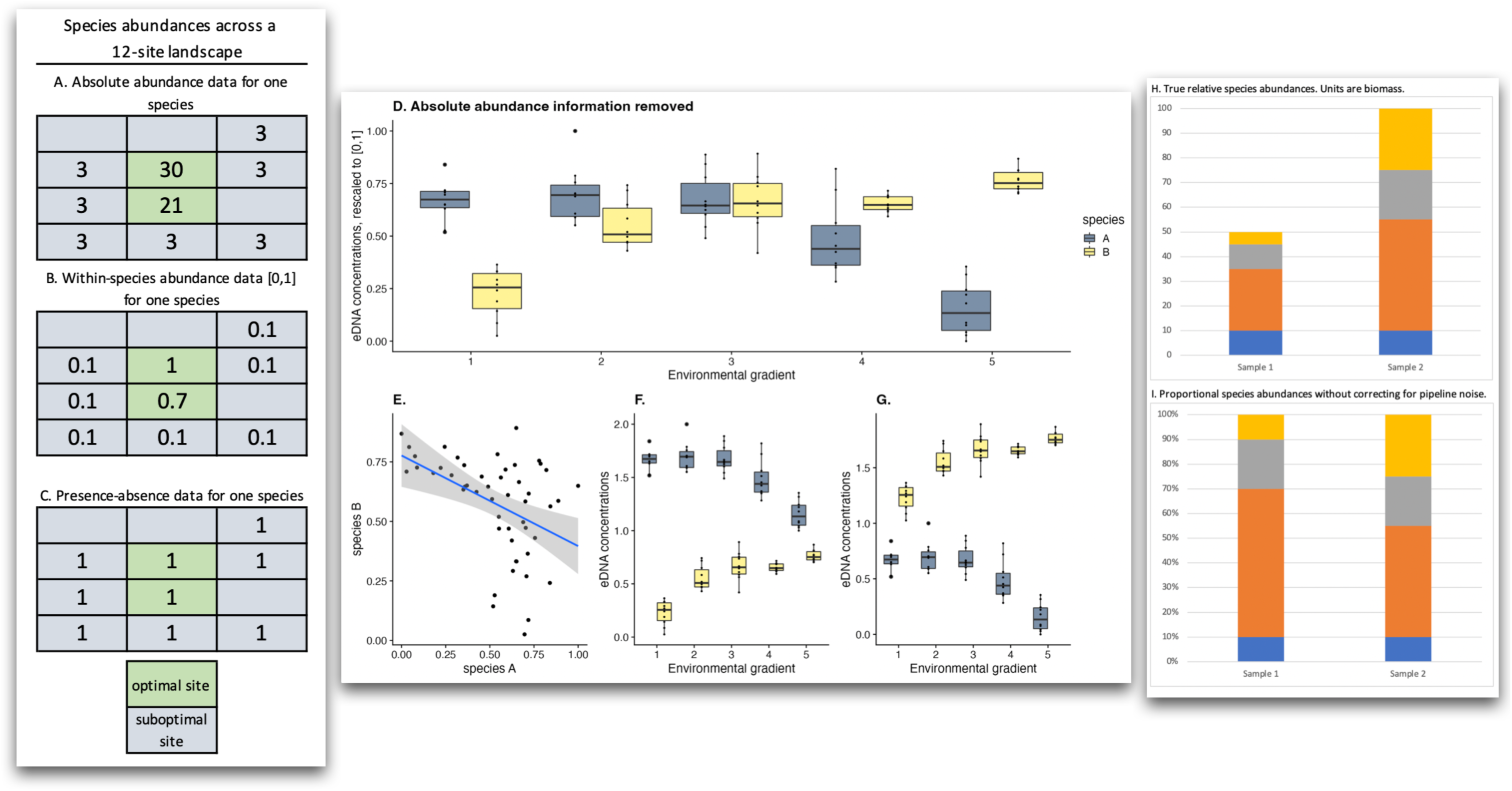
The usefulness of within-species abundance information. **A.** Imagine that a species is found in many sites but that only two sites are optimal (green cells) with high abundances, with the rest suboptimal (grey cells), with low abundances. **B**. Even though species pipeline biases make it difficult to recover absolute abundances from DNA-based data, it is straightforward to use a spike-in to recover within-species abundance data (shown by rescaling to [0,1]), still revealing that the green sites appear to be optimal habitat. **C**. If the DNA-based data were converted to presence/absence, the distinction between green and grey habitat would be lost. **D**. Along an environmental gradient from left to right, let species A decrease and species B increase in eDNA concentration (rescaled to [0,1]). **E**. The two species are seen to be negatively correlated over the gradient, even though absolute abundance information is unavailable. Example adapted from Rojahn et al. (2021) who combined a similar result with additional information to infer the competitive exclusion of a native fish species by an invasive species. **F**. **G**. Due to species pipeline biases, absolute abundances are not known, and either species A or B could be absolutely more abundant. **H**. Two samples with different absolute and relative species abundances. **I**. If only across-species quantification is achieved (e.g. via forward or reverse metagenomics), it is valid to compare across species *within* each sample only, revealing that the dark orange species has the highest relative abundance in both samples 1 and 2. However, it would not be valid to conclude that the dark orange species has a greater absolute biomass in Sample 1 than in Sample 2, as can be seen by inspection of the true absolute abundances in **H**. However, also achieving within-species quantification (via a spike-in) would make it possible to compare how each species’ *absolute* abundances vary across samples. (**308 words**)

As we review and demonstrate below, some approaches remove pipeline noise, some remove species biases, and some remove both*. Our take-home message is that removing only pipeline noise to achieve within-species quantification can be enough* to improve the inference of species interactions, the modelling of population dynamics and species distributions, the biomonitoring of environmental state and change, and the inference of false positives and negatives (Carraro et al. 2020, 2021; Abrego et al. 2021; Rojahn et al. 2021, and Figure 2).

### Mini-review of methods to extract abundance information

#### Multiplexed individual barcoding’

The most straightforward approach is to DNA-barcode all the individual organisms and count them up, which achieves both within- and across-species quantification. This method only works on taxa that have body sizes suitable for separating individuals, like bees (Gueuning et al. 2019). Once separated, individuals or portions thereof (like a leg) are placed in separate wells of a 96-well plate and individually PCR’d. Each PCR requires a uniquely tagged pair of PCR primers, which allows all the PCR products to be pooled and then sequenced *en masse* on Illumina (Meier et al. 2016; Ratnasingham 2019; Creedy et al. 2020), PacBio (Hebert et al. 2018), or MinION (Srivathsan et al. 2021). This method now costs much less than $1 per individual. Wührl et al. (2021) further increase throughput with a robotic pipettor and camera that visually identifies small insects to higher taxonomic rank and sorts them into 96-well plates. However, this method is difficult to apply to very large numbers of individuals and cannot be applied to trace DNA or microbial taxa. Note that this approach could also be carried out via machine-learning-accelerated visual identifications of photos of arthropods (Schneider et al. 2022).

#### Presence-absence in multiple subsamples

Presence-absence across multiple subsamples can be used as an index of within-species abundance. For instance, Abrego et al. (2021) summed all weekly detections (presences) per species in their mitogenomic arthropod dataset to estimate an annual abundance measure for each species. However, pipeline noise can still be reflected in presence/absence data, albeit more weakly, especially when many subsamples are used. This method can achieve partial within-species quantification but probably not across-species quantification.

#### Design less biased PCR primers

In some cases, the target taxon is nearly uniform in body size and DNA-extraction efficiency, and it can be possible to design PCR primers that bind similarly across species. For instance, Schenk et al. (2019) have reported that primers for the 28S D3-D5 and 18S V4 regions return nematode read frequencies that accurately recover relative species abundances, Verkuil et al. (2020) have reported that modified COI primers can recover the relative biomasses of insect orders from Pied Flycatcher faeces, and Ershova et al. (2021) have reported that increasing COI primer degeneracy (Leray-XT) can recover relative biomasses of marine zooplankton. This method achieves across-species quantification, albeit with error, but not within-species quantification.

#### Quantitative/Digital-Droplet PCR

qPCR and ddPCR (quantitative and digital droplet PCR) can be used to estimate the sample DNA concentration of one species per assay. ddPCR is more sensitive than is qPCR (Brys et al. 2021) and allows the detection of single copies of target DNA and absolute quantification through the partitioning of the PCR reaction into 20,000 droplets and subsequent fluorescent detection of droplets that contain the target DNA (Hindson et al. 2011). This paper does not review q/ddPCR except to note that single-species q/ddPCR applied to aquatic trace DNA can achieve within-species quantification, provided that one corrects for capture bias and noise in the form of variation in water discharge rates, surface-area to mass ratio and/or eDNA transport and diffusion (Levi et al. 2019; Pochardt et al. 2020; Fukaya et al. 2021; Yates et al. 2021b, 2021a; Rourke et al. 2022; Shelton et al. 2022b). If applied to multiple species and if statistical models that relate DNA copy number to abundance can be fitted (Levi et al. 2019; Pochardt et al. 2020; Fukaya et al. 2021), then across-species quantification can also be achieved, albeit with non-trivial amounts of error. See also Rourke et al. (2022) for a recent, comprehensive review.

#### Spike-in DNA

To achieve within-species quantification, researchers have advocated adding a fixed amount of an arbitrary DNA sequence to each sample, after tissue lysis and before DNA extraction. This ‘spike-in’, also known as an internal standard (ISD, Harrison et al. 2021), must have a sequence that does not match any species that could be in the samples and be flanked by primer binding sequences that match the primers used to amplify the samples (Smets et al. 2016; Deagle et al. 2018; Tkacz et al. 2018; Ushio et al. 2018; Harrison et al. 2021; Tsuji et al. 2022). By design, each sample receives the same amount of spike-in, and all samples should therefore return the same number of spike-in reads after PCR and sequencing. However, due to pipeline noise, some samples return more spike-in reads because more of the sample’s DNA made it through the metabarcoding pipeline; those samples have OTUs with ‘too many reads’. Some samples return fewer spike-in reads because less of the sample’s DNA made it through the metabarcoding pipeline; those samples have OTUs with ‘too few reads’. The correction step is simple: divide each sample’s OTU sizes by the number of spike-in reads in that sample (Ji et al. 2020; Abrego et al. 2021). OTUs in samples with large numbers of spike-in reads are reduced in size more than OTUs in samples with small numbers of spike-in reads. Alternatively, the number of spike-in reads per sample can be input as an offset term in a multivariate statistical model (Wang et al. 2012). This latter approach can be understood as estimating *α_i_* in Eq. 1 using 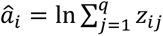 where we have spike-in reads (*z_ij_*) for *q* species (or synthetic sequences).

As an example, and following the pioneering work of Zhou et al. (2013), Ji et al. (2020) mapped whole-genome-sequenced (WGS, aka ‘shotgun sequencing’) datasets of insects to mitochondrial genomes and barcodes and achieved nearly perfect within-species quantification (barcodes R^2^ = 93%, mitogenomes R^2^ = 95%) and almost direct proportionality between mapped reads and input DNA-mass. The high accuracy was largely achieved by employing a spike-in correction. However, the regression lines that related read number to input DNA for each species all had different intercepts, reflecting uncorrected species biases, and thus across-species quantification was not achieved. Harrison et al. (2021) provide an excellent, complementary review of the recent literature on spike-ins and also describe an alternative approach for modelling non-spike-corrected (‘compositional’) datasets. Figure 1 provides a worked example of spike-in correction, and the Excel spreadsheet used to produce Figure 1 is in Supplementary Materials.

#### Model-based pipeline-noise estimation

A related approach is to try to use the data itself to estimate the pipeline noise, rather than a physical spike-in. To do this we could fit the model stated in Eq. 1 to data. However, fitting this full model with row effects can be computationally intensive, especially for large datasets, so a simple alternative is to approximate *α_i_* using a one-step estimator (Warton 2022):

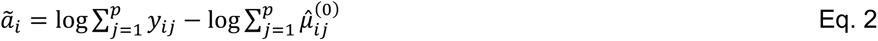

where *y_ij_* is the number of reads for OTU *j* in sample 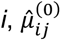 is its predicted value from a model that does not include a row effect, and 7 is the total number of OTUs. We can then include *α_i_* as an offset in future models to (approximately) correct for pipeline bias.

The reason Equation 2 has two terms in it is that there are two reasons that a sample might end up generating many sequence reads: by chance (pipeline noise) and/or because some (or many) of the OTUs are abundant in the site where the sample was taken (ecology). Thus, if one has informative predictors ***x****_i_* that can successfully predict which OTUs should be abundant in which samples, then it becomes possible to separate the two effects. 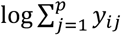 is a function of both effects, 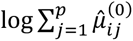 estimates the effect of the predictors on the OTUs (ecology), and their difference isolates the row effect (pipeline noise). This is related to the spike-in approach, the main difference being that the spike-in formula (for ***α****_i_*) has no second term involving *α_i_* since, by design, the same amount of each spike-in species is included in every sample (the spike-in has no ecology). An important difference here however is that because the same data are being used to estimate both pipeline noise (***α****_i_*) and ecological effects (***b***), it will be difficult to tease these effects apart if the two are correlated. In fact, the common practice of adjusting samples to equimolar concentration before sequencing confounds these two effects. This problem does not however affect estimation of compositional effects (***b****_j_*), often the main quantity of interest.

#### Unique Molecular Identifiers (UMIs)

A UMI is a series of ∼7-12 random bases (‘NNNNNNN’) added to the forward primer as an ultra-high diversity tag (Hoshino and Inagaki 2017). Seven Ns produce 4^7^ = 16 384 uniquely identified forward primer molecules. Species contributing abundant DNA to a sample will capture many of these primer molecules and thus amplify many unique UMIs, while species contributing scarce DNA will amplify a low number. The relationship between UMI richness and DNA abundance is roughly linear but asymptotes for species with very high DNA abundance. After sequencing, the number of UMIs per OTU correlates with the starting number of template DNA molecules per species in that sample (Hoshino and Inagaki 2017; Hoshino et al. 2021). This method thus mimics q/ddPCR in that if statistical relationships between DNA copy number and true abundance can be estimated, across-species quantification can be achieved. Within-species quantification can be achieved by also adding a spike-in.

#### Estimate and eliminate PCR bias

Silverman et al. (2021) propose a straightforward way to estimate PCR bias, by pooling all samples to ensure that all species are present, and subjecting the pooled sample to different numbers of PCR cycles *x_i_*, from low to high. For any given pair of species 1 and 2, the ratio of their reads 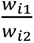 after a given number of cycles is their starting DNA ratio 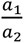 multiplied by their relative amplification bias 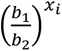, which increases with the number of cycles. This relationship can be linearised, and given the post-PCR relative read numbers at all cycle numbers, starting DNA ratios (and relative amplification biases) can be estimated.’

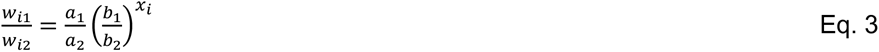

However, PCR is not the only source of species pipeline bias (e.g. Iwaszkiewicz-Eggebrecht et al. 2022), and McLaren et al. (2019) have pointed out that although it is not possible to estimate *a priori* the whole set of species biases in a given amplicon or metagenomic dataset (because an unknown number of factors of unknown strengths combine to create the biases), it is reasonable to assume that *the ratio of the biases of every pair of species is fixed*. Given this, Williamson et al. (2021; see also Clausen and Willis 2022) showed that if first one is able to estimate the absolute abundances of a *subset* of species in the samples (via multiple, species-specific q/ddPCR assays or flow cytometry), it is possible to infer the absolute abundances of all the species by inferring their ratios with the q/ddPCR-quantified species, allowing one to achieve across-species quantification. The authors dub this a ‘multiview data structure’ because there are two views into the community of interest: q/ddPCR and sequencing. Note that because q/ddPCR is carried out after many of the wet-lab steps have been carried out, multiview modelling does not remove pipeline noise, and a spike-in is still needed to achieve within-species quantification. Shelton et al. (2022a) advocate a similar approach but via the construction of a ‘mock community’ containing tissue of all species of interest and subjecting it to the same PCR protocol as the samples. From this mock community, species-specific PCR biases are calculated and used to extract across-species abundance information.

#### Forward and reverse metagenomics

Another way to achieve across-species quantification is to avoid PCR by using a metagenomic approach. For marine phytoplankton, Pierella Karlusich et al. (2022) used shotgun-sequenced counts of the (mostly) single-copy *psbO* gene, which is part of the photosystem II complex, to estimate species relative abundances. For land plants, Lang et al. (2019) showed that WGS datasets from pollen samples mapped to the variable protein-coding regions in chloroplast genomes can achieve accurate across-species quantification, finding that read frequency correlated strongly and linearly with pollen-grain frequency in a nearly 1:1 relationship (R^2^ = 86.7%, linear regression). At the same time, Peel et al. (2019) showed that it is possible to skip the labour of assembling and annotating chloroplast genomes, by using long-read sequences produced by the MinION sequencers from Oxford Nanopore Technologies (ONT). In this protocol, unassembled genome skims of individual plant species, ideally sequenced at ≥1.0X depth, are used as reference databases. Mixed-species query samples of pollen are sequenced on MinIONs. The reads from each (reference) genome skim are mapped to each (query) long read, and each long read is assigned to the species whose skim maps at the highest percent coverage. This ‘reverse metagenomic’ (RevMet) protocol achieves across-species quantification, allowing biomass-dominant species to be identified in mixed-species pollen samples (and potentially, in root masses). Because RevMet uses the whole genome, it avoids species biases and pipeline noise created by ratios of chloroplasts to cells varying across species, condition, tissues, and age, and it can potentially be applied to any taxon for which it is possible to generate large numbers of individual genome skims, potentially including soil fauna. However, metagenomics by itself does not remove pipeline noise and would have to be paired with a spike-in to achieve within-species quantification.

To sum up, multiple methods exist to extract abundance information from DNA-based datasets (Table 1). Some achieve within-species quantification by removing pipeline noise, some achieve across-species quantification by removing species biases, and some achieve both or can be combined to achieve both.

It is useful to understand that many ecological questions can be tackled with only within-species quantification (Figure 2). In the second half of this paper, we therefore provide a detailed protocol and experimental validation of spike-ins to achieve within-species quantification for metabarcoding datasets.

We carry out two tests. First, we start with a sample of known composition (a ‘mock soup’ of 52 OTUs), and from this, we create a dilution gradient of 7 samples with a spike-in. We show the successful use of the spike-in correction to remove pipeline noise and recover the dilution gradient. We then repeat the experiment with seven Malaise trap samples, which have the advantage of being more realistic but the disadvantage of having unknown compositions.

Again, we show the successful use of the spike-in to recover the seven dilution gradients made from the seven samples.

## Methods

### 2.1 Mock soup construction

286 arthropods were collected in Kunming, China (25°8’23” N, 102°44’17” E) (Luo et al. 2022). DNA was extracted from each individual using the DNeasy Blood & Tissue Kit (Qiagen GmbH, Germany). Genomic DNA concentration of each individual was quantified from three replicates using PicoGreen fluorescent dye. 658-bp COI barcoding sequences were PCR’d with Folmer primers (LCO1490 and HCO2198) (Folmer et al. 1994) and Sanger-sequenced. After the 658-bp COI sequences were trimmed to 313 bp based on our metabarcoding primers (see *2.4 Primer design*), 286 arthropods were clustered to 168 OTUs at 97% similarity. We selected 52 individuals with genomic DNA > 20 ng/µl, representing 52 OTUs.

We created a mock-soup gradient of seven dilution levels. First, we created the highest concentration-level soup by pooling 61 ng of each of the 52 OTUs. The next soup was created by pooling 48.8 ng (= 0.8 x 61) of each of the 52 OTUs, and so on to create a gradient of seven mock soups of differing absolute abundances, stepping down 0.8X each time. To make it possible to check for mundane experimental error (as opposed to failure of the spike-in to recover the gradient), we independently created this mock-soup gradient three times, for ntot = 21 independent poolings (Figure 3 A).

**Figure 3.**
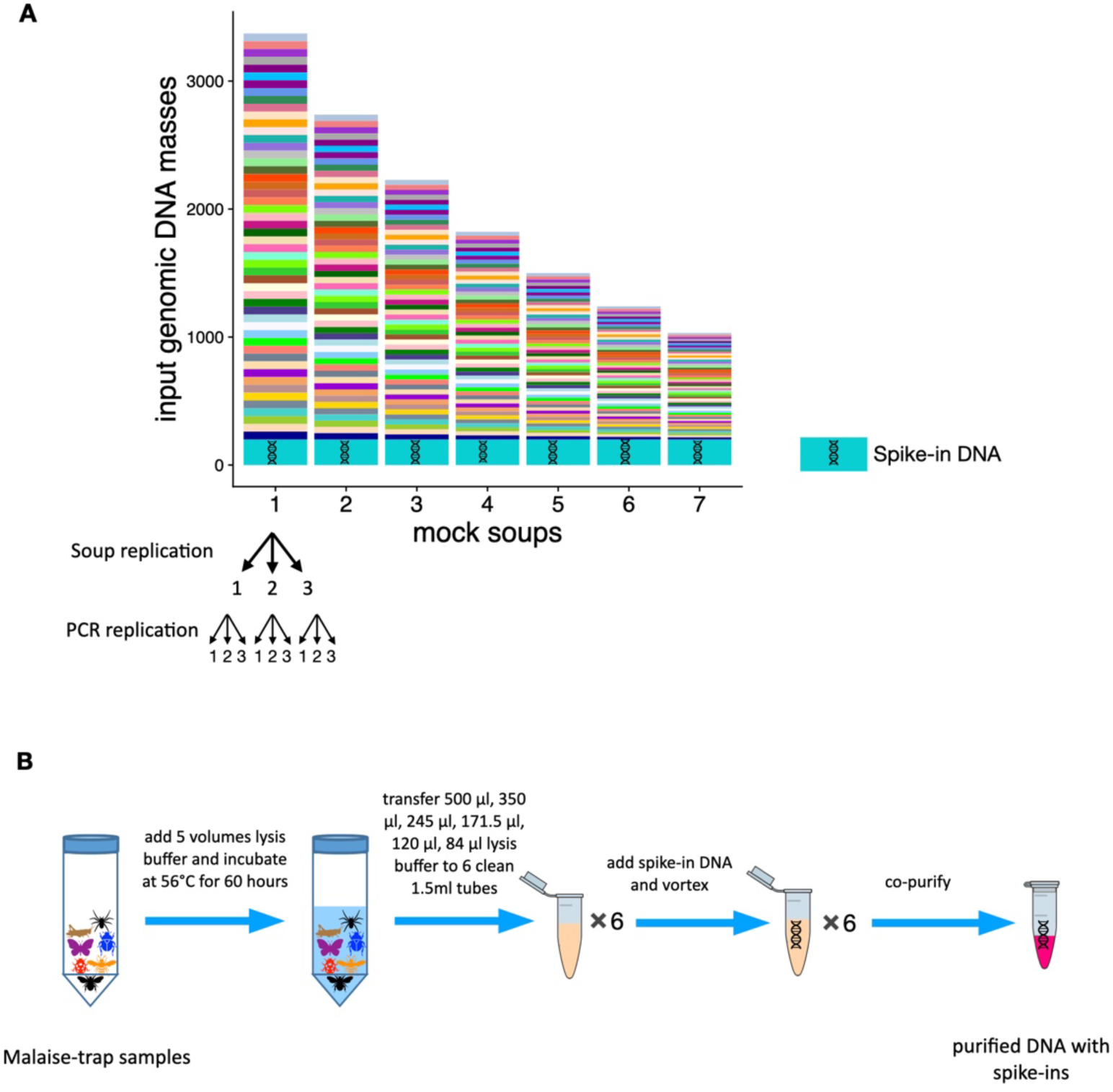
Preparation of mock and Malaise-trap soups. **A**. Mock soups. Each mock soup was constructed with equal masses of purified DNA from 52 OTUs. From soup “a” to soup “g”, the input genomic masses of each of the 52 OTUs were 61, 48.8, 39, 31.2, 25, 20 and 16 ng. The same mass of spike-in DNA was then added to each soup (green DNA molecule). Each of the seven soups was made in triplicate, and all 21 soups were PCR’d in triplicate following the Begum pipeline (Yang et al. 2021) to detect and remove false reads. **B**. Malaise-trap-sample protocol. Each bulk sample of arthropods was non-destructively DNA-extracted by soaking in 5X volume of lysis buffer. From each of the 7 samples, 500 µl, 350 µl, 245 µl, 171.5 µl, 120 µl, and 84 µl lysis buffer was used to create 6 dilution soups, a fixed amount of spike-in DNA was added, and the mixture was co-purified. **(157 words**)

### 2.2 Preparation of Malaise-trap samples

244 Malaise-trap samples from 96 sites, using 99.9 % ethanol as the trapping liquid, were collected in and near a 384 km^2^ forested landscape containing the H.J. Andrews Experimental Forest (44.2° N, 122.2° W), Oregon, United States in July 2018 (Luo et al. 2022). Traps were left for 7 non-rainy days. To equalize biomass across individuals, we only kept the heads of large individuals (body lengths >2 cm) and then transferred the samples to fresh 99.9% ethanol to store at room temperature until extraction. The samples were air dried individually on filter papers for less than an hour and then transferred to 50 ml tubes or 5 ml tubes according to sample volume. The samples were then weighed. DNA was non-destructively extracted by soaking the samples in lysis buffer, using the protocol from Ji et al. (2020) and Nielsen et al. (2019). For this study, we selected seven samples spread over the study area, each of which is an independent test of our ability to recover the dilution gradient. After completion of lysis, we serially diluted the 7 samples by using 0.7X lysis buffer volume (500 µl, 350 µl, 245 µl, 171.5 µl, 120 µl and 84 µl) to create six soups per sample (ntot = 42). We used QIAquick PCR purification kit (Qiagen GmbH, Germany) following the manufacturer instructions to purify lysis buffer on one spin column per soup (Figure 3 B). We used a shallower gradient (0.7X) because our starting DNA amount was lower than with the mock soups.

### 2.3 Adding spike-in DNA

#### 2.3.1 Spike-in DNA

For our spike-ins, we used three insect species from China (Lepidoptera:Bombycidae, Coleoptera:Elateridae, Coleoptera:Mordellidae), none of which is expected to appear in the Oregon Malaise-trap samples. An alternative is to use one or more synthetic, random DNA sequences (Tkacz et al. 2018). Each of our three spike-ins is represented by a 658-bp COI fragment (Table S1) with primer binding sites that match the Folmer primers HCO2198 and LCO1490. For long-term storage, we inserted the COI fragments as plasmids into monoclonal bacteria. Plasmids were extracted using TIANprep Mini Plasmid Kit (Beijing, China) following manufacturer’s instructions.

#### 2.3.2 Adding spike-in to the mock soups

Adding too much spike-in wastes sequencing data, while adding too little risks loss of abundance information in at least some samples when the number of spike-in reads is too low to use as a reliable correction factor. Thus, we quantified the COI copy numbers of the mock soups and the spike-in DNA by qPCR (Table S2, Figure S1) and chose a volume so that spike-in reads should make up 1% of the total number of COI copies in the lowest-concentration mock soups, balancing efficiency with reliability. We used all three spike-in species here and mixed them (Bombycidae:Elateridae:Mordellidae) in a ratio of 1:2:4, which was added directly to the mock soups’ DNA since they were already purified.

#### 2.3.3 Adding spike-in to the Malaise-trap samples

From the 244 Malaise-trap samples, we first extracted 17 Malaise-trap samples without adding spike-ins, and then we used qPCR to quantify the mean COI concentrations of these 17 samples in order to decide how much spike-in to add. Before adding the spike-ins, we discovered that the Bombycid DNA spike-in had degraded, and so we used only two spike-in species for the Malaise trap samples, at a ratio of 1:9 (Mordellidae:Elateridae). We then chose 7 other samples for this study. In these samples, lysis buffer (500 µl, 350 µl, 245 µl, 171.5 µl, 120 µl, 84 µl) from each sample was transferred into clean 1.5 ml tubes, and the spike-in DNA was added. We then purified the DNA with the Qiagen QIAquick PCR purification kit, following the manufacturer instructions. DNA was eluted with 200 µl of elution buffer. In this way, the spike-in DNA was co-purified, co-amplified, and co-sequenced along with the sample DNA (Figure 3 B). We also recorded the total lysis buffer volume of each sample, for downstream correction.

### 2.4 Primer design

For this study, we simultaneously tested two methods for extracting abundance information: spike-ins and UMIs (Unique Molecular Identifiers). UMI tagging requires a two-step PCR procedure (Lundberg et al. 2013; Hoshino and Inagaki 2017), first using tagging primers and then using amplification primers (Figure S2). The tagging primers include (1) the Leray-FolDegenRev primer pair to amplify the 313-bp COI amplicon of interest, (2) a 1- or 2-nucleotide heterogeneity spacer on both the forward and reverse primers to increase sequence entropy for the Illumina sequencer, (3) the same 6-nucleotide sequence on both the forward and reverse primers to ‘twin-tag’ the samples for downstream demultiplexing, (4) a 5N random sequence on the forward primer and a 4N random sequence on the reverse primer (9N total) as the UMI tags, (5) and parts of the Illumina universal adapter sequences to anneal to the 3’ ends of the forward and reverse primers for the second PCR. By splitting the 9N UMI into 5N + 4N over the forward and reverse primers, we avoid primer dimers. The amplification primers include (1) an index sequence on the forward primer pair for Illumina library demultiplexing, and (2) the full length of the Illumina adapter sequences. For further explanation of the design of the tagging primers (except for the UMI sequences), see Yang et al. (2021).

### 2.5 PCR and the Begum pipeline

The first PCR amplifies COI and concatenates sample tags and UMIs and runs for only two cycles using KAPA 2G Robust HS PCR Kit (Basel, Roche KAPA Biosystems). We used the mlCOIintF-FolDegenRev primer pair (Yu et al. 2012, p. 2012; Leray et al. 2013), which amplifies a 313-bp fragment of the COI barcode; and we followed the Begum protocol (Zepeda-Mendoza et al. 2016; Yang et al. 2021), which is a wet-lab and bioinformatic pipeline that combines multiple independent PCR replicates per sample, twin-tagging and false positive controls to remove tag-jumping and reduce erroneous sequences. Twin-tagging means using the same tag sequence on both the forward and reverse primers in a PCR, and we use this design because during library index PCR for Illumina sequencing, occasional incomplete extensions can create new primers that already contain the tag of one amplicon, resulting in chimeric sequences with tags from two different amplicons (Schnell et al. 2015). Tag jumps thus almost always result in non-matching tag sequences, and these are identified and removed in the Begum pipeline. We performed 3 PCR replicates per sample, which means we used 3 different twin-tags to distinguish the 3 independent PCR replicates. Begum removes erroneous sequences by filtering out the reads that appear in a low number of PCR replicates (e.g. only one PCR) at a low number of copies per PCR (e.g. only 2 copies), because true sequences are more likely to appear in multiple PCRs with higher copy numbers per PCR. The 20 µl reaction mix included 4 µl Enhancer, 4 µl Buffer A, 0.4 µl dNTP (10 mM), 0.8 µl per primer (10 mM), 0.08 µl KAPA 2G HotStart DNA polymerase (Basel, Roche KAPA Biosystems), 5 µl template DNA and 5 µl water. PCR conditions were initial denaturation at 95°C for 3 minutes, followed by two cycles of denaturation at 95°C for 1 minute, annealing at 50°C for 90 seconds, and extension at 72°C for 2 minutes. Then the products were purified with 14 µl of KAPA pure beads (Roche KAPA Biosystems, Switzerland) to remove the primers and PCR reagents and were eluted into 16 µl of water.

The second PCR amplifies the tagged templates for building the libraries that can be sequenced directly on Illumina platform. The 50 µl reaction mix included 5 µl TAKARA buffer, 4 µl dNTP (10 mM), 1.2 µl per primer (10 mM), 0.25 µl TAKARA Taq DNA polymerase, 15 µl DNA product from the first PCR, and 23.35 µl water. PCR conditions were initial denaturation at 95°C for 3 minutes, 5 cycles of denaturation at 95°C for 30 seconds, annealing at 59°C for 30 seconds (-1 °C per cycle), extension at 72°C for 30 seconds, followed by 25 cycles of denaturation at 95°C for 30 seconds, annealing at 55°C for 30 seconds, extension at 72°C for 30 seconds; a final extension at 72°C for 5 minutes, and cool down to 4°C.

From all second PCR products, 2 µl was roughly quantified on 2% agarose gel with Image Lab 2.0 (Bio-Rad, USA). For each set of PCR reactions with the same index, amplicons were mixed at equimolar ratios to make a pooled library. One PCR negative control were set for each library. We sent our samples to Novogene (Tianjin, China) to do PE250 sequencing on Illumina NovaSeq 6000, requiring a 0.8 GB raw data from each PCR reaction.

### 2.6 Bioinformatic processing

AdapterRemoval 2.1.7 was used to remove any remaining adapters from the raw data (Schubert et al., 2016). Sickle 1.33 was used to trim away low-quality bases at the 3’ends. BFC V181 was used to denoise the reads (Li, 2015). Read merging was performed using Pandaseq 2.11 (Masella et al. 2012). Begum was used to demultiplex the reads by sample tag and to filter out erroneous reads (https://github.com/shyamsg/Begum, accessed 07 Sep 2021). We allowed 2-bp primer mismatches to the twin-tags while demultiplexing, and we filtered at a stringency of accepting only reads that appeared in at least two PCRs at a minimum copy number of 4 reads per PCR, with minimum length of 300 bp. This stringency minimized the false positive reads in the negative PCR control.

For mock-soup data, we need to compare the UMI and read numbers in each PCR set. However, Begum cannot recognize UMIs. Also because of our complicated primer structure, there is no software available for our data to count the UMIs per OTU in each PCR set. Thus, we wrote a custom bash script to process the mock-soup data from the Pandaseq output files, which include all the UMIs, tags, and primers. First, we used *Begum*-filtered sequences as a reference to filter reads for each PCR set and put the UMI information on read headers. Then we carried out reference-based OTU clustering for each PCR set with QIIME 1.9.1 (`pick_otus.py -m uclust_ref -s 0.99`) (Caporaso et al. 2010; Edgar 2010), using the OTU representative sequences from barcoding Sanger sequencing as the reference, counted UMIs and reads for each OTU in each PCR set, and generated two OTU tables, separately with UMI and read numbers.

For the Malaise-trap data, we directly used the Begum pipeline. After Begum filtering, vsearch 2.14.1 `(--uchime_denovo`) (Rognes et al. 2016) was used to remove chimeras. Sumaclust 1.0.2 was used to cluster the sequences of Malaise-trap samples into 97% similarity OTUs. The python script tabulateSumaclust.py from the DAMe toolkit was used to generate the OTU table. Finally, we applied the R package LULU 0.1.0 with default parameters to merge oversplit OTUs (Frøslev et al. 2017). The OTU table and OTU representative sequences were used for downstream analysis.

### 2.7 Statistical analyses

All statistical analyses were carried out in *R* 4.1.0 (R Core Team 2021), and we used the {lme4} 1.1-27 package (Bates et al. 2015) to fit linear mixed-effects models, using OTU, soup replicate, and PCR replicates as random factors, to isolate the variance explained by the sole (fixed-effect) predictor of interest: OTU size. Model syntax is given in the legend of Figure 4. We used the {MuMIn} 1.43.17 package (CRAN.R-project.org/package=MuMIn, accessed 2 Jan 2022) to calculate the variance explained by fixed effects only (marginal R^2^). To carry out spike-in correction, we first calculated a weighted mean from the added spike-ins (e.g. mean(Bombycidae + Elateridae/2 + Mordellidae/4)), rescaled the new mean spike-in so that the smallest value is equal to 1, and divided each row’s OTU size and UMI number by the weighted, scaled spike-in.

**Figure 4.**
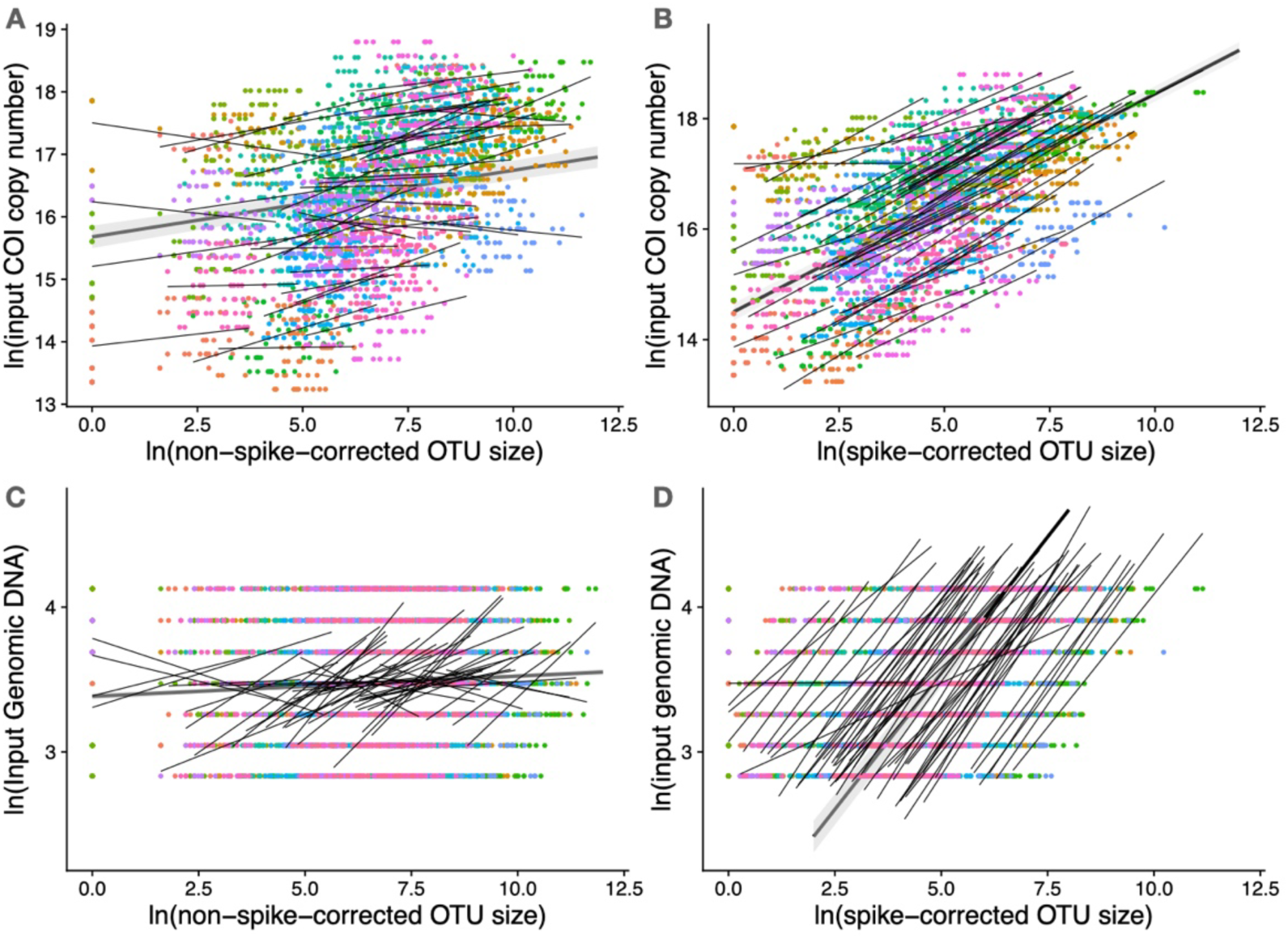
Recovery of within-species abundance change in COI copy number and in genomic DNA concentration in the mock-soup experiment. For visualisation, all data points are shown (including all soup and PCR replicates), each thin line is fit to one of the OTUs across the seven serially diluted mock-soup samples, and the thick line represents the fitted model in which OTUs were treated as a random factor. **A**. Non-spike-corrected OTU size (number of reads per OTU per soup) poorly predicts within-species variation in input COI copy number (linear mixed-effects model, marginal R^2^ = 0.04, conditional R^2^ = 0.85). **B**. Spike-corrected OTU size successfully predicts within-species variation in input COI copy number (mixed-effects linear model, marginal R^2^ = 0.42, conditional R^2^ = 0.96), but species bias remains, as can be seen in the orders-of-magnitude variation in intercepts. **C**. Non-spike-corrected read number poorly predicts within-species variation in input genomic DNA concentration (linear mixed-effects model, marginal R^2^ = 0.01, conditional R^2^ = 0.01). **D**. Spike-corrected read number successfully predicts within-species variation in input genomic DNA concentration but more poorly for species represented by small OTUs (linear mixed- effects model, marginal R^2^ = 0.52, conditional R^2^ = 0.95) despite species bias (Figure 1). Model syntax: `lme4::lmer(log.input_gDNA or log.inputCOI_copynumber ∼ log.OTUsize + (log.OTUsize | OTUID) + (1 |soupRep/pcrRep)`) (Bates et al. 2015). Marginal R^2^ is variance explained by the fixed effect, and conditional R^2^ is variance explained by the whole model. (**226 words**)

## Results

### 3.1 Bioinformatic processing of the Malaise-trap samples and the mock soups

Five libraries yielded a total of 283,319,770 paired-end reads, of which 247,285,097 were merged successfully in Pandaseq. After *Begum* sorting and demultiplexing, which removed a large number of tag-jumped reads and some reads <300 bp length, we retained 106,649,397 reads. After *Begum*’s filtering of erroneous reads, we retained 76,289,802 reads, and after *de-novo* chimera removal, we retained 73,818,971 reads. Sequences were clustered at 97% similarity into 1,188 OTUs, and LULU combined the OTUs of the Malaise-trap samples into 435 OTUs. After removing the spike-in OTUs, the seven Malaise-trap samples contained a total of 432 OTUs. All 52 OTUs of the 7 mock soups were recovered.

### 3.2 Mock soups, COI copy number

Without spike-in correction, OTU size (numbers of reads per OTU) predicts almost none of the within-species (dilution-gradient-caused) variation in COI copy number (R^2^ = 0.04, all values marginal R^2^), but with spike-in correction, OTU size predicts 42.0% of the variation (Figure 4 AB). As expected, UMI number by itself does not predict input COI copy number (R^2^ = 0.05), but with spike-in correction, UMI number does predict COI copy number (R^2^ = 0.42) (Figure S3 AB). Also as expected, spike-in correction does not achieve across-species quantification, as shown by the orders of magnitude variation in intercepts across the 52 OTUs. Note that this experiment pooled DNA extracts with equalised concentrations of genomic DNA mass per species, which suggests that PCR bias is the main source of species bias in this dataset.

### 3.4 Mock soup within-species abundance in input genomic DNA mass

Of course, our goal is to estimate not COI copy number but specimen biomass. We thus tested how well OTU size and UMI numbers predicted genomic DNA concentration. Non-spike-corrected OTU size and UMI number both failed to predict input genomic DNA mass (R^2^ < 0.02 for both, Figure 4 C and Figure S3 C), but spike-corrected OTU size and UMI number again both successfully predicted input genomic DNA mass (R^2^ = 0.53 and 0.52, Figure 4 D and Figure S3 D).

### 3.5 Malaise-trap within-species abundance recovery

Recall that each of the 7 selected Malaise-trap samples was serially diluted by 0.7X to create six soups per sample. Non-spike-corrected OTU size did not predict within-species variation in input genomic-DNA mass (p = 0.33) (Figure 5 A), but spike-corrected OTU size again did predict within-species variation in input genomic-DNA mass (R^2^ = 0.53) (Figure 5 B).

**Figure 5.**
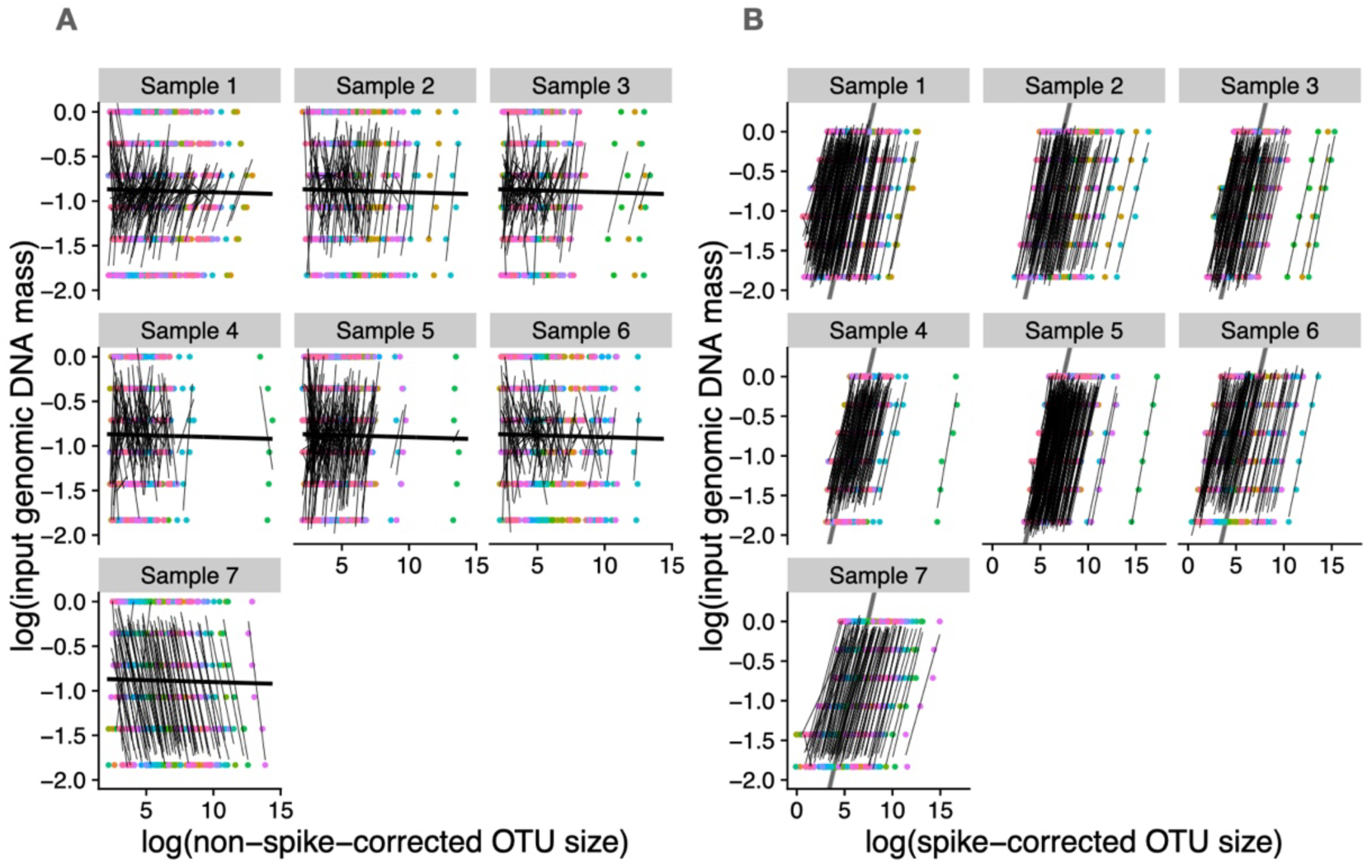
Prediction of within-species variation in genomic DNA concentration in the Malaise-trap samples. For visualisation, each thin line is fit to an OTU’s serial dilution made from each of the seven Malaise-trap samples, and the thick lines are the fitted model with sample and OTU as random factors. There are 176, 113, 111, 104, 196, 110, and 82 OTUs in samples 1-7, respectively. **A**. Non-spike-corrected OTU size (read number per OTU and sample) does not predict within-species variation in genomic DNA concentration (marginal R^2^ = 0.0, conditional R^2^ = 0.0). **B**. Spike-corrected OTU size successfully predicts within-species variation in genomic DNA concentration (marginal R^2^ = 0.53, conditional R^2^ = 0.98) despite species bias, represented by the different intercepts. A similar protocol was followed in Ji et al. (2020), where it was called “FSL correction”. **Full model syntax**: `lme4::lmer(log.input_gDNA ∼ log.OTUsize + (1 | sample/OTUID)`). (**145 words**)

## Discussion

We propose that there is a useful distinction to be made between *within*-species and *across*-species abundance information (Figures 1, 2). Within-species abundance information can be enough to improve the inference of species interactions, the modelling of population dynamics and species distributions, the biomonitoring of environmental state and change, and the inference of false positives and negatives (Carraro et al. 2020, 2021; Abrego et al. 2021; Rojahn et al. 2021, and Figure 2). We thus recommend that future quantitative eDNA studies should make clear which abundance measure is being estimated.

We experimentally show that spike-ins allow the recovery of within-species abundance change, by removing pipeline noise (Figures 4, 5), even given the equimolar pooling step before library prep. In both experiments, we used a multi-species spike-in. The potential benefit of multiple species is the option to detect experimental error, which could be exposed by the spike-ins deviating strongly from their input ratios (Ji et al. 2020), but the cost is usage of sequence data on spike-in reads. Ushio et al. (2018) have also shown that spike-ins recover within-species abundance change, and they moreover showed that a spike-in can be used on trace fish eDNA in water samples. We note that Ushio et al.’s method is more complex than our method of counting the number of spike-in reads per sample, and so the optimal method for trace DNA remains an open research question.

In our first test, we serially diluted 52 OTUs into seven mock soups, and after spike-in correction (Figure 3), we were able to recover within-species abundance change in both input COI copy number and input genomic DNA (Figure 4), the latter of which should be more closely correlated with organism biomass. In our second test, we serially diluted each of the seven Malaise-trap soups into six soups (Figure 3), and we were able to recover within-species abundance change in input genomic DNA (Figure 5).

Finally, our experimental protocol included Unique Molecular Identifiers (UMIs), and we find that they can also recover within-species abundance change (Figure S3), but UMIs require a laborious two-step PCR protocol for no additional quantification benefit over the spike-in (Figure S3). On the other hand, UMIs have other advantages that could recommend them over a physical spike-in, such as not taking up sequencing data, which could make them more suitable for trace DNA sample types, contamination detection, and error correction. Contaminant and erroneous sequences should be present at low abundances and thus capture few UMIs (Fields et al. 2021).

Additional alternatives to external spike-ins include a method introduced by Lundberg et al. (2021), who describe a two-step PCR method to use a single-copy host gene as a built-in spike-in. Also, in the Supplementary Information code for Figure 4 (S4), we apply the model-based pipeline-noise estimator to the mock-soup dataset and achieve an R^2^=11.8% for prediction of COI copy number, which lies between the R^2^ values achieved for the non-physical-spike-corrected (R^2^=0.04) and physical-spike-corrected values (R^2^=0.42) (Figure 4 B). We also achieve an R^2^=21.3% for prediction of genomic DNA, again intermediate between the non- physical-spike-corrected (R^2^=0.0) and physical-spike-corrected values (R^2^=0.53) (Figure 4 D). In the Malaise-trap data (Supplementary Information S5), the model-based approach performed poorly at recovering genomic concentration. The issue was that samples had been pooled to equimolar concentration, which led to strong confounding of pipeline noise and differences in total abundance across samples. The model-based approach did however correctly infer that there were no compositional effects in this dataset, consistent with a dilution gradient. This behaviour is as expected for the model-based method – it will recover relative not absolute DNA concentrations, hence is a tool best used to study effects on compositional not total abundance.

Statistical analysis of DNA-based datasets will also need to exploit better within-species abundance information. The most straightforward method is to incorporate spike-in counts as an offset term in general linear models. For species distribution modelling, there is a need for software packages to utilise abundance data that ranges continuously from [0,1], whereas to our knowledge, practitioners can effectively now only choose between presence/absence and absolute-abundance data.

We conclude with the acknowledgment that relative species abundance remains the more difficult abundance-estimation problem, given the many hidden sources of species bias along metabarcoding and metagenomic pipelines (McLaren et al. 2019), but promising solutions are now starting to be available for amplicon (Silverman et al. 2021; Williamson et al. 2021; Shelton et al. 2022a) and metagenomic datasets (Lang et al. 2019; Peel et al. 2019). Note that even if species biases can be corrected by using one of these techniques, it is still necessary to use a spike-in to correct for pipeline noise.

## Acknowledgments

We thank Sarah Bourlat, Nathan Geraldi, Lucie Zinger, and three anonymous reviewers for very helpful comments on the manuscript. The authors were supported by the Key Research Program of Frontier Sciences, CAS (QYZDY-SSW-SMC024), Supported by the Strategic Priority Research Program of Chinese Academy of Sciences, Grant No. XDA20050202, the University of East Anglia, the State Key Laboratory of Genetic Resources and Evolution (GREKF19-01, GREKF20-01, GREKF21-01) at the Kunming Institute of Zoology, the University of East Anglia, and the University of Chinese Academy of Sciences.

## Data Accessibility Statement

Data and R scripts for Figures 4, 5, S3, and the model-based estimator are available in Supplementary Information as RStudio projects (S4 and S5).

All sequence data, reference files, folder structure, output files and scripts (32.5 GB) are available at DataDryad’s designated temporary URL for review. The dataset will receive a DOI upon acceptance of the manuscript. (https://datadryad.org/stash/share/0rJ5Yy2PRIv5UpVrCS95Wf7pY0J2R_Hqic6DWyMe <u>aD8</u>).

Most of the sequence data are experimental mixtures and thus have no value outside of this study. The sequence data for the Malaise-trap experiment (n=7) will be submitted to SRA with the rest of the Malaise-trap dataset (ntot=121) with the publication analysing that dataset.

## Benefit Statement

There are no benefits to report. These samples were collected in Kunming, Yunnan, China (mock soup experiments) and HJ Andrews Experimental Forest, Oregon, China (Malaise-trap experiment).

## Supplementary Information: Unabridged Methods

### 1 Mock soup construction

#### 1.1 Input species

We used Malaise traps to collect 286 arthropods in Kunming, China (25°8’23” N, 102°44’17” E). DNA was extracted from each individual using the DNeasy Blood & Tissue Kit (Qiagen GmbH, Germany). Mean genomic DNA per species concentration was quantified from three replicates using PicoGreen fluorescent dye. We DNA-barcoded the individuals using the Folmer primer pair LCO1490 5’- GGTCAACAAATCATAAAGATATTGG -3’, and HC02198 (5’- TAAACTTCAGGGTGACCAAAAAATCA -3’) (Folmer et.al 1994), with PCR parameters of initial denaturation at 95°C for 3 minutes, followed by 34 cycles of denaturation at 94°C for 30 seconds, annealing at 50°C for 30 seconds and extension at 72°C for 1 minute, a final extension at 72° C for 5 minutes, and a cooldown to 4°C. After the 658- bp COI sequences were cut to 313 bp based on our metabarcoding primers (see *4 Primer design*), 286 arthropods were clustered into 168 OTUs at 97% similarity. To create mock soups, we selected 52 individuals with genomic DNA >20 ng/µl, representing 52 OTUs.

#### 1.2 OTU quantification

We quantified COI concentrations of the 52 OTUs using the mean of three qPCRs with the *Leray-FolDegenRev* primer pair (Yu et al. 2012; Leray et al. 2013). First, to create a standard DNA curve, we used a purified and sequenced COI amplicon (Lepidoptera Bombycidae *Bombyx mori*) and quantified its concentration with a Qubit, from which we calculated COI copy number from the COI concentration (ng/µl), molar mass (233530.15 g/mol), and the Avogadro constant.

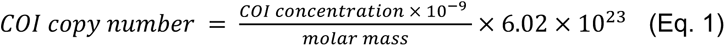

This was 0.5X serially diluted into six concentration levels, as follows:

1. Start by diluting the above PCR amplicon to 100x by transferring 5 µl of amplicon to 495 µl of Ultragrade water. Vortex for 20 sec. Label as Tube 1.
2. Label 5 tubes from 2 to 6 and add 250 µl of water into each tube.
3. Transfer 250 µl from Tube 1 to Tube 2, and vortex for 20 sec.
4. Transfer 250 µl from Tube 2 to Tube 3, and vortex for 20 sec.
5. Continue the dilution series to Tube 6.
6. Use Tubes 1-6 as the standard DNA gradient for qPCR.

For each of the 52 OTUs, we used qPCR to quantify COI concentration (copy/µl). qPCRs were performed in a total volume of 20 µl with 10 µl of TAKARA SYBR premix Ex Taq (TaKaRa Biosystems, Dalian, China), 0.4 µl of Rox, 0.8 µl each of 10 µM of Leray and FolDegenRev primers (synthesized by Invitrogen, Shanghai, China), 2 µl of template DNA, and 6 µl of PCR-grade water. The qPCR conditions were 30 sec of initial denaturation at 95°C, followed by 35 cycles of denaturation at 95°C for 10 sec, 30 sec of annealing at 50°C, and 30 sec of extension at 72°C. Each OTU was quantified twice.

**Figure S1.**
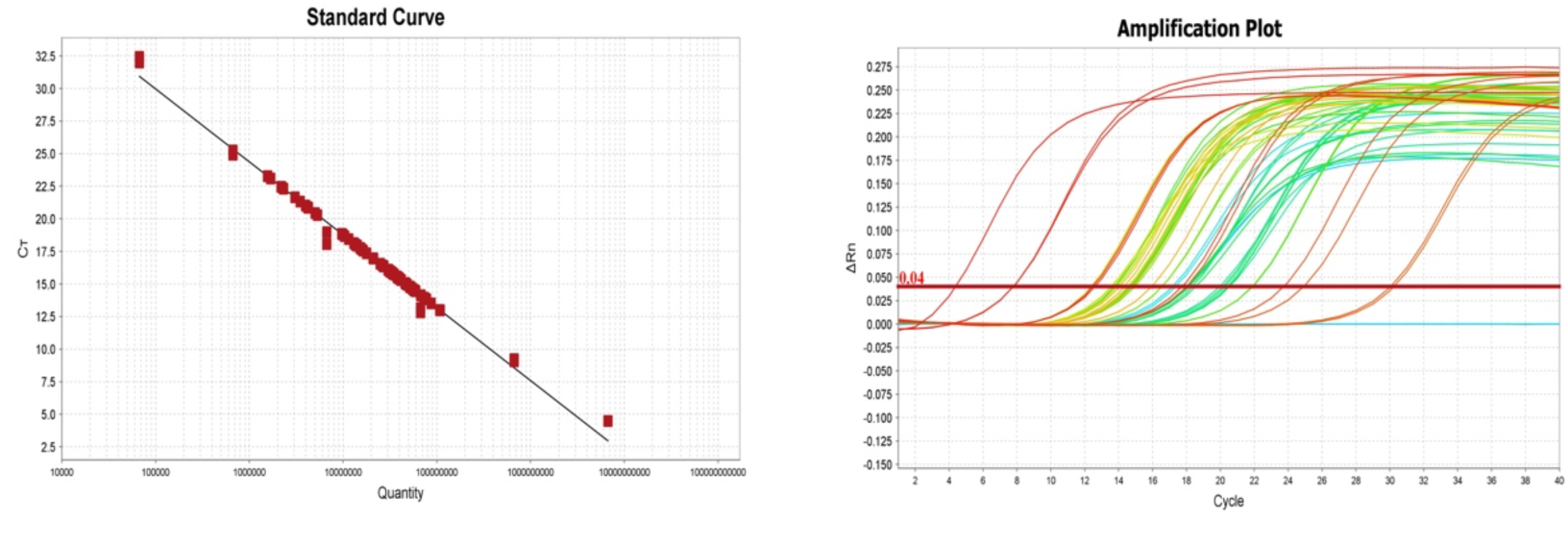
Left. qPCR standard curve (CT ∼ input-COI concentration). Right. qPCR amplification curves, with the 12 (= 2 replicates x 6 concentrations) red lines representing the standard-DNA curves. The other colours represent the 52 OTUs, each amplified twice. All OTU curves lie within the bounds of the standard-DNA curves, allowing interpolation of absolute COI concentrations.

#### 1.3 Creation of mock soups

Each mock soup was constructed with equal masses of purified genomic DNA from 52 OTUs. Input genomic DNA masses of each OTU from soup 1 to soup 7 decreased by 0.8X to create seven mock soups of differing absolute abundances. The individual-OTU genomic DNA masses for the seven mock soups were 61, 48.8, 39, 31.2, 25, 20, and 16 ng and were independently added to the mock soups. Each of the seven soups was made in triplicate (ntot = 21).

### 2 Preparation of Malaise-trap samples

#### 2.1 Malaise trap sampling

244 Malaise-trap samples from 96 sites, using 99.9% ethanol as the trapping liquid, were collected in and near a 384 km^2^ forested landscape containing the H.J. Andrews Experimental Forest (44.2° N, 122.2° W), Oregon, United States in July 2018. Traps were left for 7 non-rainy days. To equalize biomass across individuals, we only kept the heads of large individuals (body lengths >2 cm) and then transferred the samples to fresh 99.9% ethanol to store at room temperature until extraction.

#### 2.2 DNA extraction

The samples were air dried individually on filter papers for less than an hour and then transferred to 50 ml tubes or 5 ml tubes according to sample volume. The samples were then weighed. DNA was non-destructively extracted by soaking the samples in lysis buffer, using the protocol from Ji et al. (2020) and Nielsen et al. (2019). In short, we added lysis buffer to each sample at a 5 ml:1 mg volume:air- dried-mass ratio and incubated the samples in an incubator at 56°C, shaking for 60 hrs. The recipe to make 50 ml of lysis buffer is 2 ml of 1M Tris-HCl buffer (PH 8.0), 1 ml of 5M sodium chloride, 10 ml of 10% Sodium Dodecyl Sulfate, 150 µl of 1M calcium chloride, 34.225 ml of PCR-grade water, 2 ml of 1M Dithiothreitol, and 625 µl of Proteinase K solution (20 mg/ml). After extraction, visual checks confirmed that the extracted insects largely retained their body shapes, although they became darker. Note that non-destructive DNA extraction increases the effects of species biases since species with hard exoskeletons likely release less tissue.

For this study, we selected seven samples spread over the study area. After completion of lysis, we serially diluted the 7 samples by using 0.7X lysis buffer volume (500 µl, 350 µl, 245 µl, 171.5 µl, 120 µl and 84 µl) to create six soups per sample (ntot = 42). We used QIAquick PCR purification kit (Qiagen GmbH, Germany) by following the manufacturer instructions to purify lysis buffer on one spin column per soup.

### 3 Adding spike-in DNA

#### 3.1 Spike-in DNA

For our spike-ins, we used three insect species from China, none of which is expected to appear in the Oregon Malaise-trap samples. (An alternative to our protocol is to use one or more synthetic, random DNA sequences (Tkacz et al. 2018)). Each of our three spike-ins is represented by a 658-bp COI fragment (Tables S1, S2) with primer binding sites that match the Folmer primers HCO2198 and LCO1490. For long-term storage, we inserted the COI fragments as plasmids into monoclonal bacteria. Plasmids were extracted using TIANprep Mini Plasmid Kit (Beijing, China) following manufacturer’s instructions.

qPCR was used to quantify the three spike-in DNA COI concentrations (ng/µl) using TAKARA SYBR Premix Ex Taq (TaKaRa Bio, Japan). COI copy numbers of the three spike-in DNA were calculated using equation 1.

In our plan, we will mix the three spike-ins in a ratio of 1:2:4 (Bombycidae:Elateridae:Mordellidae), to check for error and degradation; the reads from the three spike-ins should be found in the ratio of 1:2:4, but if, for instance, one of the spike-ins is not at the correct ratio with the other two, we can omit that species’ reads and recalculate the spike-in correction.

**Table S1.**
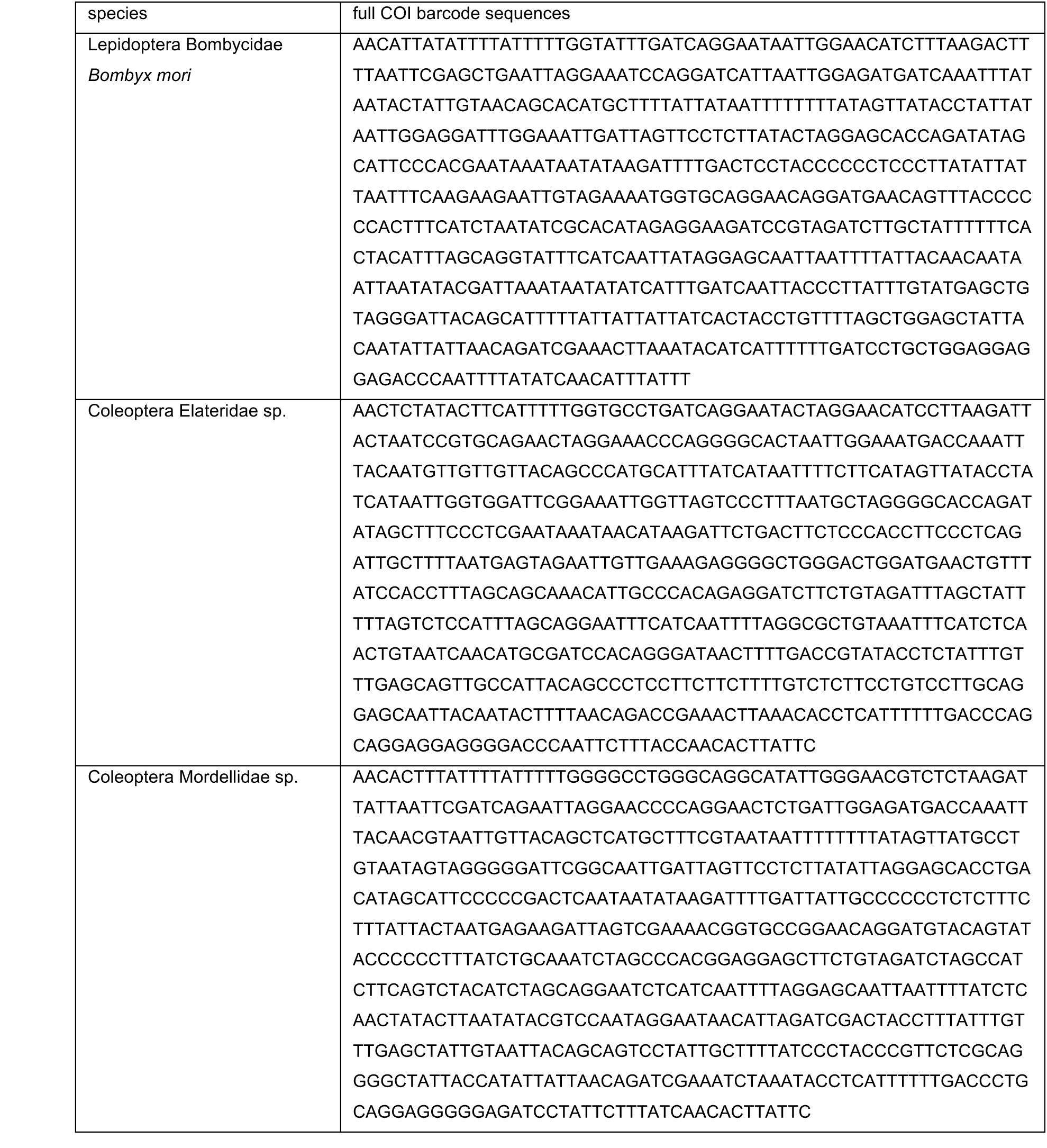
DNA-barcode sequences of the three spike-in species.

**Table S2.** COI concentrations of the three spike-in species

| species | COI concentration (ng/μl) | COI copy number (copy/μl) |
| --- | --- | --- |
| Lepidoptera Bombycidae<br>*Bombyx mori* | 0.0002 | $5.17 \times 10^5$ |
| Coleoptera Elateridae sp. | 0.3914 | $1.02 \times 10^9$ |
| Coleoptera Mordellidae sp. | 0.05807 | $1.51 \times 10^8$ |

#### 3.2 Adding spike-in to the mock soups

Adding too much spike-in wastes sequencing data, while adding too little spike-in risks the loss of abundance information because in some samples, the number of spike-in reads will be too low to use as a reliable correction factor. Thus, we quantified the COI copy numbers of the mock soups by qPCR and chose a volume so that spike-in reads should make up 1% of the total number of COI copies in the lowest-concentration mock soups, balancing efficiency with reliability.

Because we mixed 52 individual OTUs to create the mock soups, we can sum the input COI copy numbers of all 52 OTUs to calculate the COI copy number of the lowest- concentration mock soup (5.25 * 10^8^ copies). We used 1% of this number to calculate the mass of total spike-in to add to all 21 soups (5.25 * 10^6^ copies). To mix spike-ins DNA conveniently, the Elateridae spike-in was diluted 1000-fold to 1.02 * 10^6^ copies and the Mordellidae spike-in was diluted 100-fold to 1.51 * 10^6^ copies before mixing.

Then we added 1.45 µl of *Bombyx* spike-in (1.45 * 5.17 *10^5^ = 0.75 * 10^6^), 1.5 µl of diluted Elateridae (1.5 * 1.02 * 10^6^ = 1.5 * 10^6^) and 2 µl of diluted Mordellidae (2 * 1.51 * 10^6^ = 3 * 10^6^) to each soup.

The sum of the three spike-ins was thus: (0.75+1.5+3)× 10^6^ = 5.25× 10^6^, and their ratio was: 0.75 × 10^6^ :1.5 ×10^6^ :3 × 10^6^=1:2:4.

Spike-in DNA was added directly to the mock soups’ DNA since they were already purified.

#### 3.3 Adding spike-in to the Malaise-trap samples

From the 244 Malaise-trap samples, we first extracted 17 Malaise-trap samples without adding spike-ins, and then in order to decide how much spike-in should be added, we used qPCR to quantify the COI concentrations of these 17 Malaise-trap samples, using the same protocol that we used for the mock-soup OTUs. The mean COI concentration per sample was 0.006 ng/µl, for each of which we added 200 µl volume. Mean COI mass per aliquot was thus 1.2 ng (0.006 ng/µl x 200 µl), and 1% is 0.012 ng.

Because the Bombycid DNA spike-in had degraded after we arrived in University of Oregon where we extracted DNA from the Malaise-trap samples, we used only the other two spike-in species for the Malaise trap samples, at a ratio of 1:9 (Mordellidae:Elateridae). Mordellidae and Elateridae were re-quantified by qPCR and were diluted 100-fold. We added 2.7 µl of diluted Elateridae (2.7 µl * 0.004 ng/µl = 0.0108 ng) and 2 µl of diluted Mordellidae (2 µl * 0.0006 ng/µl = 0.0012 ng) to make spike-ins’ mixture for each Malaise-trap sample.

The sum of the two spike-ins: 0.0012 ng + 0.0108 ng = 0.012 ng, and their ratio was: 0.0012:0.0108 = 1:9.

We used the same spike-in mass for all the Malaise-trap samples except for the 17 samples that had already been extracted. For the selected 7 samples used in this study, lysis buffer (500 µl, 350 µl, 245 µl, 171.5 µl, 120 µl and 84 µl) from each sample was transferred into a clean 1.5 ml tube, and the spike-in DNA was added and vortexed for 10 sec. We then extracted DNA with the Qiagen QIAquick PCR purification kit, following the manufacturer instructions. DNA was eluted with 200 µl of elution buffer. In this way, the spike-in DNA was co-purified, co-amplified, and co-sequenced along with the sample DNA (Figure 5). We also recorded the total lysis buffer volume of each sample, for downstream correction.

### 4 Primer design

For this study, we simultaneously tested two methods for extracting abundance information: spike-ins and UMIs (Unique Molecular Identifiers). UMI tagging requires a two-step PCR procedure, first using tagging primers and then using amplification primers (Figure S2). The tagging primers include (1) the Leray-FolDegenRev primer pair to amplify the 313-bp COI amplicon of interest, (2) a 1- or 2-nucleotide heterogeneity spacer on both the forward and reverse primers to increase sequence entropy for the Illumina sequencer, (3) the same 6-nucleotide sequence on both the forward and reverse primers to ‘twin-tag’ the samples for downstream demultiplexing, (4) a 5N random sequence on the forward primer and a 4N random sequence on the reverse primer (9N total) to act as the UMI tags for across-species abundance estimation, (5) and parts of Illumina universal adapter sequences to anneal to the 3’ ends of the forward and reverse primer regions for the second PCR. By splitting the 9N UMI into 5N + 4N over the forward and reverse primers, we avoid primer dimers. The amplification primers include (1) the annealing primer pair to bind to the amplicons of the first PCR, (2) the full length of the Illumina adapter sequences. For further explanation of the design of the tagging primers (except for the UMI sequences), see Yang et al. (2021).

**Figure S2.**
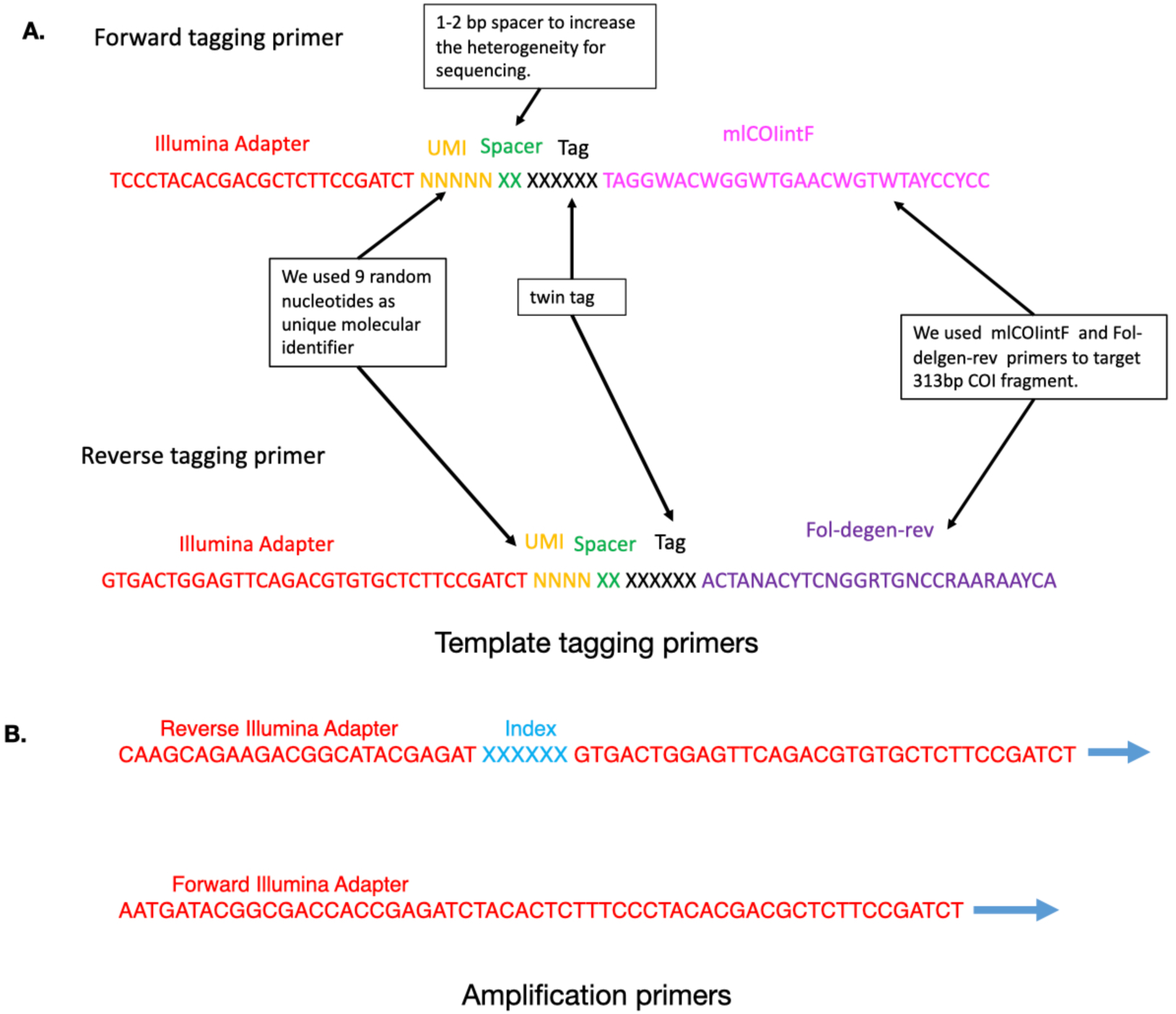
Two-step PCR protocol. A. In the first step, we amplified a 313bp COI amplicon using the Leray-FolDegenRev primers. To Both the forward and reverse primers, we added a 1-2 nucleotide heterogeneity spacer, the same 6-nucleotide sample tag (twin tags), a 9-random-nucleotide UMI (unique molecular identifier, split as 5 Ns on the forward and 4 Ns on the reverse), and a part of Illumina adapter sequence to anneal to the PCR primers in the second step. B. In the second step, we concatenate a library index sequence and the forward and reverse Illumina adapter sequences. Additional details on primer design (other than the UMI portion) can be found in Yang et al. (2021)

### 5 PCR and the Begum pipeline

We performed a two-step PCR (Lundberg et al. 2013).

The first PCR amplifies COI and concatenates sample tags and UMIs and runs for only two cycles using KAPA 2G Robust HS PCR Kit (Basel, Roche KAPA Biosystems). We used the mlCOIintF-FolDegenRev primer pair (Yu et al. 2012; Leray et al. 2013), which amplifies a 313-bp fragment of the COI barcode; and we followed the *Begum* protocol (Zepeda-Mendoza et al. 2018, Yang et al. 2021), which is a wet-lab and bioinformatic pipeline that combines multiple independent PCR replicates per sample, twin-tagging and false positive controls to remove tag-jumping and reduce erroneous sequences. Twin-tagging means using the same tag sequence on both the forward and reverse primer in a PCR. We performed 3 PCR replicates per sample, which means we used 3 different twin-tags to distinguish the 3 independent PCR replicates. *Begum* removes erroneous sequences by filtering out the sequences that appear in a low number of PCR replicates (e.g. one) at a low number of copies per PCR (e.g. 4 copies), because true sequences are more likely to appear in multiple PCRs and with high copy numbers. The 20 µl reaction mix included 4 µl Enhancer, 4 µl Buffer A, 0.4 µl dNTP (10 mM), 0.8 µl of 10 mM forward primer, 0.8 µl of 10 mM reverse primer, 0.08 µl KAPA 2G HotStart DNA polymerase (Basel, Roche KAPA Biosystems), 5 µl template DNA and 5 µl water. PCR conditions were initial denaturation at 95°C for 3 minutes, followed by two cycles of denaturation at 95°C for 1 minute, annealing at 50°C for 90 seconds, and extension at 72°C for 2 minutes. Then the products were purified with 14 µl of KAPA pure beads (Roche KAPA Biosystems, Switzerland) to remove the primers and PCR reagents and were eluted into 16 µl of water.

The second PCR amplifies the tagged templates for building the libraries which can be sequenced directly on Illumina platform. The 50 µl reaction mix included 5 µl TAKARA buffer, 4 µl dNTP (10 mM), 1.2 µl of 10mM forward primer, 1.2 µl of 10mM reverse primer, 0.25 µl TAKARA Taq DNA polymerase, 15 µl DNA product from the first PCR, and 23.35 µl water. PCR conditions were initial denaturation at 95°C for 3 minutes, 5 cycles of denaturation at 95°C for 30 seconds, annealing at 59°C for 30 seconds (-1 °C per cycle), extension at 72°C for 30 seconds, followed by 25 cycles of denaturation at 95°C for 30 seconds, annealing at 55°C for 30 seconds, extension at 72°C for 30 seconds; a final extension at 72°C for 5 minutes, and cooldown to 4°C.

### 6 Bioinformatic processing

*AdapterRemoval* 2.1.7 was used to remove any remaining adapters from the raw data (Schubert et al. 2016). *Sickle* 1.33 was used to trim away low-quality bases at the 3’ends. *BFC* v181 was used to denoise the reads (Li 2015). Read merging was performed using *Pandaseq* 2.11 (Masella et al. 2012). *Begum* was used to demultiplex the reads by sample tag and to filter out erroneous reads (https://github.com/shyamsg/Begum, accessed 07 Sep 2021). We allowed 2-bp primer mismatches to the twin-tags while demultiplexing, and we filtered at a stringency of accepting only reads that appeared in at least two PCRs at a minimum copy number of 4 reads per PCR, with minimum length of 300 bp. This stringency minimized the false positive reads in the negative PCR control. *vsearch* 2.14.1 (Rognes et al. 2016) was used to remove chimeras `(--uchime_denovo`). *Sumaclust* 1.0.2 was used to cluster the sequences into 97% similarity OTUs. The python script *tabulateSumaclust.py* from the DAMe toolkit was used generate the OTU table. Finally, we applied the R package LULU 0.1.0 with default parameters to merge oversplit OTUs (Frøslev et al. 2017). We also removed any OTUs in which we found stop codons. The OTU table and OTU representative sequences were used for downstream analysis.

*Begum* removed UMIs while sorting tags. Because of our complicated primer structure, there is no software available for our data to count the number of UMIs per OTU. We wrote our own bash scripts to process data from the Pandaseq-merged files, which includes all the UMIs, tags, and primers. We first used *Begum*-filtered sequences as a reference to filter reads in each PCR set and put the UMI information on the read headers. Then we carried out reference-based OTU clustering for each PCR with QIIME 1.9.1 (`pick_otus.py -m uclust_ref -s 0.99`), using the OTU representative sequences as the reference, counted UMIs and reads for each OTU in each PCR set, and generated the UMI and READ OTU tables.

## UMI Results

**Figure S3.**
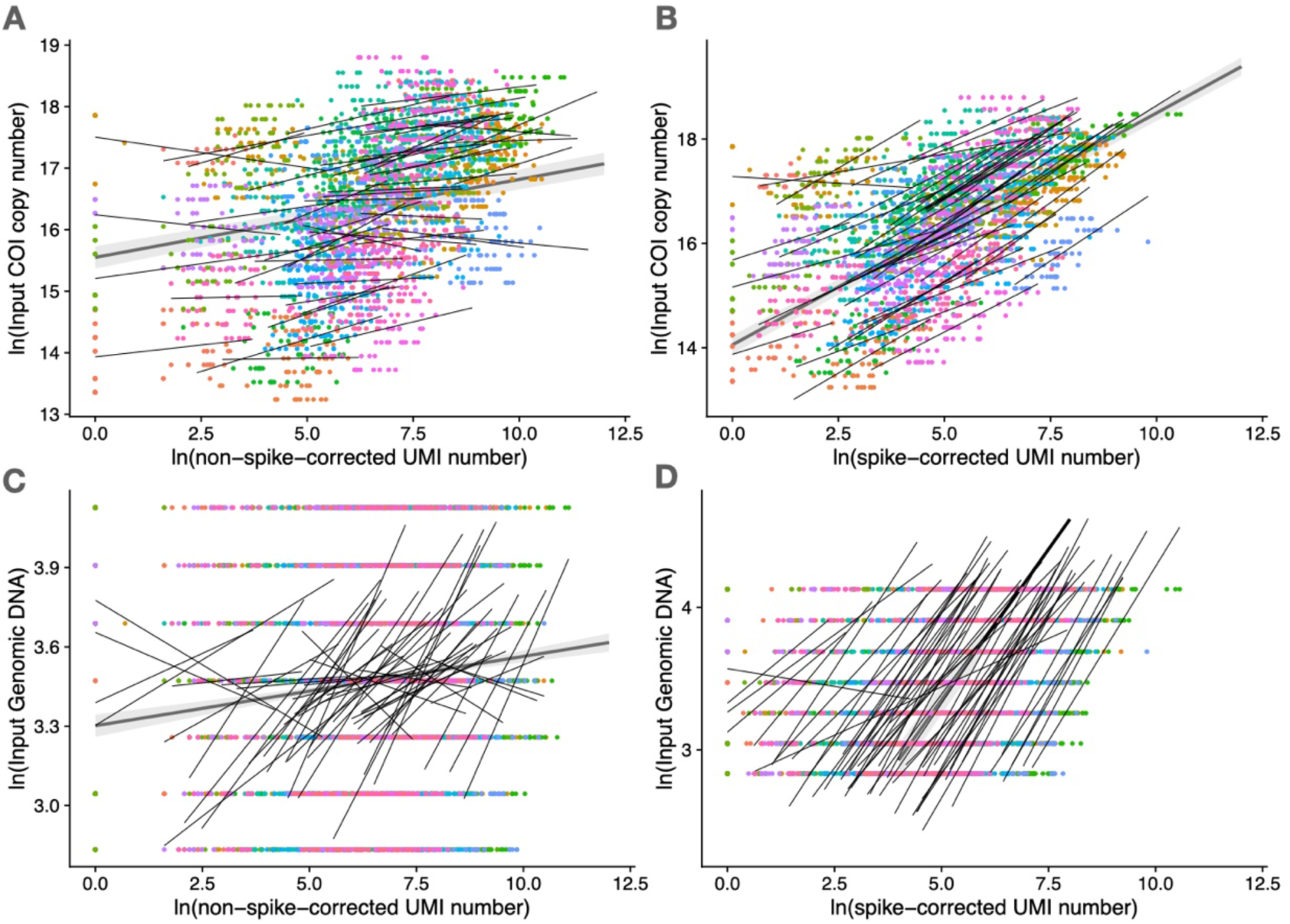
Recovery of mock soup within-species abundance change with UMI in COI copy number and in genomic DNA concentration. For visualisation, all data points are shown (including all soup and PCR replicates), each thin line is fit to one of the OTUs across the seven serially diluted mock-soup samples, and the thick line represents the fitted model in which OTUs were treated as a random factor. **A**. UMI number (per OTU per soup) poorly predicts within-species variation in input COI copy number (linear mixed-effects model, marginal R^2^ = 0.05, conditional R^2^ = 0.85). **B**. Spike-corrected UMI number successfully predicts within-species variation in input COI copy number (mixed-effects linear model, marginal R^2^ = 0.43, conditional R^2^ = 0.95), but species bias remains, as can be seen in the orders-of-magnitude variation in intercepts. **C**. Non-spike-corrected UMI number poorly predicts within-species variation in input genomic DNA concentration (linear mixed-effects model, marginal R^2^ = 0.01, conditional R^2^ = 0.03). **D**. Spike-corrected UMI number successfully predicts within- species variation in input genomic DNA concentration (linear mixed-effects model, marginal R^2^ = 0.52, conditional R^2^ = 0.94) despite species bias (Figure 1). Model syntax: `lme4::lmer(log.input_gDNA or log.inputCOI_copynumber ∼ log.UMInumber + (log.UMInumber | OTUID) + (1 |soupRep/pcrRep)`) (Bates et al. 2015). Marginal R^2^ represents variance explained by the fixed effect, and conditional R^2^ represents variance explained by the whole model. **(222 words)**

## References

Abrego, N., T. Roslin, T. Huotari, Y. Ji, N. M. Schmidt, J. Wang, D. W. Yu, et al. 2021. Accounting for species interactions is necessary for predicting how arctic arthropod communities respond to climate change. Ecography 44:885–896.

Amend, A. S., K. A. Seifert, and T. D. Bruns. 2010. Quantifying microbial communities with 454 pyrosequencing: does read abundance count? Molecular Ecology 19:5555– 5565.

Bates, D., M. Mächler, B. Bolker, and S. Walker. 2015. Fitting Linear Mixed-Effects Models Using lme4. Journal of Statistical Software 67:1–48.

Bell, K. L., J. Fowler, K. S. Burgess, E. K. Dobbs, D. Gruenewald, B. Lawley, C. Morozumi, et al. 2017. Applying Pollen DNA Metabarcoding to the Study of Plant– Pollinator Interactions. Applications in Plant Sciences 5:1600124.

Brys, R., D. Halfmaerten, S. Neyrinck, Q. Mauvisseau, J. Auwerx, M. Sweet, and J. Mergeay. 2021. Reliable eDNA detection and quantification of the European weather loach (*Misgurnus fossilis*). Journal of Fish Biology 98:399–414.

Caporaso, J. G., J. Kuczynski, J. Stombaugh, K. Bittinger, F. D. Bushman, E. K. Costello, N. Fierer, et al. 2010. QIIME allows analysis of high-throughput community sequencing data. Nature Methods 7:335–336.

Carraro, L., E. Mächler, R. Wüthrich, and F. Altermatt. 2020. Environmental DNA allows upscaling spatial patterns of biodiversity in freshwater ecosystems. Nature Communications 11:3585.

Carraro, L., J. B. Stauffer, and F. Altermatt. 2021. How to design optimal eDNA sampling strategies for biomonitoring in river networks. Environmental DNA 3:157–172.

Clausen, D. S., and A. D. Willis. 2022. Modeling complex measurement error in microbiome experiments. arXiv:2204.12733 [stat].

Creedy, T. J., H. Norman, C. Q. Tang, K. Qing Chin, C. Andujar, P. Arribas, R. S. O’Connor, et al. 2020. A validated workflow for rapid taxonomic assignment and monitoring of a national fauna of bees (Apiformes) using high throughput DNA barcoding. Molecular Ecology Resources 20:40–53.

Deagle, B. E., L. J. Clarke, J. A. Kitchener, A. M. Polanowski, and A. T. Davidson. 2018. Genetic monitoring of open ocean biodiversity: An evaluation of DNA metabarcoding for processing continuous plankton recorder samples. Molecular Ecology Resources 18:391–406.

Deagle, B. E., A. C. Thomas, J. C. McInnes, L. J. Clarke, E. J. Vesterinen, E. L. Clare, T. R. Kartzinel, et al. 2019. Counting with DNA in metabarcoding studies: How should we convert sequence reads to dietary data? Molecular Ecology 28:391–406.

deWaard, J. R., V. Levesque-Beaudin, S. L. deWaard, N. V. Ivanova, J. T. A. McKeown, R. Miskie, S. Naik, et al. 2019. Expedited assessment of terrestrial arthropod diversity by coupling Malaise traps with DNA barcoding. Genome 62:85–95.

Doi, H., K. Fukaya, S. Oka, K. Sato, M. Kondoh, and M. Miya. 2019. Evaluation of detection probabilities at the water-filtering and initial PCR steps in environmental DNA metabarcoding using a multispecies site occupancy model. Scientific Reports 9:3581.

Dorazio, R. M., and R. A. Erickson. 2018. ednaoccupancy : An R package for multiscale occupancy modelling of environmental DNA data. Molecular Ecology Resources 18:368–380.

Edgar, R. C. 2010. Search and clustering orders of magnitude faster than BLAST. Bioinformatics 26:2460–2461.

Elbrecht, V., and F. Leese. 2015. Can DNA-Based Ecosystem Assessments Quantify Species Abundance? Testing Primer Bias and Biomass—Sequence Relationships with an Innovative Metabarcoding Protocol. PLOS ONE 10:e0130324.

Erickson, R. A. 2019. Sampling Designs for Landscape-level eDNA Monitoring Programs. Integr Environ Assess Manag 12.

Ershova, E. A., O. S. Wangensteen, R. Descoteaux, C. Barth-Jensen, and K. Præbel. 2021. Metabarcoding as a quantitative tool for estimating biodiversity and relative biomass of marine zooplankton. (I. Bradbury, ed.)ICES Journal of Marine Science fsab171.

Fields, B., S. Moeskjær, V. Friman, S. U. Andersen, and J. P. W. Young. 2021. MAUI- seq: Metabarcoding using amplicons with unique molecular identifiers to improve error correction. Molecular Ecology Resources 21:703–720.

Folmer, O., M. Black, W. Hoeh, R. Lutz, and R. Vrijenhoek. 1994. DNA primers for amplification of mitochondrial cytochrome c oxidase subunit I from diverse metazoan invertebrates. Molecular Marine Biology and Biotechnology 3:294–299.

Frøslev, T. G., R. Kjøller, H. H. Bruun, R. Ejrnæs, A. K. Brunbjerg, C. Pietroni, and A. J. Hansen. 2017. Algorithm for post-clustering curation of DNA amplicon data yields reliable biodiversity estimates. Nature Communications 8:1188.

Fukaya, K., H. Murakami, S. Yoon, K. Minami, Y. Osada, S. Yamamoto, R. Masuda, et al. 2021. Estimating fish population abundance by integrating quantitative data on environmental DNA and hydrodynamic modelling. Molecular Ecology 30:3057–3067.

Garrido-Sanz, L., M. À. Senar, and J. Piñol. 2021. Relative species abundance estimation in artificial mixtures of insects using mito-metagenomics and a correction factor for the mitochondrial DNA copy number. Molecular Ecology Resources 1755-0998.13464.

Griffin, J. E., E. Matechou, A. S. Buxton, D. Bormpoudakis, and R. A. Griffiths. 2020. Modelling environmental DNA data; Bayesian variable selection accounting for false positive and false negative errors. Journal of the Royal Statistical Society: Series C (Applied Statistics) 69:377–392.

Gueuning, M., D. Ganser, S. Blaser, M. Albrecht, E. Knop, C. Praz, and J. E. Frey. 2019. Evaluating next-generation sequencing (NGS) methods for routine monitoring of wild bees: Metabarcoding, mitogenomics or NGS barcoding. Molecular Ecology Resources 19:847–862.

Harrison, J. G., W. John Calder, B. Shuman, and C. Alex Buerkle. 2021. The quest for absolute abundance: The use of internal standards for DNA-based community ecology. Molecular Ecology Resources 21:30–43.

Hebert, P. D. N., T. W. A. Braukmann, S. W. J. Prosser, S. Ratnasingham, J. R. deWaard, N. V. Ivanova, D. H. Janzen, et al. 2018. A Sequel to Sanger: amplicon sequencing that scales. BMC Genomics 19:219.

Hindson, B. J., K. D. Ness, D. A. Masquelier, P. Belgrader, N. J. Heredia, A. J. Makarewicz, I. J. Bright, et al. 2011. High-Throughput Droplet Digital PCR System for Absolute Quantitation of DNA Copy Number. Analytical Chemistry 83:8604–8610.

Hoshino, T., and F. Inagaki. 2017. Application of Stochastic Labeling with Random- Sequence Barcodes for Simultaneous Quantification and Sequencing of Environmental 16S rRNA Genes. PLOS ONE 12:e0169431.

Hoshino, T., R. Nakao, H. Doi, and T. Minamoto. 2021. Simultaneous absolute quantification and sequencing of fish environmental DNA in a mesocosm by quantitative sequencing technique. Scientific Reports 11:4372.

Iwaszkiewicz-Eggebrecht, E., E. Granqvist, M. Buczek, M. Prus, T. Roslin, A. J. M. Tack, A. F. Andersson, et al. 2022. Optimizing insect metabarcoding using replicated mock communities. preprint.

Ji, Y., T. Huotari, T. Roslin, N. M. Schmidt, J. Wang, D. W. Yu, and O. Ovaskainen. 2020. SPIKEPIPE: A metagenomic pipeline for the accurate quantification of eukaryotic species occurrences and intraspecific abundance change using DNA barcodes or mitogenomes. Molecular Ecology Resources 20:256–267.

Krehenwinkel, H., M. Wolf, J. Y. Lim, A. J. Rominger, W. B. Simison, and R. G. Gillespie. 2017. Estimating and mitigating amplification bias in qualitative and quantitative arthropod metabarcoding. Scientific Reports.

Lang, D., M. Tang, J. Hu, and X. Zhou. 2019. Genome-skimming provides accurate quantification for pollen mixtures. Molecular Ecology Resources 19:1433–1446.

Leray, M., J. Y. Yang, C. P. Meyer, S. C. Mills, N. Agudelo, V. Ranwez, J. T. Boehm, et al. 2013. A new versatile primer set targeting a short fragment of the mitochondrial COI region for metabarcoding metazoan diversity: application for characterizing coral reef fish gut contents. Frontiers in Zoology 10:34.

Levi, T., J. M. Allen, D. Bell, J. Joyce, J. R. Russell, D. A. Tallmon, S. C. Vulstek, et al. 2019. Environmental DNA for the enumeration and management of Pacific salmon. Molecular Ecology Resources 19:597–608.

Li, H. 2015. BFC: correcting Illumina sequencing errors. Bioinformatics 31:2885–2887.

Lundberg, D. S., P. P. N. Ayutthaya, A. Strauß, G. Shirsekar, W.-S. Lo, T. Lahaye, and D. Weigel. 2021. Host-associated microbe PCR (hamPCR) enables convenient measurement of both microbial load and community composition. eLife 10:e66186.

Lundberg, D. S., S. Yourstone, P. Mieczkowski, C. D. Jones, and J. L. Dangl. 2013. Practical innovations for high-throughput amplicon sequencing. Nature Methods 10:999–1002.

Luo, M., Y. Ji, D. Warton, and D. W. Yu. 2022. Dataset for “Extracting abundance information from DNA-based data”. https://datadryad.org/stash/share/0rJ5Yy2PRIv5UpVrCS95Wf7pY0J2R_Hqic6DWyMea D8.

Lyet, A., L. Pellissier, A. Valentini, T. Dejean, A. Hehmeyer, and R. Naidoo. 2021. eDNA sampled from stream networks correlates with camera trap detection rates of terrestrial mammals. Scientific Reports 11:11362.

Masella, A. P., A. K. Bartram, J. M. Truszkowski, D. G. Brown, and J. D. Neufeld. 2012. PANDAseq: paired-end assembler for illumina sequences. BMC Bioinformatics 13:31.

McLaren, M. R., A. D. Willis, and B. J. Callahan. 2019. Consistent and correctable bias in metagenomic sequencing experiments. eLife 8:e46923.

Meier, R., W. Wong, A. Srivathsan, and M. Foo. 2016. $1 DNA barcodes for reconstructing complex phenomes and finding rare species in specimen-rich samples. Cladistics 32:100–110.

Nielsen, M., M. T. P. Gilbert, T. Pape, and K. Bohmann. 2019. A simplified DNA extraction protocol for unsorted bulk arthropod samples that maintains exoskeletal integrity. Environmental DNA 1:144–154.

Pauvert, C., M. Buée, V. Laval, V. Edel-Hermann, L. Fauchery, A. Gautier, I. Lesur, et al. 2019. Bioinformatics matters: The accuracy of plant and soil fungal community data is highly dependent on the metabarcoding pipeline. Fungal Ecology 41:23–33.

Peel, N., L. V. Dicks, M. D. Clark, D. Heavens, L. Percival-Alwyn, C. Cooper, R. G. Davies, et al. 2019. Semi-quantitative characterisation of mixed pollen samples using MinION sequencing and Reverse Metagenomics (RevMet). Methods in Ecology and Evolution 10:1690–1701.

Pierella Karlusich, J. J., E. Pelletier, L. Zinger, F. Lombard, A. Zingone, S. Colin, J. M. Gasol, et al. 2022. A robust approach to estimate relative phytoplankton cell abundances from metagenomes. Molecular Ecology Resources 1755-0998.13592.

Piñol, J., G. Mir, P. Gomez-Polo, and N. Agustí. 2015. Universal and blocking primer mismatches limit the use of high-throughput DNA sequencing for the quantitative metabarcoding of arthropods. Molecular Ecology Resources 15:819–830.

Piñol, J., M. A. Senar, and W. O. C. Symondson. 2019. The choice of universal primers and the characteristics of the species mixture determine when DNA metabarcoding can be quantitative. Molecular Ecology 28:407–419.

Pochardt, M., J. M. Allen, T. Hart, S. D. L. Miller, D. W. Yu, and T. Levi. 2020. Environmental DNA facilitates accurate, inexpensive, and multiyear population estimates of millions of anadromous fish. Molecular Ecology Resources 20:457–467.

R Core Team. 2021. R: A language and environment for statistical computing. R Foundation for Statistical Computing, Vienna, Austria.

Ratnasingham, S. 2019. mBRAVE: The Multiplex Barcode Research And Visualization Environment. Biodiversity Information Science and Standards 3:e37986.

Rognes, T., T. Flouri, B. Nichols, C. Quince, and F. Mahé. 2016. VSEARCH: a versatile open source tool for metagenomics. PeerJ 4:e2584.

Rojahn, J., L. Pearce, D. M. Gleeson, R. P. Duncan, D. M. Gilligan, and J. Bylemans. 2021. The value of quantitative environmental DNA analyses for the management of invasive and endangered native fish. Freshwater Biology fwb.13779.

Rourke, M. L., A. M. Fowler, J. M. Hughes, M. K. Broadhurst, J. D. DiBattista, S. Fielder, J. Wilkes Walburn, et al. 2022. Environmental DNA (eDNA) as a tool for assessing fish biomass: A review of approaches and future considerations for resource surveys. Environmental DNA 4:9–33.

Schenk, J., S. Geisen, N. Kleinboelting, and W. Traunspurger. 2019. Metabarcoding data allow for reliable biomass estimates in the most abundant animals on earth. Metabarcoding and Metagenomics 3:e46704.

Schneider, S., G. W. Taylor, S. C. Kremer, P. Burgess, J. McGroarty, K. Mitsui, A. Zhuang, et al. 2022. Bulk arthropod abundance, biomass and diversity estimation using deep learning for computer vision. Methods in Ecology and Evolution 13:346–357.

Schnell, I. B., K. Bohmann, and M. T. P. Gilbert. 2015. Tag jumps illuminated - reducing sequence-to-sample misidentifications in metabarcoding studies. Molecular Ecology Resources 15:1289–1303.

Schubert, M., S. Lindgreen, and L. Orlando. 2016. AdapterRemoval v2: rapid adapter trimming, identification, and read merging. BMC Research Notes 9:88.

Shelton, A. O., Z. J. Gold, A. J. Jensen, E. D’Agnese, E. Andruszkiewicz, and P. Kelly. 2022a. Toward Quantitative Metabarcoding. BioRXiv.

Shelton, A. O., J. L. O’Donnell, J. F. Samhouri, N. Lowell, G. D. Williams, and R. P. Kelly. 2016. A framework for inferring biological communities from environmental DNA. Ecological Applications 26:1645–1659.

Shelton, A. O., A. Ramón-Laca, A. Wells, J. Clemons, D. Chu, B. E. Feist, R. P. Kelly, et al. 2022b. Environmental DNA provides quantitative estimates of Pacific hake abundance and distribution in the open ocean. Proceedings of the Royal Society B: Biological Sciences 289:20212613.

Silverman, J. D., R. J. Bloom, S. Jiang, H. K. Durand, E. Dallow, S. Mukherjee, and L. A. David. 2021. Measuring and mitigating PCR bias in microbiota datasets. (A. C. McHardy, ed.)PLOS Computational Biology 17:e1009113.

Smets, W., J. W. Leff, M. A. Bradford, R. L. McCulley, S. Lebeer, and N. Fierer. 2016. A method for simultaneous measurement of soil bacterial abundances and community composition via 16S rRNA gene sequencing. Soil Biology and Biochemistry 96:145– 151.

Srivathsan, A., L. Lee, K. Katoh, E. Hartop, S. N. Kutty, J. Wong, D. Yeo, et al. 2021. MinION barcodes: biodiversity discovery and identification by everyone, for everyone. BioRXiv doi:10.1101/2021.03.09.434692.

Stauffer, S., M. Jucker, T. Keggin, V. Marques, M. Andrello, S. Bessudo, M.-C. Cheutin, et al. 2021. How many replicates to accurately estimate fish biodiversity using environmental DNA on coral reefs? BioRxiv doi:10.1101/2021.05.26.445742.

Steinke, D., T. W. Braukmann, L. Manerus, A. Woodhouse, and V. Elbrecht. 2021. Effects of Malaise trap spacing on species richness and composition of terrestrial arthropod bulk samples. Metabarcoding and Metagenomics 5:e59201.

Tang, M., C. J. Hardman, Y. Ji, G. Meng, S. Liu, M. Tan, S. Yang, et al. 2015. High- throughput monitoring of wild bee diversity and abundance via mitogenomics. Methods in Ecology and Evolution 6:1034–1043.

Thalinger, B., A. Rieder, A. Teuffenbach, Y. Pütz, T. Schwerte, J. Wanzenböck, and M. Traugott. 2021. The Effect of Activity, Energy Use, and Species Identity on Environmental DNA Shedding of Freshwater Fish. Frontiers in Ecology and Evolution 9:623718.

Thomas, A. C., B. E. Deagle, J. P. Eveson, C. H. Harsch, and A. W. Trites. 2016. Quantitative DNA metabarcoding: improved estimates of species proportional biomass using correction factors derived from control material. Molecular Ecology Resources 16:714–726.

Tkacz, A., M. Hortala, and P. S. Poole. 2018. Absolute quantitation of microbiota abundance in environmental samples. Microbiome 6:110.

Tsuji, S., R. Inui, R. Nakao, S. Miyazono, M. Saito, T. Kono, and Y. Akamatsu. 2022. Quantitative environmental DNA metabarcoding reflects quantitative capture data of fish community obtained by electrical shocker. BioRXiv.

Ushio, M., H. Murakami, R. Masuda, T. Sado, M. Miya, S. Sakurai, H. Yamanaka, et al. 2018. Quantitative monitoring of multispecies fish environmental DNA using high- throughput sequencing. Metabarcoding and Metagenomics 2:e23297.

Verkuil, Y. I., M. Nicolaus, R. Ubels, M. M. Dietz, J. M. Samplonius, A. Galema, K. Kiekebos, et al. 2020. DNA metabarcoding successfully quantifies relative abundances of arthropod taxa in songbird diets: a validation study using camera-recorded diets. BioRxiv doi:10.1101/2020.11.26.399535.

Wang, Y., U. Naumann, S. T. Wright, and D. I. Warton. 2012. mvabund - an R package for model-based analysis of multivariate abundance data. Methods in Ecology and Evolution 3:471–474.

Warton, D. 2022. Eco-Stats - Data Analysis in Ecology. Methods in Statistical Ecology (1st ed.). Springer International Publishing, Switzerland.

Williamson, B. D., J. P. Hughes, and A. D. Willis. 2021. A multiview model for relative and absolute microbial abundances. Biometrics doi:10.1111/biom.13503.

Wührl, L., C. Pylatiuk, M. Giersch, F. Lapp, T. von Rintelen, M. Balke, S. Schmidt, et al. 2021. DiversityScanner: Robotic discovery of small invertebrates with machine learning methods. BioRxiv doi:10.1101/2021.05.17.444523.

Yang, C., K. Bohmann, X. Wang, W. Cai, N. Wales, Z. Ding, S. Gopalakrishnan, et al. 2021. Biodiversity Soup II: A bulk-sample metabarcoding pipeline emphasizing error reduction. Methods in Ecology and Evolution 12:1252–1264.

Yates, M. C., M. E. Cristescu, and A. M. Derry. 2021a. Integrating physiology and environmental dynamics to operationalize environmental DNA (eDNA) as a means to monitor freshwater macro-organism abundance. Molecular Ecology mec.16202.

Yates, M. C., D. M. Glaser, J. R. Post, M. E. Cristescu, D. J. Fraser, and A. M. Derry. 2021b. The relationship between eDNA particle concentration and organism abundance in nature is strengthened by allometric scaling. Molecular Ecology 30:3068–3082.

Yu, D. W., Y. Ji, B. C. Emerson, X. Wang, C. Ye, C. Yang, and Z. Ding. 2012. Biodiversity soup: metabarcoding of arthropods for rapid biodiversity assessment and biomonitoring. Methods in Ecology and Evolution 3:613–623.

Zepeda-Mendoza, M. L., K. Bohmann, A. Carmona Baez, and M. T. P. Gilbert. 2016. DAMe: a toolkit for the initial processing of datasets with PCR replicates of double- tagged amplicons for DNA metabarcoding analyses. BMC Research Notes 9:255.

Zhou, X., Y. Li, S. Liu, Q. Yang, X. Su, L. Zhou, M. Tang, et al. 2013. Ultra-deep sequencing enables high-fidelity recovery of biodiversity for bulk arthropod samples without PCR amplification. GigaScience 2:4.

