## Supplementary material for "Extracting abundance information from DNA-based data": S4: mock soup mixed model_20220731.Rmd

````
---
title: "Mock soup dilution gradient experiment, general linear mixed-effects models"
author: "Doug"
date: "31 July 2022"
output:
  html_document: default
  pdf_document: default
---

```{r setup, eval=TRUE, include=FALSE}
# boilerplate package loading chunk

# script-specific packages
suppressPackageStartupMessages({
  library(sjPlot)
  library(broom.mixed)
  library(lme4)
  library(MuMIn)
  library(lmtest)
  library(lattice)
  library(equatiomatic)
  library(DataExplorer)
  library(mvabund)
})

# general-use packages
suppressPackageStartupMessages({
  library(tidyverse)
  library(here)
  library(fs)
  library(glue)
  library(readxl)
  library(cowplot)
  library(lubridate)
  library(patchwork)
  library(broom)
  library(ggeffects)
  library(viridis)
  library(arsenal) # for tableby()
  library(waldo) # for compare()
  library(sjmisc) # for rotate_df()
  library(envDocument)
  library(inspectdf)
  library(conflicted)
  library(knitr)
  library(beepr)
  library(pivottabler)
  library(furrr)
  library(scales)
  library(janitor)
  library(tictoc)
  library(ragg)
})

# Sometimes, two or more packages use the same function names. The {conflicted} package lets you set which package gets precedence. For example, the next line enforces that filter() refers to the {dplyr} package. If you want to use the command filter() from a different package, you just need to precede it with the desired package name like this: stats::filter.
conflict_prefer("filter", "dplyr", quiet = TRUE)
conflict_prefer("mutate", "dplyr", quiet = TRUE)
conflict_prefer("select", "dplyr", quiet = TRUE)
conflict_prefer("summarise", "dplyr", quiet = TRUE)
conflict_prefer("first", "dplyr", quiet = TRUE)
conflict_prefer("here", "here", quiet = TRUE)
conflict_prefer("separate", "tidyr", quiet = TRUE)
conflict_prefer("unite", "tidyr", quiet = TRUE)
conflict_prefer("trim", "sjmisc", quiet = TRUE)
conflict_prefer("rescale", "scales", quiet = TRUE)
conflict_prefer("intersect", "dplyr", quiet = TRUE)
conflict_prefer("setdiff", "dplyr", quiet = TRUE) # w/out this, R crashes
conflict_prefer("to_factor", "sjmisc", quiet = TRUE)
conflict_prefer("trim", "glue", quiet = TRUE)
conflict_prefer("discard", "purrr", quiet = TRUE)
conflict_prefer("extract", "tidyr", quiet = TRUE)
conflict_prefer("col_factor", "readr", quiet = TRUE)

# R version
R.version.string
# Print real numbers, not scientific notation.
options(scipen = 999)

env_doc("table")
```

```{r load and explore data}
here()
mydata <- read_table(here("data", "mocksoup_longtable.txt"))

introduce(mydata)
plot_intro(mydata)
plot_missing(mydata)
plot_bar(mydata)
plot_histogram(mydata)
```


Relevant variable names

OTUID: species

sampleName:  sample, dilution, PCR 

readNum:  number of COI reads

mtagNum:  number of UMIs

soup: 7 dilution levels

exp:  experiment (the whole dataset is "within-species")

soupRep:  3 replicates per soup

PCR:  3 PCRs per soupRep

lib: Illumina library

readSpike:  number of spike-in reads (summed over both spike-ins)

mtagSpike:  number of UMIs

readNumSpikeCorr:  readNum / (readSpike/min(readSpike)) 
  We used readSpike/min(readSpike) so that readNumSpikecorr is not too small.

mtagNumSpikeCorr:  mtagNum / (mtagSpike/min(mtagSpike)) 
  We used mtagSpike/min(mtagSpike) so that mtagNumSpikeCorr is not too small.
  
inputCOINum:  input of COI copy number of this species in this soup:  response variable

inputGDNA:  input of genomic DMNA of this species in this soup:  response variable


```{r log predictors and responses}
mydata$log.readNum <- log(mydata$readNum + 1)
mydata$log.inputCOINum <- log(mydata$inputCOINum + 1)
mydata$log.mtagNum <- log(mydata$mtagNum + 1)
mydata$log.inputGDNA <- log(mydata$inputGDNA + 1)
mydata$log.mtagNumSpikeCorr <- log(mydata$mtagNumSpikeCorr + 1)
mydata$log.readNumSpikeCorr <- log(mydata$readNumSpikeCorr + 1)
plot_missing(mydata) # HAP222 and HAP264 OTUs don't have inputCOINum values, amounting to 3.85% of the data
plot_histogram(mydata) 
str(mydata)

```


################################ mixed effects models ############################

Figure 4A
```{r COI non-spike-corrected readNum}
lmer_coi_read1 <- lmer(log.inputCOINum ~ log.readNum +
  (log.readNum | OTUID) +
  (1 | soupRep / pcrRep), data = mydata, REML = F)
# singular fit
lmer_coi_read2 <- lmer(log.inputCOINum ~ log.readNum +
  (1 | OTUID) +
  (1 | soupRep / pcrRep), data = mydata, REML = F)
# singular fit
lmer_coi_read3 <- lmer(log.inputCOINum ~ log.readNum +
  (1 | soupRep / pcrRep), data = mydata, REML = F)
# singular fit
AIC(lmer_coi_read1, lmer_coi_read2, lmer_coi_read3)  
# all the models fit very poorly (not a surprise given that the predictor is not spike-corrected). Of these poor models, choose lmer_coi_read1 to carry out the fixed-effect test


# test fixed effect
lmer_coi_read4 <- lmer(log.inputCOINum ~ 1 +
  (log.readNum | OTUID) +
  (1 | soupRep / pcrRep), data = mydata, REML = F)
# singular fit
AIC(lmer_coi_read1, lmer_coi_read4) # removal of fixed effect increases AIC
anova(lmer_coi_read1, lmer_coi_read4) # keep fixed effect

# estimate parameters with lmer_coi_read1
lmer_coi_read5 <- lmer(log.inputCOINum ~ log.readNum +
  (log.readNum | OTUID) +
  (1 | soupRep / pcrRep), data = mydata, REML = TRUE)
# singular fit
summary(lmer_coi_read5)
r.squaredGLMM(lmer_coi_read5) 
#             R2m       R2c
# [1,] 0.04196802 0.8528916
# R2m = marginal R-squared (variance explained by fixed effects only)
# R2c = conditional R-squared (variance explained by random + fixed effects)

sresid <- resid(lmer_coi_read5, type = "pearson") # Extract the standardised residuals, check underline assumption of normally distibuted
hist(sresid)
plot(lmer_coi_read5) # note that the model fit is very poor, so the residuals plots will also be poor

pred.mm2 <- ggpredict(lmer_coi_read5, terms = c("log.readNum"))

# Plot the predictions
(p2 <- ggplot() +
  geom_line(
    data = pred.mm2, aes(x = x, y = predicted),
    colour = "black", size = 1
  ) + # slope
  geom_ribbon(
    data = pred.mm2, aes(
      x = x, ymin = predicted - std.error,
      ymax = predicted + std.error
    ),
    fill = "lightgrey", alpha = 0.5
  ) +
  geom_point(data = mydata, aes(
    x = log.readNum, y = log.inputCOINum,
    colour = OTUID
  ), size = 0.8) +
  geom_smooth(
    data = mydata, aes(
      x = log.readNum, y = log.inputCOINum,
      group = OTUID
    ),
    method = "lm", se = FALSE, size = 0.3, colour = "black", alpha = 0.5,
    linetype = 1
  ) +
  labs(x = "ln(non-spike-corrected OTU size)", y = "ln(input COI copy number)") +
  theme_cowplot() +
  theme(legend.position = "none"))
```

Figure 4B
```{r COI spike-corrected readNum}
# test random effects models
# with random slope
lmer_coi_readSpikeCorr1 <- lmer(log.inputCOINum ~ log.readNumSpikeCorr +
  (log.readNumSpikeCorr | OTUID) +
  (1 | soupRep / pcrRep), data = mydata, REML = FALSE)
summary(lmer_coi_readSpikeCorr1)
plot(lmer_coi_readSpikeCorr1)

# without random slope
lmer_coi_readSpikeCorr2 <- lmer(log.inputCOINum ~ log.readNumSpikeCorr +
  (1 | OTUID) +
  (1 | soupRep / pcrRep), data = mydata, REML = FALSE)
summary(lmer_coi_readSpikeCorr2)
plot(lmer_coi_readSpikeCorr2)

# without random OTUID 
lmer_coi_readSpikeCorr3 <- lmer(log.inputCOINum ~ log.readNumSpikeCorr +
   (1 | soupRep/pcrRep), data = mydata, REML = FALSE)
# singular fit
summary(lmer_coi_readSpikeCorr3)
plot(lmer_coi_readSpikeCorr3)

AIC(lmer_coi_readSpikeCorr1, lmer_coi_readSpikeCorr2, lmer_coi_readSpikeCorr3) # lmer_coi_readSpikeCorr1 has lower AIC, keep random intercept and slope, keep OTUID, soupRep, pcrRep

# test fixed effects
lmer_coi_readSpikeCorr4 <- lmer(log.inputCOINum ~ 1 +
  (log.readNumSpikeCorr | OTUID) +
  (1 | soupRep / pcrRep), data = mydata, REML = FALSE)
AIC(lmer_coi_readSpikeCorr1, lmer_coi_readSpikeCorr4) # lmer_coi_readSpikeCorr1
anova(lmer_coi_readSpikeCorr1, lmer_coi_readSpikeCorr4) # significantly different. keep fixed effect

# parameter estimates
lmer_coi_readSpikeCorr5 <- lmer(log.inputCOINum ~ log.readNumSpikeCorr +
  (log.readNumSpikeCorr | OTUID) +
  (1 | soupRep / pcrRep), data = mydata, REML = TRUE)
summary(lmer_coi_readSpikeCorr5)
r.squaredGLMM(lmer_coi_readSpikeCorr5) 
#            R2m       R2c
# [1,] 0.4160562 0.9584505
# R2m = marginal R-squared (variance explained by fixed effects only)
# R2c = conditional R-squared (variance explained by random + fixed effects)

sresid <- resid(lmer_coi_readSpikeCorr5, type = "pearson")
hist(sresid)
plot(lmer_coi_readSpikeCorr5)
plot(sresid ~ lmer_coi_readSpikeCorr5@frame[["log.readNumSpikeCorr"]])

pred.mm1 <- ggpredict(lmer_coi_readSpikeCorr5, terms = c("log.readNumSpikeCorr"))

(p1 <- ggplot() +
  geom_line(
    data = pred.mm1, aes(x = x, y = predicted),
    colour = "black", size = 1
  ) +
  geom_ribbon(
    data = pred.mm1, aes(
      x = x, ymin = predicted - std.error,
      ymax = predicted + std.error
    ),
    fill = "lightgrey", alpha = 0.3
  ) +
  geom_point(data = mydata, aes(
    x = log.readNumSpikeCorr, y = log.inputCOINum,
    colour = OTUID
  ), size = 0.8) +
  geom_smooth(
    data = mydata, aes(
      x = log.readNumSpikeCorr, y = log.inputCOINum,
      group = OTUID
    ),
    method = "lm", se = FALSE, size = 0.3, colour = "black", alpha = 0.5,
    linetype = 1
  ) +
  labs(x = "ln(spike-corrected OTU size)", y = "ln(input COI copy number)") +
  theme_cowplot() +
  theme(legend.position = "none"))
```

Figure 4C
```{r gDNA non-spike-corrected readNum}
lmer_gDNA_read1 <- lmer(log.inputGDNA ~ log.readNum +
  (log.readNum | OTUID) +
  (1 | soupRep / pcrRep), data = mydata, REML = F)
# singular fit
lmer_gDNA_read2 <- lmer(log.inputGDNA ~ log.readNum +
  (1 | OTUID) +
  (1 | soupRep / pcrRep), data = mydata, REML = F)
# singular fit
lmer_gDNA_read3 <- lmer(log.inputGDNA ~ log.readNum +
  (1 | soupRep / pcrRep), data = mydata, REML = F)
# singular fits for all models

AIC(lmer_gDNA_read1, lmer_gDNA_read2, lmer_gDNA_read3) # lmer_gDNA_read3
# all the models are poor (since we are using non-spike-corrected readNum)

# test fixed effect
lmer_gDNA_read4 <- lmer(log.inputGDNA ~ 1 +
  (1 | soupRep / pcrRep), data = mydata, REML = F)
# singular fit
AIC(lmer_gDNA_read3, lmer_gDNA_read4) # keep fixed effect
anova(lmer_gDNA_read3, lmer_gDNA_read4) 


######## final model###########
lmer_gDNA_read5 <- lmer(log.inputGDNA ~ log.readNum +
  (1 | soupRep / pcrRep), data = mydata, REML = T)
# singular fit
r.squaredGLMM(lmer_gDNA_read5) # 0.005 0.005
#              R2m         R2c
# [1,] 0.004874685 0.004874685

sresid <- resid(lmer_gDNA_read5, type = "pearson") # Extract the standardised residuals, check underline assumption of normally distibuted
hist(sresid)
plot(sresid ~ mydata$log.readNum) # poor model means weird residuals

pred.mm4 <- ggpredict(lmer_gDNA_read5, terms = c("log.readNum"))

# Plot the predictions
(p4 <- ggplot() +
  geom_line(
    data = pred.mm4, aes(x = x, y = predicted),
    colour = "black", size = 1
  ) + # slope
  geom_ribbon(
    data = pred.mm4, aes(
      x = x, ymin = predicted - std.error,
      ymax = predicted + std.error
    ),
    fill = "lightgrey", alpha = 0.5
  ) +
  geom_point(data = mydata, aes(
    x = log.readNum, y = log.inputGDNA,
    colour = OTUID
  ), size = 0.8) +
  geom_smooth(
    data = mydata, aes(
      x = log.readNum, y = log.inputGDNA,
      group = OTUID
    ),
    method = "lm", se = FALSE, size = 0.3, colour = "black", alpha = 0.5,
    linetype = 1
  ) +
  ylim(2.3, 4.7) +
  labs(x = "ln(non-spike-corrected OTU size)", y = "ln(Input Genomic DNA)") +
  theme_cowplot() +
  theme(legend.position = "none"))
```

Figure 4D
```{r gDNA spike-corrected readNum}
lmer_gDNA_readSpikeCorr1 <- lmer(log.inputGDNA ~ log.readNumSpikeCorr +
   (log.readNumSpikeCorr | OTUID) +
   (1 | soupRep / pcrRep), data = mydata, REML = FALSE)

lmer_gDNA_readSpikeCorr2 <- lmer(log.inputGDNA ~ log.readNumSpikeCorr +
   (1 | OTUID) +
   (1 | soupRep / pcrRep), data = mydata, REML = FALSE)

lmer_gDNA_readSpikeCorr3 <- lmer(log.inputGDNA ~ log.readNumSpikeCorr +
   (1 | soupRep / pcrRep), data = mydata, REML = FALSE)
# singular fit

AIC(lmer_gDNA_readSpikeCorr1, lmer_gDNA_readSpikeCorr2, lmer_gDNA_readSpikeCorr3) # lmer_gDNA_readSpikeCorr1 has lower AIC, use random intercept and slope, keep OTUID, soupRep, pcrRep

# test fixed effects
lmer_gDNA_readSpikeCorr4 <- lmer(log.inputGDNA ~ 1 +
   (log.readNumSpikeCorr | OTUID) +
   (1 | soupRep / pcrRep), data = mydata, REML = F)

AIC(lmer_gDNA_readSpikeCorr1, lmer_gDNA_readSpikeCorr4) # choose lmer_gDNA_readSpikeCorr1
anova(lmer_gDNA_readSpikeCorr1, lmer_gDNA_readSpikeCorr4) # significantly different. keep fixed effect


# fit model
lmer_gDNA_readSpikeCorr5 <- lmer(log.inputGDNA ~ log.readNumSpikeCorr +
   (log.readNumSpikeCorr | OTUID) +
   (1 | soupRep / pcrRep), data = mydata, REML = TRUE)
summary(lmer_gDNA_readSpikeCorr5)
r.squaredGLMM(lmer_gDNA_readSpikeCorr5) # 0.531 0.96
#            R2m       R2c
# [1,] 0.5314312 0.9450565

sresid <- resid(lmer_gDNA_readSpikeCorr5, type = "pearson")
hist(sresid)
plot(lmer_gDNA_readSpikeCorr5)
plot(sresid ~ lmer_gDNA_readSpikeCorr5@frame[["log.readNumSpikeCorr"]]) # wider residuals with small OTUs

pred.mm3 <- ggpredict(lmer_gDNA_readSpikeCorr5, terms = c("log.readNumSpikeCorr"))

(p3 <- ggplot() +
  geom_line(
    data = pred.mm3, aes(x = x, y = predicted),
    colour = "black", size = 1
  ) + 
  geom_ribbon(
    data = pred.mm3, aes(
      x = x, ymin = predicted - std.error,
      ymax = predicted + std.error
    ),
    fill = "lightgrey", alpha = 0.5
  ) +
  geom_point(data = mydata, aes(
    x = log.readNumSpikeCorr, y = log.inputGDNA,
    colour = OTUID
  ), size = 0.8) +
  geom_smooth(
    data = mydata, aes(
      x = log.readNumSpikeCorr, y = log.inputGDNA,
      group = OTUID
    ),
    method = "lm", se = FALSE, size = 0.3, colour = "black", alpha = 0.5,
    linetype = 1
  ) +
  ylim(2.3, 4.7) +
  labs(x = "ln(spike-corrected OTU size)", y = "ln(input genomic DNA)") +
  theme_cowplot() +
  theme(legend.position = "none"))
```

Figure 4. Recovery of within-species abundance change in COI copy number and in genomic DNA concentration in the mock-soup experiment.
```{r plot all}
(p2 + p1) / (p4 + p3)
```


Supplementary information S3.  Using UMI number instead of spike-ins to correct pipeline noise. 

```{r gDNA non-spike-corrected UMI number}
lmer_gDNA_UMI1 <- lmer(log.inputGDNA ~ log.mtagNum +
  (log.mtagNum | OTUID) +
  (1 | soupRep / pcrRep), data = mydata, REML = F)
# singular fit
lmer_gDNA_UMI2 <- lmer(log.inputGDNA ~ log.mtagNum +
  (1 | OTUID) +
  (1 | soupRep / pcrRep), data = mydata, REML = F)
# singular fit
lmer_gDNA_UMI3 <- lmer(log.inputGDNA ~ log.mtagNum +
  (1 | soupRep / pcrRep), data = mydata, REML = F)
# singular fit

AIC(lmer_gDNA_UMI1, lmer_gDNA_UMI2, lmer_gDNA_UMI3) # lmer_gDNA_UMI1

# fixed effect
lmer_gDNA_UMI4 <- lmer(log.inputGDNA ~ 1 +
  (log.mtagNum | OTUID) +
  (1 | soupRep / pcrRep), data = mydata, REML = F)
AIC(lmer_gDNA_UMI1, lmer_gDNA_UMI4) # lmer_gDNA_UMI1
anova(lmer_gDNA_UMI1, lmer_gDNA_UMI4) # lmer_gDNA_UMI1

lmer_gDNA_UMI5 <- lmer(log.inputGDNA ~ log.mtagNum +
  (log.mtagNum | OTUID) +
  (1 | soupRep / pcrRep), data = mydata, REML = T)

r.squaredGLMM(lmer_gDNA_UMI5) # MuMIn package 0.015 0.033
# R2m = marginal R-squared (variance explained by fixed effects only)
# R2c = conditional R-squared (variance explained by random + fixed effects)

sresid <- resid(lmer_gDNA_UMI5, type = "pearson") # Extract the standardised residuals, check underline assumption of normally distibuted
hist(sresid)

plot(lmer_gDNA_UMI5) # gives a heteroscedasticity plot

plot(sresid ~ mydata$log.mtagNum)

pred.mm5 <- ggpredict(lmer_gDNA_UMI5, terms = c("log.mtagNum"))

# Plot the predictions
(p5 <- ggplot() +
  geom_line(data = pred.mm5, aes(x = x, y = predicted),
            colour = "black", size = 1) + # slope
  geom_ribbon(
    data = pred.mm5, aes(
      x = x, ymin = predicted - std.error,
      ymax = predicted + std.error
    ),
    fill = "lightgrey", alpha = 0.5
  ) +
  geom_point(data = mydata, aes(
    x = log.mtagNum, y = log.inputGDNA,
    colour = OTUID
  ), size = 0.8) +
  geom_smooth(
    data = mydata, aes(
      x = log.mtagNum, y = log.inputGDNA,
      group = OTUID
    ),
    method = "lm", se = FALSE, size = 0.3, colour = "black", alpha = 0.5,
    linetype = 1
  ) +
  labs(y = "ln(Input Genomic DNA)", x = "ln(non-spike-corrected UMI number)") +
  theme_cowplot() +
  theme(legend.position = "none"))
```

```{r gDNA spike-corrected UMI number}
lmer_gDNA_UMIspikeCorr1 <- lmer(log.inputGDNA ~ log.mtagNumSpikeCorr +
  (log.mtagNumSpikeCorr | OTUID) +
  (1 | soupRep / pcrRep), data = mydata, REML = F)
lmer_gDNA_UMIspikeCorr2 <- lmer(log.inputGDNA ~ log.mtagNumSpikeCorr +
  (1 | OTUID) +
  (1 | soupRep / pcrRep), data = mydata, REML = F)
lmer_gDNA_UMIspikeCorr3 <- lmer(log.inputGDNA ~ log.mtagNumSpikeCorr +
  (1 | soupRep / pcrRep), data = mydata, REML = F)
# singular fit

AIC(lmer_gDNA_UMIspikeCorr1, lmer_gDNA_UMIspikeCorr2, lmer_gDNA_UMIspikeCorr3) # lmer_gDNA_UMIspikeCorr1

# fixed effect
lmer_gDNA_UMIspikeCorr4 <- lmer(log.inputGDNA ~ 1 +
  (log.mtagNumSpikeCorr | OTUID) +
  (1 | soupRep / pcrRep), data = mydata, REML = F)
# failure to converge
AIC(lmer_gDNA_UMIspikeCorr1, lmer_gDNA_UMIspikeCorr4) # lmer_gDNA_UMIspikeCorr1

# parameter estimate
lmer_gDNA_UMIspikeCorr5 <- lmer(log.inputGDNA ~ log.mtagNumSpikeCorr +
  (log.mtagNumSpikeCorr | OTUID) +
  (1 | soupRep / pcrRep), data = mydata, REML = T)

r.squaredGLMM(lmer_gDNA_UMIspikeCorr5) # 0.52 0.94
sresid <- resid(lmer_gDNA_UMIspikeCorr5, type = "pearson")
hist(sresid)
plot(lmer_gDNA_UMIspikeCorr5)
plot(sresid ~ mydata$log.mtagNumSpikeCorr)

pred.mm6 <- ggpredict(lmer_gDNA_UMIspikeCorr5, terms = c("log.mtagNumSpikeCorr"))

(p6 <- ggplot() +
  geom_line(data = pred.mm6, aes(x = x, y = predicted),
            colour = "black", size = 1) + # slope
  geom_ribbon(
    data = pred.mm6, aes(
      x = x, ymin = predicted - std.error,
      ymax = predicted + std.error
    ),
    fill = "lightgrey", alpha = 0.5
  ) +
  geom_point(data = mydata, aes(
    x = log.mtagNumSpikeCorr, y = log.inputGDNA,
    colour = OTUID
  ), size = 0.8) +
  geom_smooth(
    data = mydata, aes(
      x = log.mtagNumSpikeCorr, y = log.inputGDNA,
      group = OTUID
    ),
    method = "lm", se = FALSE, size = 0.3, colour = "black", alpha = 0.5,
    linetype = 1
  ) +
  ylim(2.3, 4.7) + 
  labs(y = "ln(Input Genomic DNA)", x = "ln(spike-corrected UMI number)") +
  theme_cowplot() +
  theme(legend.position = "none"))
```

```{r COI non-spike-corrected UMI number}
lmer_COI_UMIspike1 <- lmer(log.inputCOINum ~ log.mtagNum +
  (log.mtagNum | OTUID) +
  (1 | soupRep / pcrRep), data = mydata, REML = F)
# singular fit
lmer_COI_UMIspike2 <- lmer(log.inputCOINum ~ log.mtagNum +
  (1 | OTUID) +
  (1 | soupRep / pcrRep), data = mydata, REML = F)
# singular fit
lmer_COI_UMIspike3 <- lmer(log.inputCOINum ~ log.mtagNum +
  (1 | soupRep / pcrRep), data = mydata, REML = F)
# singular fit

AIC(lmer_COI_UMIspike1, lmer_COI_UMIspike2, lmer_COI_UMIspike3) # lmer_gDNA_UMIspikeCorr1

# fixed effect
lmer_COI_UMIspike4 <- lmer(log.inputCOINum ~ 1 +
  (1 | OTUID) +
  (1 | soupRep / pcrRep), data = mydata, REML = F)
# singular fit
AIC(lmer_COI_UMIspike1, lmer_COI_UMIspike4) # lmer_COI_UMIspikeCorr1
anova(lmer_COI_UMIspike1, lmer_COI_UMIspike4) 

# parameter estimate
lmer_COI_UMIspike5 <- lmer(log.inputCOINum ~ log.mtagNum +
  (log.mtagNum | OTUID) +
  (1 | soupRep / pcrRep), data = mydata, REML = T)
# singular fit
r.squaredGLMM(lmer_COI_UMIspike5) # 0.054 0.85
# R2m = marginal R-squared (variance explained by fixed effects only)
# R2c = conditional R-squared (variance explained by random + fixed effects)

sresid <- resid(lmer_COI_UMIspike5, type = "pearson") # Extract the standardised residuals, check underline assumption of normally distibuted
hist(sresid)

plot(lmer_COI_UMIspike5) # gives a heteroscedasticity plot

# plot(sresid ~ mydata$log.mtagNum)

pred.mm7 <- ggpredict(lmer_COI_UMIspike5, terms = c("log.mtagNum"))

# Plot the predictions
(p7 <- ggplot() +
  geom_line(data = pred.mm7, aes(x = x, y = predicted),
            colour = "black", size = 1) + # slope
  geom_ribbon(
    data = pred.mm7, aes(
      x = x, ymin = predicted - std.error,
      ymax = predicted + std.error
    ),
    fill = "lightgrey", alpha = 0.5
  ) +
  geom_point(data = mydata, aes(
    x = log.mtagNum, y = log.inputCOINum,
    colour = OTUID
  ), size = 0.8) +
  geom_smooth(
    data = mydata, aes(
      x = log.readNum, y = log.inputCOINum,
      group = OTUID
    ),
    method = "lm", se = FALSE, size = 0.3, colour = "black", alpha = 0.5,
    linetype = 1
  ) +
  labs(y = "ln(Input COI copy number)", x = "ln(non-spike-corrected UMI number)") +
  theme_cowplot() +
  theme(legend.position = "none"))
```

```{r COI spike-corrected UMI number}
lmer_COI_UMIspikeCorr1 <- lmer(log.inputCOINum ~ log.mtagNumSpikeCorr +
  (log.mtagNumSpikeCorr | OTUID) +
  (1 | soupRep / pcrRep), data = mydata, REML = F)
lmer_COI_UMIspikeCorr2 <- lmer(log.inputCOINum ~ log.mtagNumSpikeCorr +
  (1 | OTUID) +
  (1 | soupRep / pcrRep), data = mydata, REML = F)
lmer_COI_UMIspikeCorr3 <- lmer(log.inputCOINum ~ log.mtagNumSpikeCorr +
  (1 | soupRep / pcrRep), data = mydata, REML = F)
# singular fit

AIC(lmer_COI_UMIspikeCorr1, lmer_COI_UMIspikeCorr2, lmer_COI_UMIspikeCorr3) # lmer_gDNA_UMIspikeCorr1

# fixed effect
lmer_COI_UMIspikeCorr4 <- lmer(log.inputCOINum ~ 1 +
  (log.mtagNumSpikeCorr | OTUID) +
  (1 | soupRep / pcrRep), data = mydata, REML = F)

AIC(lmer_COI_UMIspikeCorr1, lmer_COI_UMIspikeCorr4) # lmer_COI_UMIspikeCorr1
anova(lmer_COI_UMIspikeCorr1, lmer_COI_UMIspikeCorr4) 

# estimate parameters
lmer_COI_UMIspikeCorr5 <- lmer(log.inputCOINum ~ log.mtagNumSpikeCorr +
  (log.mtagNumSpikeCorr | OTUID) +
  (1 | soupRep / pcrRep), data = mydata, REML = T)
r.squaredGLMM(lmer_COI_UMIspikeCorr5) # 0.434 0.95

sresid <- resid(lmer_COI_UMIspikeCorr5, type = "pearson")
hist(sresid)
plot(lmer_COI_UMIspikeCorr5)

pred.mm8 <- ggpredict(lmer_COI_UMIspikeCorr5, terms = c("log.mtagNumSpikeCorr"))

(p8 <- ggplot() +
  geom_line(data = pred.mm8, aes(x = x, y = predicted),
            colour = "black", size = 1) + # slope
  geom_ribbon(
    data = pred.mm8, aes(
      x = x, ymin = predicted - std.error,
      ymax = predicted + std.error
    ),
    fill = "lightgrey", alpha = 0.5
  ) +
  geom_point(data = mydata, aes(
    x = log.mtagNumSpikeCorr, y = log.inputCOINum,
    colour = OTUID
  ), size = 0.8) +
  geom_smooth(
    data = mydata, aes(
      x = log.mtagNumSpikeCorr, y = log.inputCOINum,
      group = OTUID
    ),
    method = "lm", se = FALSE, size = 0.3, colour = "black", alpha = 0.5,
    linetype = 1
  ) +
  labs(y = "ln(Input COI copy number)", x = "ln(spike-corrected UMI number)") +
  theme_cowplot() +
  theme(legend.position = "none"))


```

Figure S3 UMI on mocksoups
```{r figures}
(p7 + p8) / (p5 + p6) + plot_annotation(tag_levels = 'A') 
```


#### Test of Warton spike-in estimator on full dataset, with OTUID and soupRep/pcrRep   

Model-based pipeline-noise estimation: offset0 (ã from Eq. 2)

The following code uses the model-based estimator to estimate a spike-in corrector directly from the dataset, which we find is positive albeit weakly correlated with the spike-in (R2 = 0.23 and 0.18). N.B. The mvabund model does not take into account the nested experimental design (soupRep/PCRrep)
reformat data
```{r reformat to mydata2}
mydata2 <- mydata %>% 
  select(-mtagNum, -exp, -lib, -sampleName, -PCR, -readSpike, -mtagSpike,
         -readNumSpikeCorr, -mtagNumSpikeCorr, -inputCOINum, -inputGDNA,
         -starts_with("log.")) %>% 
  pivot_wider(names_from = OTUID, values_from = readNum)

otu <- mydata2 %>% 
  select(starts_with("HAP"))

X <- mydata2 %>% 
  select(!starts_with("HAP"))
```

```{r calculate offset0 for mydata2}
otu <- mvabund(otu)
# fit0 <- manyglm(otu ~ 1, family = "negative.binomial") # if no predictors
fit0 <- manyglm(otu ~ soup, family = "negative.binomial", data = X) # data = the env covariates. otu is the otu table
# fit0 <- manyglm(otu ~ soup, family = binomial("cloglog")) # for p/a data
offset0 <- log(rowSums(otu)) - log(rowSums(fitted(fit0))) 
mydata2$offset0 <- offset0
mydata2 <- mydata2 %>% 
  relocate(offset0, .after = pcrRep) %>% 
  pivot_longer(cols = starts_with("HAP"), 
               names_to = "OTUID", 
               values_to = "readNum") %>% 
  mutate(
    readNumOffsetCorr = readNum / exp(offset0) # correction
  )

mydata2 <- mydata %>% 
  select(OTUID, soup, soupRep, pcrRep, inputCOINum, inputGDNA, 
         readSpike, mtagSpike) %>% # physical spike-in read numbers
  left_join(mydata2) %>% 
  mutate(
    log.inputCOINum = log(inputCOINum + 1),
    log.inputGDNA = log(inputGDNA + 1),
    log.readNum = log(readNum + 1),
    log.readNumOffsetCorr = log(readNumOffsetCorr + 1),
    log.readSpike = log(readSpike + 1),
    log.mtagSpike = log(mtagSpike + 1)
  )

names(mydata2)

#  [1] "OTUID"                 "soup"                  "soupRep"              
#  [4] "pcrRep"                "inputCOINum"           "inputGDNA"            
#  [7] "readSpike"             "mtagSpike"             "offset0"              
# [10] "readNum"               "readNumOffsetCorr"     "log.inputCOINum"      
# [13] "log.inputGDNA"         "log.readNum"           "log.readNumOffsetCorr"
# [16] "log.readSpike"         "log.mtagSpike"  

# offset0 is the estimated spike-in
# log.readSpike is the log of the spike-in reads
# log.mtagSpike is the log of the UMIs, which is another way of doing spike-ins

# offset0 is correlated with 
ggplot(data = mydata2, aes(x = (offset0), y = (log.readSpike))) +
  geom_point(aes(colour = soup)) +
  geom_smooth(method = "lm", se = FALSE, size = 0.3, colour = "black", 
              alpha = 0.5, linetype = 1) +
  labs(x = "offset0", y = "ln(readSpike)") +
  theme_cowplot() +
  theme(legend.position = "right")

summary(lm(log.readSpike ~ offset0, data = mydata2))
# R2 = 0.2307

ggplot(data = mydata2, aes(x = offset0, y = log.mtagSpike, group = OTUID)) +
  geom_point(aes(colour = soup)) +
  geom_smooth(method = "lm", se = FALSE, size = 0.3, colour = "black", 
              alpha = 0.5, linetype = 1) +
  labs(x = "offset0", y = "ln(mtagSpike)") +
  theme_cowplot() +
  theme(legend.position = "none")

summary(lm(log.mtagSpike ~ offset0, data = mydata2))
# R2 = 0.1846
```

We now use these (inferior) model-estimated spike-ins to correct the OTU sizes and then try to predict the input COI copy number and the input genomic DNA, just as in Figure 4. The results are poorer than the spike-in (no surprise), but better than no correction, as can be seen via inspection of the full model regression lines (the thick black lines) 

prediction of input COI copy number
```{r COI offset-corrected readNum}
# test random effects
lmer_coi_readOffsetCorr1 <- lmer(log.inputCOINum ~ log.readNumOffsetCorr +
  (log.readNumOffsetCorr | OTUID) +
  (1 | soupRep / pcrRep), data = mydata2, REML = FALSE)
summary(lmer_coi_readOffsetCorr1)
plot(lmer_coi_readOffsetCorr1)
# singular fit caused by zero variance in pcrRep:soupRep and soupRep, but these random factors must be retained because they are part of the experimental design
lmer_coi_readOffsetCorr2 <- lmer(log.inputCOINum ~ log.readNumOffsetCorr +
  (1 | OTUID) +
  (1 | soupRep / pcrRep), data = mydata2, REML = FALSE)
summary(lmer_coi_readOffsetCorr2)
plot(lmer_coi_readOffsetCorr2)
# singular fit
lmer_coi_readOffsetCorr3 <- lmer(log.inputCOINum ~ log.readNumOffsetCorr +
   (1 | soupRep/pcrRep), data = mydata2, REML = FALSE)
# singular fit
summary(lmer_coi_readOffsetCorr3)
plot(lmer_coi_readOffsetCorr3)

AIC(lmer_coi_readOffsetCorr1, lmer_coi_readOffsetCorr2, lmer_coi_readOffsetCorr3) # lmer_coi_readOffsetCorr1 has lowest AIC, keep random intercept and slope, keep OTUID, soupRep/pcrRep

# test fixed effects
lmer_coi_readOffsetCorr4 <- lmer(log.inputCOINum ~ 1 +
  (log.readNumOffsetCorr | OTUID) +
  (1 | soupRep / pcrRep), data = mydata2, REML = F)
# singular fit
AIC(lmer_coi_readOffsetCorr1, lmer_coi_readOffsetCorr4) # lmer_coi_readOffsetCorr1 lowest AIC
anova(lmer_coi_readOffsetCorr1, lmer_coi_readOffsetCorr4) # significantly different. keep fixed effect

# parameter estimates
lmer_coi_readOffsetCorr5 <- lmer(log.inputCOINum ~ log.readNumOffsetCorr +
  (log.readNumOffsetCorr | OTUID) +
  (1 | soupRep / pcrRep), data = mydata2, REML = TRUE)
# singular fit
summary(lmer_coi_readOffsetCorr5)
r.squaredGLMM(lmer_coi_readOffsetCorr5) # 0.1182 0.8676

sresid <- resid(lmer_coi_readOffsetCorr5, type = "pearson")
hist(sresid)
plot(lmer_coi_readOffsetCorr5)

pred.mm1 <- ggpredict(lmer_coi_readOffsetCorr5, 
                      terms = c("log.readNumOffsetCorr[all]"))

(p9 <- ggplot() +
  geom_line(
    data = pred.mm1, aes(x = x, y = predicted),
    colour = "black", size = 1
  ) +
  geom_ribbon(
    data = pred.mm1, aes(
      x = x, ymin = predicted - std.error,
      ymax = predicted + std.error
    ),
    fill = "lightgrey", alpha = 0.3
  ) +
  geom_point(data = mydata2, aes(
    x = log.readNumOffsetCorr, y = log.inputCOINum,
    colour = OTUID
  ), size = 0.8) +
  geom_smooth(
    data = mydata2, aes(
      x = log.readNumOffsetCorr, y = log.inputCOINum,
      group = OTUID
    ),
    method = "lm", se = FALSE, size = 0.3, colour = "black", alpha = 0.5,
    linetype = 1
  ) +
  labs(x = "ln(Offset-corrected OTU size)", y = "ln(input COI copy number)") +
  theme_cowplot() +
  theme(legend.position = "none"))
```

should be exactly the same as "COI non-spike-corrected readNum"
```{r COI non-offset-corrected readNum}
lmer_coi_read1 <- lmer(log.inputCOINum ~ log.readNum +
  (log.readNum | OTUID) +
  (1 | soupRep / pcrRep), data = mydata2, REML = F)
# singular fit
lmer_coi_read2 <- lmer(log.inputCOINum ~ log.readNum +
  (1 | OTUID) +
  (1 | soupRep / pcrRep), data = mydata2, REML = F)
# singular fit
lmer_coi_read3 <- lmer(log.inputCOINum ~ log.readNum +
  (1 | soupRep / pcrRep), data = mydata2, REML = F)
# singular fit
AIC(lmer_coi_read1, lmer_coi_read2, lmer_coi_read3) # choose lmer_coi_read1

# test fixed effect
lmer_coi_read4 <- lmer(log.inputCOINum ~ 1 +
  (log.readNum | OTUID) +
  (1 | soupRep / pcrRep), data = mydata2, REML = F)
# singular fit
AIC(lmer_coi_read1, lmer_coi_read4)
anova(lmer_coi_read1, lmer_coi_read4) # keep fixed effect

# estimate parameters with lmer_coi_read1
lmer_coi_read5 <- lmer(log.inputCOINum ~ log.readNum +
  (log.readNum | OTUID) +
  (1 | soupRep / pcrRep), data = mydata2, REML = TRUE)
# singular fit
summary(lmer_coi_read5)
r.squaredGLMM(lmer_coi_read5) # 0.042 0.85
# R2m = marginal R-squared (variance explained by fixed effects only)
# R2c = conditional R-squared (variance explained by random + fixed effects)

sresid <- resid(lmer_coi_read5, type = "pearson") # Extract the standardised residuals, check underline assumption of normally distibuted
hist(sresid)
plot(lmer_coi_read5) # gives a heteroscedasticity plot

pred.mm2 <- ggpredict(lmer_coi_read5, terms = c("log.readNum"))

# Plot the predictions
(p10 <- ggplot() +
  geom_line(
    data = pred.mm2, aes(x = x, y = predicted),
    colour = "black", size = 1
  ) + # slope
  geom_ribbon(
    data = pred.mm2, aes(
      x = x, ymin = predicted - std.error,
      ymax = predicted + std.error
    ),
    fill = "lightgrey", alpha = 0.5
  ) +
  geom_point(data = mydata2, aes(
    x = log.readNum, y = log.inputCOINum,
    colour = OTUID
  ), size = 0.8) +
  geom_smooth(
    data = mydata2, aes(
      x = log.readNum, y = log.inputCOINum,
      group = OTUID
    ),
    method = "lm", se = FALSE, size = 0.3, colour = "black", alpha = 0.5,
    linetype = 1
  ) +
  labs(x = "ln(non-Offset-corrected OTU size)", y = "ln(input COI copy number)") +
  theme_cowplot() +
  theme(legend.position = "none"))

p10 + p9
```

prediction of input genomic DNA
```{r gDNA offset-corrected readNum}
lmer_gDNA_readOffsetCorr1 <- lmer(log.inputGDNA ~ log.readNumOffsetCorr +
   (log.readNumOffsetCorr | OTUID) +
   (1 | soupRep / pcrRep), data = mydata2, REML = FALSE)
# singular fit
summary(lmer_gDNA_readOffsetCorr1)
lmer_gDNA_readOffsetCorr2 <- lmer(log.inputGDNA ~ log.readNumOffsetCorr +
   (1 | OTUID) +
   (1 | soupRep / pcrRep), data = mydata2, REML = FALSE)
# singular fit
lmer_gDNA_readOffsetCorr3 <- lmer(log.inputGDNA ~ log.readNumOffsetCorr +
   (1 | soupRep / pcrRep), data = mydata2, REML = FALSE)
# singular fit

AIC(lmer_gDNA_readOffsetCorr1, lmer_gDNA_readOffsetCorr2, lmer_gDNA_readOffsetCorr3) # lmer_gDNA_readOffsetCorr1 has lower AIC, use random intercept and slope, keep OTUID, soupRep, pcrRep

# test fixed effects
lmer_gDNA_readOffsetCorr4 <- lmer(log.inputGDNA ~ 1 +
   (log.readNumOffsetCorr | OTUID) +
   (1 | soupRep / pcrRep), data = mydata2, REML = F)
# singular fit
AIC(lmer_gDNA_readOffsetCorr1, lmer_gDNA_readOffsetCorr4) # choose lmer_gDNA_readOffsetCorr1
anova(lmer_gDNA_readOffsetCorr1, lmer_gDNA_readOffsetCorr4) # significantly different. keep fixed effect


# fit model
lmer_gDNA_readOffsetCorr5 <- lmer(log.inputGDNA ~ log.readNumOffsetCorr +
   (log.readNumOffsetCorr | OTUID) +
   (1 | soupRep / pcrRep), data = mydata2, REML = TRUE)
# singular fit
summary(lmer_gDNA_readOffsetCorr5)
r.squaredGLMM(lmer_gDNA_readOffsetCorr5) # 0.213 0.571

sresid <- resid(lmer_gDNA_readOffsetCorr5, type = "pearson")
hist(sresid)
plot(lmer_gDNA_readOffsetCorr5)
plot(sresid ~ mydata2$log.readNumOffsetCorr)
#
pred.mm3 <- ggpredict(lmer_gDNA_readOffsetCorr5, 
                      terms = c("log.readNumOffsetCorr[all]"))

(p11 <- ggplot() +
  geom_line(
    data = pred.mm3, aes(x = x, y = predicted),
    colour = "black", size = 1
  ) + 
  geom_ribbon(
    data = pred.mm3, aes(
      x = x, ymin = predicted - std.error,
      ymax = predicted + std.error
    ),
    fill = "lightgrey", alpha = 0.5
  ) +
  geom_point(data = mydata2, aes(
    x = log.readNumOffsetCorr, y = log.inputGDNA,
    colour = OTUID
  ), size = 0.8) +
  geom_smooth(
    data = mydata2, aes(
      x = log.readNumOffsetCorr, y = log.inputGDNA,
      group = OTUID
    ),
    method = "lm", se = FALSE, size = 0.3, colour = "black", alpha = 0.5,
    linetype = 1
  ) +
  ylim(2.3, 4.7) +
  labs(x = "ln(Offset-corrected OTU size)", y = "ln(input genomic DNA)") +
  theme_cowplot() +
  theme(legend.position = "none"))
```

should be exactly the same as "gDNA non-spike-corrected readNum"
```{r gDNA non-offset-corrected readNum}
lmer_gDNA_read1 <- lmer(log.inputGDNA ~ log.readNum +
  (log.readNum | OTUID) +
  (1 | soupRep / pcrRep), data = mydata2, REML = F)
# singular fit
summary(lmer_gDNA_read1)
lmer_gDNA_read2 <- lmer(log.inputGDNA ~ log.readNum +
  (1 | OTUID) +
  (1 | soupRep / pcrRep), data = mydata2, REML = F)
# singular fit
lmer_gDNA_read3 <- lmer(log.inputGDNA ~ log.readNum +
  (1 | soupRep / pcrRep), data = mydata2, REML = F)
# singular fit

AIC(lmer_gDNA_read1, lmer_gDNA_read2, lmer_gDNA_read3) # lmer_gDNA_read3

# fixed effect
lmer_gDNA_read4 <- lmer(log.inputGDNA ~ 1 +
  (1 | soupRep / pcrRep), data = mydata2, REML = F)
# singular fit
AIC(lmer_gDNA_read3, lmer_gDNA_read4) # keep fixed effect
anova(lmer_gDNA_read3, lmer_gDNA_read4) 


######## final model###########
lmer_gDNA_read5 <- lmer(log.inputGDNA ~ log.readNum +
  (1 | soupRep / pcrRep), data = mydata2, REML = T)
# singular fit
r.squaredGLMM(lmer_gDNA_read5) # 0.00487 0.005
# R2m = marginal R-squared (variance explained by fixed effects only)
# R2c = conditional R-squared (variance explained by random + fixed effects)

sresid <- resid(lmer_gDNA_read5, type = "pearson") # Extract the standardised residuals, check underline assumption of normally distibuted
hist(sresid)
plot(lmer_gDNA_read5) # gives a heteroscedasticity plot
plot(sresid ~ mydata2$log.readNum)

pred.mm4 <- ggpredict(lmer_gDNA_read5, terms = c("log.readNum[all]"))

# Plot the predictions
(p12 <- ggplot() +
  geom_line(
    data = pred.mm4, aes(x = x, y = predicted),
    colour = "black", size = 1
  ) + # slope
  geom_ribbon(
    data = pred.mm4, aes(
      x = x, ymin = predicted - std.error,
      ymax = predicted + std.error
    ),
    fill = "lightgrey", alpha = 0.5
  ) +
  geom_point(data = mydata2, aes(
    x = log.readNum, y = log.inputGDNA,
    colour = OTUID
  ), size = 0.8) +
  geom_smooth(
    data = mydata2, aes(
      x = log.readNum, y = log.inputGDNA,
      group = OTUID
    ),
    method = "lm", se = FALSE, size = 0.3, colour = "black", alpha = 0.5,
    linetype = 1
  ) +
  ylim(2.3, 4.7) +
  labs(x = "ln(non-Offset-corrected OTU size)", y = "ln(Input Genomic DNA)") +
  theme_cowplot() +
  theme(legend.position = "none"))

p11 + p12
```

There is only a small benefit of using the model-estimated spike-in correctors on this dataset. The thick black regression lines are the whole-model fits, and they are steeper in the model-estimated "offset-corrected" models on the right. The thin lines are fit to each OTU separately. 
```{r}
(p10 + p9) / (p12 + p11)
```
````
