## Supplementary material for "Extracting abundance information from DNA-based data": S4: mock-soup-mixed-model_20220731_OPEN_IN_CHROME.html

Mock soup dilution gradient experiment, general linear mixed-effects models


### Mock soup dilution gradient experiment, general linear mixed-effects models

###### Doug

###### 31 July 2022

```
here()
```

```
## [1] "/Users/douglasyu/Library/CloudStorage/Dropbox/Working_docs/Luo_Mingjie_Oregon/qSeq paper/qseq_paper/Revision/supplementary information/S4_Mock_soup_mixed_effect_model_20220731"
```

```
mydata <- read_table(here("data", "mocksoup_longtable.txt"))
```

```
## 
## ── Column specification ────────────────────────────────────────────────────────
## cols(
##   OTUID = col_character(),
##   sampleName = col_character(),
##   readNum = col_double(),
##   mtagNum = col_double(),
##   soup = col_character(),
##   exp = col_character(),
##   soupRep = col_double(),
##   PCR = col_character(),
##   lib = col_double(),
##   pcrRep = col_double(),
##   readSpike = col_double(),
##   mtagSpike = col_double(),
##   readNumSpikeCorr = col_double(),
##   mtagNumSpikeCorr = col_double(),
##   inputCOINum = col_double(),
##   inputGDNA = col_double()
## )
```

```
introduce(mydata)
```

```
## # A tibble: 1 × 9
##    rows columns discrete_columns conti…¹ all_m…² total…³ compl…⁴ total…⁵ memor…⁶
##   <int>   <int>            <int>   <int>   <int>   <int>   <int>   <int>   <dbl>
## 1  3276      16                5      11       0     126    3150   52416  433936
## # … with abbreviated variable names ¹​continuous_columns, ²​all_missing_columns,
## #   ³​total_missing_values, ⁴​complete_rows, ⁵​total_observations, ⁶​memory_usage
```

```
plot_intro(mydata)
```

```
plot_missing(mydata)
```

```
plot_bar(mydata)
```

```
## 2 columns ignored with more than 50 categories.
## OTUID: 52 categories
## sampleName: 63 categories
```

```
plot_histogram(mydata)
```

Relevant variable names

OTUID: species

sampleName: sample, dilution, PCR

readNum: number of COI reads

mtagNum: number of UMIs

soup: 7 dilution levels

exp: experiment (the whole dataset is “within-species”)

soupRep: 3 replicates per soup

PCR: 3 PCRs per soupRep

lib: Illumina library

readSpike: number of spike-in reads (summed over both spike-ins)

mtagSpike: number of UMIs

readNumSpikeCorr: readNum / (readSpike/min(readSpike)) We used
readSpike/min(readSpike) so that readNumSpikecorr is not too small.

mtagNumSpikeCorr: mtagNum / (mtagSpike/min(mtagSpike)) We used
mtagSpike/min(mtagSpike) so that mtagNumSpikeCorr is not too small.

inputCOINum: input of COI copy number of this species in this soup:
response variable

inputGDNA: input of genomic DMNA of this species in this soup:
response variable

```
mydata$log.readNum <- log(mydata$readNum + 1)
mydata$log.inputCOINum <- log(mydata$inputCOINum + 1)
mydata$log.mtagNum <- log(mydata$mtagNum + 1)
mydata$log.inputGDNA <- log(mydata$inputGDNA + 1)
mydata$log.mtagNumSpikeCorr <- log(mydata$mtagNumSpikeCorr + 1)
mydata$log.readNumSpikeCorr <- log(mydata$readNumSpikeCorr + 1)
plot_missing(mydata) # HAP222 and HAP264 OTUs don't have inputCOINum values, amounting to 3.85% of the data
```

```
plot_histogram(mydata)
```

```
str(mydata)
```

```
## spec_tbl_df [3,276 × 22] (S3: spec_tbl_df/tbl_df/tbl/data.frame)
##  $ OTUID               : chr [1:3276] "HAP001" "HAP001" "HAP001" "HAP001" ...
##  $ sampleName          : chr [1:3276] "A1_tag01_tag01_4" "A1_tag02_tag02_4" "A1_tag03_tag03_4" "A2_tag04_tag04_4" ...
##  $ readNum             : num [1:3276] 3412 1678 1397 4196 10615 ...
##  $ mtagNum             : num [1:3276] 3052 1557 1283 3162 7350 ...
##  $ soup                : chr [1:3276] "A" "A" "A" "A" ...
##  $ exp                 : chr [1:3276] "within-spp" "within-spp" "within-spp" "within-spp" ...
##  $ soupRep             : num [1:3276] 1 1 1 2 2 2 3 3 3 1 ...
##  $ PCR                 : chr [1:3276] "tag01_tag01" "tag02_tag02" "tag03_tag03" "tag04_tag04" ...
##  $ lib                 : num [1:3276] 4 4 4 4 4 4 4 4 4 4 ...
##  $ pcrRep              : num [1:3276] 1 2 3 1 2 3 1 2 3 1 ...
##  $ readSpike           : num [1:3276] 8863 3740 4091 8055 20218 ...
##  $ mtagSpike           : num [1:3276] 6642 3199 3441 4967 9843 ...
##  $ readNumSpikeCorr    : num [1:3276] 1439 1678 1277 1948 1963 ...
##  $ mtagNumSpikeCorr    : num [1:3276] 1381 1462 1120 1913 2244 ...
##  $ inputCOINum         : num [1:3276] 100129875 100129875 100129875 100129875 100129875 ...
##  $ inputGDNA           : num [1:3276] 61 61 61 61 61 61 61 61 61 48.8 ...
##  $ log.readNum         : num [1:3276] 8.14 7.43 7.24 8.34 9.27 ...
##  $ log.inputCOINum     : num [1:3276] 18.4 18.4 18.4 18.4 18.4 ...
##  $ log.mtagNum         : num [1:3276] 8.02 7.35 7.16 8.06 8.9 ...
##  $ log.inputGDNA       : num [1:3276] 4.13 4.13 4.13 4.13 4.13 ...
##  $ log.mtagNumSpikeCorr: num [1:3276] 7.23 7.29 7.02 7.56 7.72 ...
##  $ log.readNumSpikeCorr: num [1:3276] 7.27 7.43 7.15 7.57 7.58 ...
##  - attr(*, "spec")=
##   .. cols(
##   ..   OTUID = col_character(),
##   ..   sampleName = col_character(),
##   ..   readNum = col_double(),
##   ..   mtagNum = col_double(),
##   ..   soup = col_character(),
##   ..   exp = col_character(),
##   ..   soupRep = col_double(),
##   ..   PCR = col_character(),
##   ..   lib = col_double(),
##   ..   pcrRep = col_double(),
##   ..   readSpike = col_double(),
##   ..   mtagSpike = col_double(),
##   ..   readNumSpikeCorr = col_double(),
##   ..   mtagNumSpikeCorr = col_double(),
##   ..   inputCOINum = col_double(),
##   ..   inputGDNA = col_double()
##   .. )
```

mixed effects models

Figure 4A

```
lmer_coi_read1 <- lmer(log.inputCOINum ~ log.readNum +
  (log.readNum | OTUID) +
  (1 | soupRep / pcrRep), data = mydata, REML = F)
```

```
## boundary (singular) fit: see help('isSingular')
```

```
# singular fit
lmer_coi_read2 <- lmer(log.inputCOINum ~ log.readNum +
  (1 | OTUID) +
  (1 | soupRep / pcrRep), data = mydata, REML = F)
```

```
## boundary (singular) fit: see help('isSingular')
```

```
# singular fit
lmer_coi_read3 <- lmer(log.inputCOINum ~ log.readNum +
  (1 | soupRep / pcrRep), data = mydata, REML = F)
```

```
## boundary (singular) fit: see help('isSingular')
```

```
# singular fit
AIC(lmer_coi_read1, lmer_coi_read2, lmer_coi_read3)
```

```
##                df      AIC
## lmer_coi_read1  8 4067.218
## lmer_coi_read2  6 4113.070
## lmer_coi_read3  5 9223.563
```

```
# all the models fit very poorly (not a surprise given that the predictor is not spike-corrected). Of these poor models, choose lmer_coi_read1 to carry out the fixed-effect test


# test fixed effect
lmer_coi_read4 <- lmer(log.inputCOINum ~ 1 +
  (log.readNum | OTUID) +
  (1 | soupRep / pcrRep), data = mydata, REML = F)
```

```
## boundary (singular) fit: see help('isSingular')
```

```
# singular fit
AIC(lmer_coi_read1, lmer_coi_read4) # removal of fixed effect increases AIC
```

```
##                df      AIC
## lmer_coi_read1  8 4067.218
## lmer_coi_read4  7 4095.451
```

```
anova(lmer_coi_read1, lmer_coi_read4) # keep fixed effect
```

```
## Data: mydata
## Models:
## lmer_coi_read4: log.inputCOINum ~ 1 + (log.readNum | OTUID) + (1 | soupRep/pcrRep)
## lmer_coi_read1: log.inputCOINum ~ log.readNum + (log.readNum | OTUID) + (1 | soupRep/pcrRep)
##                npar    AIC    BIC  logLik deviance  Chisq Df    Pr(>Chisq)    
## lmer_coi_read4    7 4095.5 4137.8 -2040.7   4081.5                            
## lmer_coi_read1    8 4067.2 4115.7 -2025.6   4051.2 30.233  1 0.00000003832 ***
## ---
## Signif. codes:  0 '***' 0.001 '**' 0.01 '*' 0.05 '.' 0.1 ' ' 1
```

```
# estimate parameters with lmer_coi_read1
lmer_coi_read5 <- lmer(log.inputCOINum ~ log.readNum +
  (log.readNum | OTUID) +
  (1 | soupRep / pcrRep), data = mydata, REML = TRUE)
```

```
## boundary (singular) fit: see help('isSingular')
```

```
# singular fit
summary(lmer_coi_read5)
```

```
## Linear mixed model fit by REML ['lmerMod']
## Formula: log.inputCOINum ~ log.readNum + (log.readNum | OTUID) + (1 |  
##     soupRep/pcrRep)
##    Data: mydata
## 
## REML criterion at convergence: 4059.6
## 
## Scaled residuals: 
##      Min       1Q   Median       3Q      Max 
## -1.94255 -0.80640 -0.02837  0.92313  1.99929 
## 
## Random effects:
##  Groups         Name        Variance Std.Dev. Corr 
##  OTUID          (Intercept) 1.332730 1.15444       
##                 log.readNum 0.008726 0.09341  -0.50
##  pcrRep:soupRep (Intercept) 0.000000 0.00000       
##  soupRep        (Intercept) 0.000000 0.00000       
##  Residual                   0.190370 0.43631       
## Number of obs: 3150, groups:  OTUID, 50; pcrRep:soupRep, 9; soupRep, 3
## 
## Fixed effects:
##             Estimate Std. Error t value
## (Intercept) 15.67884    0.18015  87.034
## log.readNum  0.10655    0.01714   6.216
## 
## Correlation of Fixed Effects:
##             (Intr)
## log.readNum -0.609
## optimizer (nloptwrap) convergence code: 0 (OK)
## boundary (singular) fit: see help('isSingular')
```

```
r.squaredGLMM(lmer_coi_read5)
```

```
## Warning: 'r.squaredGLMM' now calculates a revised statistic. See the help page.
```

```
## Warning in pcrRep:soupRep: numerical expression has 3150 elements: only the
## first used

## Warning in pcrRep:soupRep: numerical expression has 3150 elements: only the
## first used
```

```
##             R2m       R2c
## [1,] 0.04196802 0.8528916
```

```
#             R2m       R2c
# [1,] 0.04196802 0.8528916
# R2m = marginal R-squared (variance explained by fixed effects only)
# R2c = conditional R-squared (variance explained by random + fixed effects)

sresid <- resid(lmer_coi_read5, type = "pearson") # Extract the standardised residuals, check underline assumption of normally distibuted
hist(sresid)
```

```
plot(lmer_coi_read5) # note that the model fit is very poor, so the residuals plots will also be poor
```

```
pred.mm2 <- ggpredict(lmer_coi_read5, terms = c("log.readNum"))

# Plot the predictions
(p2 <- ggplot() +
  geom_line(
    data = pred.mm2, aes(x = x, y = predicted),
    colour = "black", size = 1
  ) + # slope
  geom_ribbon(
    data = pred.mm2, aes(
      x = x, ymin = predicted - std.error,
      ymax = predicted + std.error
    ),
    fill = "lightgrey", alpha = 0.5
  ) +
  geom_point(data = mydata, aes(
    x = log.readNum, y = log.inputCOINum,
    colour = OTUID
  ), size = 0.8) +
  geom_smooth(
    data = mydata, aes(
      x = log.readNum, y = log.inputCOINum,
      group = OTUID
    ),
    method = "lm", se = FALSE, size = 0.3, colour = "black", alpha = 0.5,
    linetype = 1
  ) +
  labs(x = "ln(non-spike-corrected OTU size)", y = "ln(input COI copy number)") +
  theme_cowplot() +
  theme(legend.position = "none"))
```

```
## `geom_smooth()` using formula 'y ~ x'
```

```
## Warning: Removed 126 rows containing non-finite values (stat_smooth).
```

```
## Warning: Removed 126 rows containing missing values (geom_point).
```

Figure 4B

```
# test random effects models
# with random slope
lmer_coi_readSpikeCorr1 <- lmer(log.inputCOINum ~ log.readNumSpikeCorr +
  (log.readNumSpikeCorr | OTUID) +
  (1 | soupRep / pcrRep), data = mydata, REML = FALSE)
summary(lmer_coi_readSpikeCorr1)
```

```
## Linear mixed model fit by maximum likelihood  ['lmerMod']
## Formula: log.inputCOINum ~ log.readNumSpikeCorr + (log.readNumSpikeCorr |  
##     OTUID) + (1 | soupRep/pcrRep)
##    Data: mydata
## 
##      AIC      BIC   logLik deviance df.resid 
##    962.0   1010.5   -473.0    946.0     3142 
## 
## Scaled residuals: 
##     Min      1Q  Median      3Q     Max 
## -3.6244 -0.6908 -0.0009  0.6198  3.6445 
## 
## Random effects:
##  Groups         Name                 Variance  Std.Dev. Corr 
##  OTUID          (Intercept)          1.1492493 1.07203       
##                 log.readNumSpikeCorr 0.0073741 0.08587  -0.55
##  pcrRep:soupRep (Intercept)          0.0027171 0.05213       
##  soupRep        (Intercept)          0.0009887 0.03144       
##  Residual                            0.0685267 0.26178       
## Number of obs: 3150, groups:  OTUID, 50; pcrRep:soupRep, 9; soupRep, 3
## 
## Fixed effects:
##                      Estimate Std. Error t value
## (Intercept)          14.50435    0.15590   93.04
## log.readNumSpikeCorr  0.39441    0.01321   29.85
## 
## Correlation of Fixed Effects:
##             (Intr)
## lg.rdNmSpkC -0.549
```

```
plot(lmer_coi_readSpikeCorr1)
```

```
# without random slope
lmer_coi_readSpikeCorr2 <- lmer(log.inputCOINum ~ log.readNumSpikeCorr +
  (1 | OTUID) +
  (1 | soupRep / pcrRep), data = mydata, REML = FALSE)
summary(lmer_coi_readSpikeCorr2)
```

```
## Linear mixed model fit by maximum likelihood  ['lmerMod']
## Formula: 
## log.inputCOINum ~ log.readNumSpikeCorr + (1 | OTUID) + (1 | soupRep/pcrRep)
##    Data: mydata
## 
##      AIC      BIC   logLik deviance df.resid 
##   1439.6   1475.9   -713.8   1427.6     3144 
## 
## Scaled residuals: 
##     Min      1Q  Median      3Q     Max 
## -3.7168 -0.7480  0.0503  0.6108  4.4756 
## 
## Random effects:
##  Groups         Name        Variance  Std.Dev.
##  OTUID          (Intercept) 0.8670657 0.93116 
##  pcrRep:soupRep (Intercept) 0.0020181 0.04492 
##  soupRep        (Intercept) 0.0008191 0.02862 
##  Residual                   0.0825792 0.28737 
## Number of obs: 3150, groups:  OTUID, 50; pcrRep:soupRep, 9; soupRep, 3
## 
## Fixed effects:
##                       Estimate Std. Error t value
## (Intercept)          14.780431   0.135812  108.83
## log.readNumSpikeCorr  0.347339   0.005174   67.14
## 
## Correlation of Fixed Effects:
##             (Intr)
## lg.rdNmSpkC -0.177
```

```
plot(lmer_coi_readSpikeCorr2)
```

```
# without random OTUID 
lmer_coi_readSpikeCorr3 <- lmer(log.inputCOINum ~ log.readNumSpikeCorr +
   (1 | soupRep/pcrRep), data = mydata, REML = FALSE)
```

```
## boundary (singular) fit: see help('isSingular')
```

```
# singular fit
summary(lmer_coi_readSpikeCorr3)
```

```
## Linear mixed model fit by maximum likelihood  ['lmerMod']
## Formula: log.inputCOINum ~ log.readNumSpikeCorr + (1 | soupRep/pcrRep)
##    Data: mydata
## 
##      AIC      BIC   logLik deviance df.resid 
##   8788.2   8818.5  -4389.1   8778.2     3145 
## 
## Scaled residuals: 
##      Min       1Q   Median       3Q      Max 
## -2.58952 -0.64926  0.08377  0.67831  3.07467 
## 
## Random effects:
##  Groups         Name        Variance   Std.Dev.
##  pcrRep:soupRep (Intercept) 0.00000000 0.000000
##  soupRep        (Intercept) 0.00005246 0.007243
##  Residual                   0.95010196 0.974732
## Number of obs: 3150, groups:  pcrRep:soupRep, 9; soupRep, 3
## 
## Fixed effects:
##                       Estimate Std. Error t value
## (Intercept)          14.862407   0.042448  350.13
## log.readNumSpikeCorr  0.329729   0.008272   39.86
## 
## Correlation of Fixed Effects:
##             (Intr)
## lg.rdNmSpkC -0.907
## optimizer (nloptwrap) convergence code: 0 (OK)
## boundary (singular) fit: see help('isSingular')
```

```
plot(lmer_coi_readSpikeCorr3)
```

```
AIC(lmer_coi_readSpikeCorr1, lmer_coi_readSpikeCorr2, lmer_coi_readSpikeCorr3) # lmer_coi_readSpikeCorr1 has lower AIC, keep random intercept and slope, keep OTUID, soupRep, pcrRep
```

```
##                         df       AIC
## lmer_coi_readSpikeCorr1  8  962.0384
## lmer_coi_readSpikeCorr2  6 1439.5796
## lmer_coi_readSpikeCorr3  5 8788.2460
```

```
# test fixed effects
lmer_coi_readSpikeCorr4 <- lmer(log.inputCOINum ~ 1 +
  (log.readNumSpikeCorr | OTUID) +
  (1 | soupRep / pcrRep), data = mydata, REML = FALSE)
AIC(lmer_coi_readSpikeCorr1, lmer_coi_readSpikeCorr4) # lmer_coi_readSpikeCorr1
```

```
##                         df       AIC
## lmer_coi_readSpikeCorr1  8  962.0384
## lmer_coi_readSpikeCorr4  7 1109.2054
```

```
anova(lmer_coi_readSpikeCorr1, lmer_coi_readSpikeCorr4) # significantly different. keep fixed effect
```

```
## Data: mydata
## Models:
## lmer_coi_readSpikeCorr4: log.inputCOINum ~ 1 + (log.readNumSpikeCorr | OTUID) + (1 | soupRep/pcrRep)
## lmer_coi_readSpikeCorr1: log.inputCOINum ~ log.readNumSpikeCorr + (log.readNumSpikeCorr | OTUID) + (1 | soupRep/pcrRep)
##                         npar     AIC    BIC  logLik deviance  Chisq Df
## lmer_coi_readSpikeCorr4    7 1109.21 1151.6 -547.60  1095.21          
## lmer_coi_readSpikeCorr1    8  962.04 1010.5 -473.02   946.04 149.17  1
##                                    Pr(>Chisq)    
## lmer_coi_readSpikeCorr4                          
## lmer_coi_readSpikeCorr1 < 0.00000000000000022 ***
## ---
## Signif. codes:  0 '***' 0.001 '**' 0.01 '*' 0.05 '.' 0.1 ' ' 1
```

```
# parameter estimates
lmer_coi_readSpikeCorr5 <- lmer(log.inputCOINum ~ log.readNumSpikeCorr +
  (log.readNumSpikeCorr | OTUID) +
  (1 | soupRep / pcrRep), data = mydata, REML = TRUE)
summary(lmer_coi_readSpikeCorr5)
```

```
## Linear mixed model fit by REML ['lmerMod']
## Formula: log.inputCOINum ~ log.readNumSpikeCorr + (log.readNumSpikeCorr |  
##     OTUID) + (1 | soupRep/pcrRep)
##    Data: mydata
## 
## REML criterion at convergence: 955.1
## 
## Scaled residuals: 
##     Min      1Q  Median      3Q     Max 
## -3.6294 -0.6903 -0.0012  0.6196  3.6433 
## 
## Random effects:
##  Groups         Name                 Variance Std.Dev. Corr 
##  OTUID          (Intercept)          1.172418 1.08278       
##                 log.readNumSpikeCorr 0.007539 0.08683  -0.55
##  pcrRep:soupRep (Intercept)          0.002718 0.05214       
##  soupRep        (Intercept)          0.001027 0.03204       
##  Residual                            0.068528 0.26178       
## Number of obs: 3150, groups:  OTUID, 50; pcrRep:soupRep, 9; soupRep, 3
## 
## Fixed effects:
##                      Estimate Std. Error t value
## (Intercept)          14.50390    0.15742   92.13
## log.readNumSpikeCorr  0.39446    0.01334   29.57
## 
## Correlation of Fixed Effects:
##             (Intr)
## lg.rdNmSpkC -0.549
```

```
r.squaredGLMM(lmer_coi_readSpikeCorr5)
```

```
## Warning in pcrRep:soupRep: numerical expression has 3150 elements: only the
## first used

## Warning in pcrRep:soupRep: numerical expression has 3150 elements: only the
## first used
```

```
##            R2m       R2c
## [1,] 0.4160562 0.9584505
```

```
#            R2m       R2c
# [1,] 0.4160562 0.9584505
# R2m = marginal R-squared (variance explained by fixed effects only)
# R2c = conditional R-squared (variance explained by random + fixed effects)

sresid <- resid(lmer_coi_readSpikeCorr5, type = "pearson")
hist(sresid)
```

```
plot(lmer_coi_readSpikeCorr5)
```

```
plot(sresid ~ lmer_coi_readSpikeCorr5@frame[["log.readNumSpikeCorr"]])
```

```
pred.mm1 <- ggpredict(lmer_coi_readSpikeCorr5, terms = c("log.readNumSpikeCorr"))

(p1 <- ggplot() +
  geom_line(
    data = pred.mm1, aes(x = x, y = predicted),
    colour = "black", size = 1
  ) +
  geom_ribbon(
    data = pred.mm1, aes(
      x = x, ymin = predicted - std.error,
      ymax = predicted + std.error
    ),
    fill = "lightgrey", alpha = 0.3
  ) +
  geom_point(data = mydata, aes(
    x = log.readNumSpikeCorr, y = log.inputCOINum,
    colour = OTUID
  ), size = 0.8) +
  geom_smooth(
    data = mydata, aes(
      x = log.readNumSpikeCorr, y = log.inputCOINum,
      group = OTUID
    ),
    method = "lm", se = FALSE, size = 0.3, colour = "black", alpha = 0.5,
    linetype = 1
  ) +
  labs(x = "ln(spike-corrected OTU size)", y = "ln(input COI copy number)") +
  theme_cowplot() +
  theme(legend.position = "none"))
```

```
## `geom_smooth()` using formula 'y ~ x'
```

```
## Warning: Removed 126 rows containing non-finite values (stat_smooth).
```

```
## Warning: Removed 126 rows containing missing values (geom_point).
```

Figure 4C

```
lmer_gDNA_read1 <- lmer(log.inputGDNA ~ log.readNum +
  (log.readNum | OTUID) +
  (1 | soupRep / pcrRep), data = mydata, REML = F)
```

```
## boundary (singular) fit: see help('isSingular')
```

```
# singular fit
lmer_gDNA_read2 <- lmer(log.inputGDNA ~ log.readNum +
  (1 | OTUID) +
  (1 | soupRep / pcrRep), data = mydata, REML = F)
```

```
## boundary (singular) fit: see help('isSingular')
```

```
# singular fit
lmer_gDNA_read3 <- lmer(log.inputGDNA ~ log.readNum +
  (1 | soupRep / pcrRep), data = mydata, REML = F)
```

```
## boundary (singular) fit: see help('isSingular')
```

```
# singular fits for all models

AIC(lmer_gDNA_read1, lmer_gDNA_read2, lmer_gDNA_read3) # lmer_gDNA_read3
```

```
##                 df      AIC
## lmer_gDNA_read1  8 3806.395
## lmer_gDNA_read2  6 3784.565
## lmer_gDNA_read3  5 3782.565
```

```
# all the models are poor (since we are using non-spike-corrected readNum)

# test fixed effect
lmer_gDNA_read4 <- lmer(log.inputGDNA ~ 1 +
  (1 | soupRep / pcrRep), data = mydata, REML = F)
```

```
## boundary (singular) fit: see help('isSingular')
```

```
# singular fit
AIC(lmer_gDNA_read3, lmer_gDNA_read4) # keep fixed effect
```

```
##                 df      AIC
## lmer_gDNA_read3  5 3782.565
## lmer_gDNA_read4  4 3796.578
```

```
anova(lmer_gDNA_read3, lmer_gDNA_read4)
```

```
## Data: mydata
## Models:
## lmer_gDNA_read4: log.inputGDNA ~ 1 + (1 | soupRep/pcrRep)
## lmer_gDNA_read3: log.inputGDNA ~ log.readNum + (1 | soupRep/pcrRep)
##                 npar    AIC  BIC  logLik deviance  Chisq Df Pr(>Chisq)    
## lmer_gDNA_read4    4 3796.6 3821 -1894.3   3788.6                         
## lmer_gDNA_read3    5 3782.6 3813 -1886.3   3772.6 16.013  1  0.0000629 ***
## ---
## Signif. codes:  0 '***' 0.001 '**' 0.01 '*' 0.05 '.' 0.1 ' ' 1
```

```
######## final model###########
lmer_gDNA_read5 <- lmer(log.inputGDNA ~ log.readNum +
  (1 | soupRep / pcrRep), data = mydata, REML = T)
```

```
## boundary (singular) fit: see help('isSingular')
```

```
# singular fit
r.squaredGLMM(lmer_gDNA_read5) # 0.005 0.005
```

```
## Warning in pcrRep:soupRep: numerical expression has 3276 elements: only the
## first used

## Warning in pcrRep:soupRep: numerical expression has 3276 elements: only the
## first used
```

```
##              R2m         R2c
## [1,] 0.004874685 0.004874685
```

```
#              R2m         R2c
# [1,] 0.004874685 0.004874685

sresid <- resid(lmer_gDNA_read5, type = "pearson") # Extract the standardised residuals, check underline assumption of normally distibuted
hist(sresid)
```

```
plot(sresid ~ mydata$log.readNum) # poor model means weird residuals
```

```
pred.mm4 <- ggpredict(lmer_gDNA_read5, terms = c("log.readNum"))

# Plot the predictions
(p4 <- ggplot() +
  geom_line(
    data = pred.mm4, aes(x = x, y = predicted),
    colour = "black", size = 1
  ) + # slope
  geom_ribbon(
    data = pred.mm4, aes(
      x = x, ymin = predicted - std.error,
      ymax = predicted + std.error
    ),
    fill = "lightgrey", alpha = 0.5
  ) +
  geom_point(data = mydata, aes(
    x = log.readNum, y = log.inputGDNA,
    colour = OTUID
  ), size = 0.8) +
  geom_smooth(
    data = mydata, aes(
      x = log.readNum, y = log.inputGDNA,
      group = OTUID
    ),
    method = "lm", se = FALSE, size = 0.3, colour = "black", alpha = 0.5,
    linetype = 1
  ) +
  ylim(2.3, 4.7) +
  labs(x = "ln(non-spike-corrected OTU size)", y = "ln(Input Genomic DNA)") +
  theme_cowplot() +
  theme(legend.position = "none"))
```

```
## `geom_smooth()` using formula 'y ~ x'
```

Figure 4D

```
lmer_gDNA_readSpikeCorr1 <- lmer(log.inputGDNA ~ log.readNumSpikeCorr +
   (log.readNumSpikeCorr | OTUID) +
   (1 | soupRep / pcrRep), data = mydata, REML = FALSE)

lmer_gDNA_readSpikeCorr2 <- lmer(log.inputGDNA ~ log.readNumSpikeCorr +
   (1 | OTUID) +
   (1 | soupRep / pcrRep), data = mydata, REML = FALSE)

lmer_gDNA_readSpikeCorr3 <- lmer(log.inputGDNA ~ log.readNumSpikeCorr +
   (1 | soupRep / pcrRep), data = mydata, REML = FALSE)
```

```
## boundary (singular) fit: see help('isSingular')
```

```
# singular fit

AIC(lmer_gDNA_readSpikeCorr1, lmer_gDNA_readSpikeCorr2, lmer_gDNA_readSpikeCorr3) # lmer_gDNA_readSpikeCorr1 has lower AIC, use random intercept and slope, keep OTUID, soupRep, pcrRep
```

```
##                          df      AIC
## lmer_gDNA_readSpikeCorr1  8  706.814
## lmer_gDNA_readSpikeCorr2  6 1186.054
## lmer_gDNA_readSpikeCorr3  5 3317.231
```

```
# test fixed effects
lmer_gDNA_readSpikeCorr4 <- lmer(log.inputGDNA ~ 1 +
   (log.readNumSpikeCorr | OTUID) +
   (1 | soupRep / pcrRep), data = mydata, REML = F)

AIC(lmer_gDNA_readSpikeCorr1, lmer_gDNA_readSpikeCorr4) # choose lmer_gDNA_readSpikeCorr1
```

```
##                          df      AIC
## lmer_gDNA_readSpikeCorr1  8 706.8140
## lmer_gDNA_readSpikeCorr4  7 859.4424
```

```
anova(lmer_gDNA_readSpikeCorr1, lmer_gDNA_readSpikeCorr4) # significantly different. keep fixed effect
```

```
## Data: mydata
## Models:
## lmer_gDNA_readSpikeCorr4: log.inputGDNA ~ 1 + (log.readNumSpikeCorr | OTUID) + (1 | soupRep/pcrRep)
## lmer_gDNA_readSpikeCorr1: log.inputGDNA ~ log.readNumSpikeCorr + (log.readNumSpikeCorr | OTUID) + (1 | soupRep/pcrRep)
##                          npar    AIC    BIC  logLik deviance  Chisq Df
## lmer_gDNA_readSpikeCorr4    7 859.44 902.10 -422.72   845.44          
## lmer_gDNA_readSpikeCorr1    8 706.81 755.57 -345.41   690.81 154.63  1
##                                     Pr(>Chisq)    
## lmer_gDNA_readSpikeCorr4                          
## lmer_gDNA_readSpikeCorr1 < 0.00000000000000022 ***
## ---
## Signif. codes:  0 '***' 0.001 '**' 0.01 '*' 0.05 '.' 0.1 ' ' 1
```

```
# fit model
lmer_gDNA_readSpikeCorr5 <- lmer(log.inputGDNA ~ log.readNumSpikeCorr +
   (log.readNumSpikeCorr | OTUID) +
   (1 | soupRep / pcrRep), data = mydata, REML = TRUE)
summary(lmer_gDNA_readSpikeCorr5)
```

```
## Linear mixed model fit by REML ['lmerMod']
## Formula: log.inputGDNA ~ log.readNumSpikeCorr + (log.readNumSpikeCorr |  
##     OTUID) + (1 | soupRep/pcrRep)
##    Data: mydata
## 
## REML criterion at convergence: 700.7
## 
## Scaled residuals: 
##     Min      1Q  Median      3Q     Max 
## -3.6412 -0.6920 -0.0103  0.6215  3.6513 
## 
## Random effects:
##  Groups         Name                 Variance Std.Dev. Corr 
##  OTUID          (Intercept)          0.748512 0.86517       
##                 log.readNumSpikeCorr 0.006914 0.08315  -0.68
##  pcrRep:soupRep (Intercept)          0.002447 0.04947       
##  soupRep        (Intercept)          0.001025 0.03202       
##  Residual                            0.063288 0.25157       
## Number of obs: 3276, groups:  OTUID, 52; pcrRep:soupRep, 9; soupRep, 3
## 
## Fixed effects:
##                      Estimate Std. Error t value
## (Intercept)           1.66196    0.12494   13.30
## log.readNumSpikeCorr  0.37640    0.01251   30.08
## 
## Correlation of Fixed Effects:
##             (Intr)
## lg.rdNmSpkC -0.672
```

```
r.squaredGLMM(lmer_gDNA_readSpikeCorr5) # 0.531 0.96
```

```
## Warning in pcrRep:soupRep: numerical expression has 3276 elements: only the
## first used

## Warning in pcrRep:soupRep: numerical expression has 3276 elements: only the
## first used
```

```
##            R2m       R2c
## [1,] 0.5314312 0.9450565
```

```
#            R2m       R2c
# [1,] 0.5314312 0.9450565

sresid <- resid(lmer_gDNA_readSpikeCorr5, type = "pearson")
hist(sresid)
```

```
plot(lmer_gDNA_readSpikeCorr5)
```

```
plot(sresid ~ lmer_gDNA_readSpikeCorr5@frame[["log.readNumSpikeCorr"]]) # wider residuals with small OTUs
```

```
pred.mm3 <- ggpredict(lmer_gDNA_readSpikeCorr5, terms = c("log.readNumSpikeCorr"))

(p3 <- ggplot() +
  geom_line(
    data = pred.mm3, aes(x = x, y = predicted),
    colour = "black", size = 1
  ) + 
  geom_ribbon(
    data = pred.mm3, aes(
      x = x, ymin = predicted - std.error,
      ymax = predicted + std.error
    ),
    fill = "lightgrey", alpha = 0.5
  ) +
  geom_point(data = mydata, aes(
    x = log.readNumSpikeCorr, y = log.inputGDNA,
    colour = OTUID
  ), size = 0.8) +
  geom_smooth(
    data = mydata, aes(
      x = log.readNumSpikeCorr, y = log.inputGDNA,
      group = OTUID
    ),
    method = "lm", se = FALSE, size = 0.3, colour = "black", alpha = 0.5,
    linetype = 1
  ) +
  ylim(2.3, 4.7) +
  labs(x = "ln(spike-corrected OTU size)", y = "ln(input genomic DNA)") +
  theme_cowplot() +
  theme(legend.position = "none"))
```

```
## `geom_smooth()` using formula 'y ~ x'
```

```
## Warning: Removed 3 row(s) containing missing values (geom_path).
```

```
## Warning: Removed 1 rows containing missing values (geom_smooth).
```

Figure 4. Recovery of within-species abundance change in COI copy
number and in genomic DNA concentration in the mock-soup experiment.

```
(p2 + p1) / (p4 + p3)
```

```
## `geom_smooth()` using formula 'y ~ x'
```

```
## Warning: Removed 126 rows containing non-finite values (stat_smooth).
```

```
## Warning: Removed 126 rows containing missing values (geom_point).
```

```
## `geom_smooth()` using formula 'y ~ x'
```

```
## Warning: Removed 126 rows containing non-finite values (stat_smooth).
## Removed 126 rows containing missing values (geom_point).
```

```
## `geom_smooth()` using formula 'y ~ x'
## `geom_smooth()` using formula 'y ~ x'
```

```
## Warning: Removed 3 row(s) containing missing values (geom_path).
```

```
## Warning: Removed 1 rows containing missing values (geom_smooth).
```

Supplementary information S3. Using UMI number instead of spike-ins
to correct pipeline noise.

```
lmer_gDNA_UMI1 <- lmer(log.inputGDNA ~ log.mtagNum +
  (log.mtagNum | OTUID) +
  (1 | soupRep / pcrRep), data = mydata, REML = F)
```

```
## boundary (singular) fit: see help('isSingular')
```

```
# singular fit
lmer_gDNA_UMI2 <- lmer(log.inputGDNA ~ log.mtagNum +
  (1 | OTUID) +
  (1 | soupRep / pcrRep), data = mydata, REML = F)
```

```
## boundary (singular) fit: see help('isSingular')
```

```
# singular fit
lmer_gDNA_UMI3 <- lmer(log.inputGDNA ~ log.mtagNum +
  (1 | soupRep / pcrRep), data = mydata, REML = F)
```

```
## boundary (singular) fit: see help('isSingular')
```

```
# singular fit

AIC(lmer_gDNA_UMI1, lmer_gDNA_UMI2, lmer_gDNA_UMI3) # lmer_gDNA_UMI1
```

```
##                df      AIC
## lmer_gDNA_UMI1  8 3774.613
## lmer_gDNA_UMI2  6 3783.022
## lmer_gDNA_UMI3  5 3781.022
```

```
# fixed effect
lmer_gDNA_UMI4 <- lmer(log.inputGDNA ~ 1 +
  (log.mtagNum | OTUID) +
  (1 | soupRep / pcrRep), data = mydata, REML = F)
```

```
## boundary (singular) fit: see help('isSingular')
```

```
AIC(lmer_gDNA_UMI1, lmer_gDNA_UMI4) # lmer_gDNA_UMI1
```

```
##                df      AIC
## lmer_gDNA_UMI1  8 3774.613
## lmer_gDNA_UMI4  7 3790.226
```

```
anova(lmer_gDNA_UMI1, lmer_gDNA_UMI4) # lmer_gDNA_UMI1
```

```
## Data: mydata
## Models:
## lmer_gDNA_UMI4: log.inputGDNA ~ 1 + (log.mtagNum | OTUID) + (1 | soupRep/pcrRep)
## lmer_gDNA_UMI1: log.inputGDNA ~ log.mtagNum + (log.mtagNum | OTUID) + (1 | soupRep/pcrRep)
##                npar    AIC    BIC  logLik deviance  Chisq Df Pr(>Chisq)    
## lmer_gDNA_UMI4    7 3790.2 3832.9 -1888.1   3776.2                         
## lmer_gDNA_UMI1    8 3774.6 3823.4 -1879.3   3758.6 17.613  1 0.00002708 ***
## ---
## Signif. codes:  0 '***' 0.001 '**' 0.01 '*' 0.05 '.' 0.1 ' ' 1
```

```
lmer_gDNA_UMI5 <- lmer(log.inputGDNA ~ log.mtagNum +
  (log.mtagNum | OTUID) +
  (1 | soupRep / pcrRep), data = mydata, REML = T)
```

```
## boundary (singular) fit: see help('isSingular')
```

```
r.squaredGLMM(lmer_gDNA_UMI5) # MuMIn package 0.015 0.033
```

```
## Warning in pcrRep:soupRep: numerical expression has 3276 elements: only the
## first used

## Warning in pcrRep:soupRep: numerical expression has 3276 elements: only the
## first used
```

```
##             R2m        R2c
## [1,] 0.01546805 0.03252212
```

```
# R2m = marginal R-squared (variance explained by fixed effects only)
# R2c = conditional R-squared (variance explained by random + fixed effects)

sresid <- resid(lmer_gDNA_UMI5, type = "pearson") # Extract the standardised residuals, check underline assumption of normally distibuted
hist(sresid)
```

```
plot(lmer_gDNA_UMI5) # gives a heteroscedasticity plot
```

```
plot(sresid ~ mydata$log.mtagNum)
```

```
pred.mm5 <- ggpredict(lmer_gDNA_UMI5, terms = c("log.mtagNum"))

# Plot the predictions
(p5 <- ggplot() +
  geom_line(data = pred.mm5, aes(x = x, y = predicted),
            colour = "black", size = 1) + # slope
  geom_ribbon(
    data = pred.mm5, aes(
      x = x, ymin = predicted - std.error,
      ymax = predicted + std.error
    ),
    fill = "lightgrey", alpha = 0.5
  ) +
  geom_point(data = mydata, aes(
    x = log.mtagNum, y = log.inputGDNA,
    colour = OTUID
  ), size = 0.8) +
  geom_smooth(
    data = mydata, aes(
      x = log.mtagNum, y = log.inputGDNA,
      group = OTUID
    ),
    method = "lm", se = FALSE, size = 0.3, colour = "black", alpha = 0.5,
    linetype = 1
  ) +
  labs(y = "ln(Input Genomic DNA)", x = "ln(non-spike-corrected UMI number)") +
  theme_cowplot() +
  theme(legend.position = "none"))
```

```
## `geom_smooth()` using formula 'y ~ x'
```

```
lmer_gDNA_UMIspikeCorr1 <- lmer(log.inputGDNA ~ log.mtagNumSpikeCorr +
  (log.mtagNumSpikeCorr | OTUID) +
  (1 | soupRep / pcrRep), data = mydata, REML = F)
```

```
## Warning in checkConv(attr(opt, "derivs"), opt$par, ctrl = control$checkConv, :
## Model failed to converge with max|grad| = 0.00280686 (tol = 0.002, component 1)
```

```
lmer_gDNA_UMIspikeCorr2 <- lmer(log.inputGDNA ~ log.mtagNumSpikeCorr +
  (1 | OTUID) +
  (1 | soupRep / pcrRep), data = mydata, REML = F)
lmer_gDNA_UMIspikeCorr3 <- lmer(log.inputGDNA ~ log.mtagNumSpikeCorr +
  (1 | soupRep / pcrRep), data = mydata, REML = F)
```

```
## boundary (singular) fit: see help('isSingular')
```

```
# singular fit

AIC(lmer_gDNA_UMIspikeCorr1, lmer_gDNA_UMIspikeCorr2, lmer_gDNA_UMIspikeCorr3) # lmer_gDNA_UMIspikeCorr1
```

```
##                         df      AIC
## lmer_gDNA_UMIspikeCorr1  8 1624.234
## lmer_gDNA_UMIspikeCorr2  6 2134.862
## lmer_gDNA_UMIspikeCorr3  5 3517.856
```

```
# fixed effect
lmer_gDNA_UMIspikeCorr4 <- lmer(log.inputGDNA ~ 1 +
  (log.mtagNumSpikeCorr | OTUID) +
  (1 | soupRep / pcrRep), data = mydata, REML = F)
```

```
## Warning in checkConv(attr(opt, "derivs"), opt$par, ctrl = control$checkConv, :
## Model failed to converge with max|grad| = 0.00335872 (tol = 0.002, component 1)
```

```
# failure to converge
AIC(lmer_gDNA_UMIspikeCorr1, lmer_gDNA_UMIspikeCorr4) # lmer_gDNA_UMIspikeCorr1
```

```
##                         df      AIC
## lmer_gDNA_UMIspikeCorr1  8 1624.234
## lmer_gDNA_UMIspikeCorr4  7 1751.368
```

```
# parameter estimate
lmer_gDNA_UMIspikeCorr5 <- lmer(log.inputGDNA ~ log.mtagNumSpikeCorr +
  (log.mtagNumSpikeCorr | OTUID) +
  (1 | soupRep / pcrRep), data = mydata, REML = T)

r.squaredGLMM(lmer_gDNA_UMIspikeCorr5) # 0.52 0.94
```

```
## Warning in pcrRep:soupRep: numerical expression has 3276 elements: only the
## first used
```

```
## Warning in pcrRep:soupRep: numerical expression has 3276 elements: only the
## first used
```

```
##            R2m       R2c
## [1,] 0.5154425 0.9379727
```

```
sresid <- resid(lmer_gDNA_UMIspikeCorr5, type = "pearson")
hist(sresid)
```

```
plot(lmer_gDNA_UMIspikeCorr5)
```

```
plot(sresid ~ mydata$log.mtagNumSpikeCorr)
```

```
pred.mm6 <- ggpredict(lmer_gDNA_UMIspikeCorr5, terms = c("log.mtagNumSpikeCorr"))

(p6 <- ggplot() +
  geom_line(data = pred.mm6, aes(x = x, y = predicted),
            colour = "black", size = 1) + # slope
  geom_ribbon(
    data = pred.mm6, aes(
      x = x, ymin = predicted - std.error,
      ymax = predicted + std.error
    ),
    fill = "lightgrey", alpha = 0.5
  ) +
  geom_point(data = mydata, aes(
    x = log.mtagNumSpikeCorr, y = log.inputGDNA,
    colour = OTUID
  ), size = 0.8) +
  geom_smooth(
    data = mydata, aes(
      x = log.mtagNumSpikeCorr, y = log.inputGDNA,
      group = OTUID
    ),
    method = "lm", se = FALSE, size = 0.3, colour = "black", alpha = 0.5,
    linetype = 1
  ) +
  ylim(2.3, 4.7) + 
  labs(y = "ln(Input Genomic DNA)", x = "ln(spike-corrected UMI number)") +
  theme_cowplot() +
  theme(legend.position = "none"))
```

```
## `geom_smooth()` using formula 'y ~ x'
```

```
## Warning: Removed 4 row(s) containing missing values (geom_path).
```

```
lmer_COI_UMIspike1 <- lmer(log.inputCOINum ~ log.mtagNum +
  (log.mtagNum | OTUID) +
  (1 | soupRep / pcrRep), data = mydata, REML = F)
```

```
## boundary (singular) fit: see help('isSingular')
```

```
# singular fit
lmer_COI_UMIspike2 <- lmer(log.inputCOINum ~ log.mtagNum +
  (1 | OTUID) +
  (1 | soupRep / pcrRep), data = mydata, REML = F)
```

```
## boundary (singular) fit: see help('isSingular')
```

```
# singular fit
lmer_COI_UMIspike3 <- lmer(log.inputCOINum ~ log.mtagNum +
  (1 | soupRep / pcrRep), data = mydata, REML = F)
```

```
## boundary (singular) fit: see help('isSingular')
```

```
# singular fit

AIC(lmer_COI_UMIspike1, lmer_COI_UMIspike2, lmer_COI_UMIspike3) # lmer_gDNA_UMIspikeCorr1
```

```
##                    df      AIC
## lmer_COI_UMIspike1  8 4046.493
## lmer_COI_UMIspike2  6 4099.009
## lmer_COI_UMIspike3  5 9217.905
```

```
# fixed effect
lmer_COI_UMIspike4 <- lmer(log.inputCOINum ~ 1 +
  (1 | OTUID) +
  (1 | soupRep / pcrRep), data = mydata, REML = F)
```

```
## boundary (singular) fit: see help('isSingular')
```

```
# singular fit
AIC(lmer_COI_UMIspike1, lmer_COI_UMIspike4) # lmer_COI_UMIspikeCorr1
```

```
##                    df      AIC
## lmer_COI_UMIspike1  8 4046.493
## lmer_COI_UMIspike4  5 4209.880
```

```
anova(lmer_COI_UMIspike1, lmer_COI_UMIspike4)
```

```
## Data: mydata
## Models:
## lmer_COI_UMIspike4: log.inputCOINum ~ 1 + (1 | OTUID) + (1 | soupRep/pcrRep)
## lmer_COI_UMIspike1: log.inputCOINum ~ log.mtagNum + (log.mtagNum | OTUID) + (1 | soupRep/pcrRep)
##                    npar    AIC    BIC  logLik deviance  Chisq Df
## lmer_COI_UMIspike4    5 4209.9 4240.2 -2099.9   4199.9          
## lmer_COI_UMIspike1    8 4046.5 4094.9 -2015.2   4030.5 169.39  3
##                               Pr(>Chisq)    
## lmer_COI_UMIspike4                          
## lmer_COI_UMIspike1 < 0.00000000000000022 ***
## ---
## Signif. codes:  0 '***' 0.001 '**' 0.01 '*' 0.05 '.' 0.1 ' ' 1
```

```
# parameter estimate
lmer_COI_UMIspike5 <- lmer(log.inputCOINum ~ log.mtagNum +
  (log.mtagNum | OTUID) +
  (1 | soupRep / pcrRep), data = mydata, REML = T)
```

```
## boundary (singular) fit: see help('isSingular')
```

```
# singular fit
r.squaredGLMM(lmer_COI_UMIspike5) # 0.054 0.85
```

```
## Warning in pcrRep:soupRep: numerical expression has 3150 elements: only the
## first used

## Warning in pcrRep:soupRep: numerical expression has 3150 elements: only the
## first used
```

```
##             R2m       R2c
## [1,] 0.05399271 0.8545995
```

```
# R2m = marginal R-squared (variance explained by fixed effects only)
# R2c = conditional R-squared (variance explained by random + fixed effects)

sresid <- resid(lmer_COI_UMIspike5, type = "pearson") # Extract the standardised residuals, check underline assumption of normally distibuted
hist(sresid)
```

```
plot(lmer_COI_UMIspike5) # gives a heteroscedasticity plot
```

```
# plot(sresid ~ mydata$log.mtagNum)

pred.mm7 <- ggpredict(lmer_COI_UMIspike5, terms = c("log.mtagNum"))

# Plot the predictions
(p7 <- ggplot() +
  geom_line(data = pred.mm7, aes(x = x, y = predicted),
            colour = "black", size = 1) + # slope
  geom_ribbon(
    data = pred.mm7, aes(
      x = x, ymin = predicted - std.error,
      ymax = predicted + std.error
    ),
    fill = "lightgrey", alpha = 0.5
  ) +
  geom_point(data = mydata, aes(
    x = log.mtagNum, y = log.inputCOINum,
    colour = OTUID
  ), size = 0.8) +
  geom_smooth(
    data = mydata, aes(
      x = log.readNum, y = log.inputCOINum,
      group = OTUID
    ),
    method = "lm", se = FALSE, size = 0.3, colour = "black", alpha = 0.5,
    linetype = 1
  ) +
  labs(y = "ln(Input COI copy number)", x = "ln(non-spike-corrected UMI number)") +
  theme_cowplot() +
  theme(legend.position = "none"))
```

```
## `geom_smooth()` using formula 'y ~ x'
```

```
## Warning: Removed 126 rows containing non-finite values (stat_smooth).
```

```
## Warning: Removed 126 rows containing missing values (geom_point).
```

```
lmer_COI_UMIspikeCorr1 <- lmer(log.inputCOINum ~ log.mtagNumSpikeCorr +
  (log.mtagNumSpikeCorr | OTUID) +
  (1 | soupRep / pcrRep), data = mydata, REML = F)
lmer_COI_UMIspikeCorr2 <- lmer(log.inputCOINum ~ log.mtagNumSpikeCorr +
  (1 | OTUID) +
  (1 | soupRep / pcrRep), data = mydata, REML = F)
lmer_COI_UMIspikeCorr3 <- lmer(log.inputCOINum ~ log.mtagNumSpikeCorr +
  (1 | soupRep / pcrRep), data = mydata, REML = F)
```

```
## boundary (singular) fit: see help('isSingular')
```

```
# singular fit

AIC(lmer_COI_UMIspikeCorr1, lmer_COI_UMIspikeCorr2, lmer_COI_UMIspikeCorr3) # lmer_gDNA_UMIspikeCorr1
```

```
##                        df      AIC
## lmer_COI_UMIspikeCorr1  8 1835.076
## lmer_COI_UMIspikeCorr2  6 2344.344
## lmer_COI_UMIspikeCorr3  5 8884.312
```

```
# fixed effect
lmer_COI_UMIspikeCorr4 <- lmer(log.inputCOINum ~ 1 +
  (log.mtagNumSpikeCorr | OTUID) +
  (1 | soupRep / pcrRep), data = mydata, REML = F)

AIC(lmer_COI_UMIspikeCorr1, lmer_COI_UMIspikeCorr4) # lmer_COI_UMIspikeCorr1
```

```
##                        df      AIC
## lmer_COI_UMIspikeCorr1  8 1835.076
## lmer_COI_UMIspikeCorr4  7 1957.893
```

```
anova(lmer_COI_UMIspikeCorr1, lmer_COI_UMIspikeCorr4)
```

```
## Data: mydata
## Models:
## lmer_COI_UMIspikeCorr4: log.inputCOINum ~ 1 + (log.mtagNumSpikeCorr | OTUID) + (1 | soupRep/pcrRep)
## lmer_COI_UMIspikeCorr1: log.inputCOINum ~ log.mtagNumSpikeCorr + (log.mtagNumSpikeCorr | OTUID) + (1 | soupRep/pcrRep)
##                        npar    AIC    BIC  logLik deviance  Chisq Df
## lmer_COI_UMIspikeCorr4    7 1957.9 2000.3 -971.95   1943.9          
## lmer_COI_UMIspikeCorr1    8 1835.1 1883.5 -909.54   1819.1 124.82  1
##                                   Pr(>Chisq)    
## lmer_COI_UMIspikeCorr4                          
## lmer_COI_UMIspikeCorr1 < 0.00000000000000022 ***
## ---
## Signif. codes:  0 '***' 0.001 '**' 0.01 '*' 0.05 '.' 0.1 ' ' 1
```

```
# estimate parameters
lmer_COI_UMIspikeCorr5 <- lmer(log.inputCOINum ~ log.mtagNumSpikeCorr +
  (log.mtagNumSpikeCorr | OTUID) +
  (1 | soupRep / pcrRep), data = mydata, REML = T)
r.squaredGLMM(lmer_COI_UMIspikeCorr5) # 0.434 0.95
```

```
## Warning in pcrRep:soupRep: numerical expression has 3150 elements: only the
## first used

## Warning in pcrRep:soupRep: numerical expression has 3150 elements: only the
## first used
```

```
##            R2m       R2c
## [1,] 0.4343609 0.9501764
```

```
sresid <- resid(lmer_COI_UMIspikeCorr5, type = "pearson")
hist(sresid)
```

```
plot(lmer_COI_UMIspikeCorr5)
```

```
pred.mm8 <- ggpredict(lmer_COI_UMIspikeCorr5, terms = c("log.mtagNumSpikeCorr"))

(p8 <- ggplot() +
  geom_line(data = pred.mm8, aes(x = x, y = predicted),
            colour = "black", size = 1) + # slope
  geom_ribbon(
    data = pred.mm8, aes(
      x = x, ymin = predicted - std.error,
      ymax = predicted + std.error
    ),
    fill = "lightgrey", alpha = 0.5
  ) +
  geom_point(data = mydata, aes(
    x = log.mtagNumSpikeCorr, y = log.inputCOINum,
    colour = OTUID
  ), size = 0.8) +
  geom_smooth(
    data = mydata, aes(
      x = log.mtagNumSpikeCorr, y = log.inputCOINum,
      group = OTUID
    ),
    method = "lm", se = FALSE, size = 0.3, colour = "black", alpha = 0.5,
    linetype = 1
  ) +
  labs(y = "ln(Input COI copy number)", x = "ln(spike-corrected UMI number)") +
  theme_cowplot() +
  theme(legend.position = "none"))
```

```
## `geom_smooth()` using formula 'y ~ x'
```

```
## Warning: Removed 126 rows containing non-finite values (stat_smooth).
```

```
## Warning: Removed 126 rows containing missing values (geom_point).
```

Figure S3 UMI on mocksoups

```
(p7 + p8) / (p5 + p6) + plot_annotation(tag_levels = 'A')
```

```
## `geom_smooth()` using formula 'y ~ x'
```

```
## Warning: Removed 126 rows containing non-finite values (stat_smooth).
```

```
## Warning: Removed 126 rows containing missing values (geom_point).
```

```
## `geom_smooth()` using formula 'y ~ x'
```

```
## Warning: Removed 126 rows containing non-finite values (stat_smooth).
## Removed 126 rows containing missing values (geom_point).
```

```
## `geom_smooth()` using formula 'y ~ x'
## `geom_smooth()` using formula 'y ~ x'
```

```
## Warning: Removed 4 row(s) containing missing values (geom_path).
```

###### Test of Warton spike-in estimator on full dataset, with OTUID and soupRep/pcrRep

Model-based pipeline-noise estimation: offset0 (ã from Eq. 2)

The following code uses the model-based estimator to estimate a
spike-in corrector directly from the dataset, which we find is positive
albeit weakly correlated with the spike-in (R2 = 0.23 and 0.18). N.B.
The mvabund model does not take into account the nested experimental
design (soupRep/PCRrep) reformat data

```
mydata2 <- mydata %>% 
  select(-mtagNum, -exp, -lib, -sampleName, -PCR, -readSpike, -mtagSpike,
         -readNumSpikeCorr, -mtagNumSpikeCorr, -inputCOINum, -inputGDNA,
         -starts_with("log.")) %>% 
  pivot_wider(names_from = OTUID, values_from = readNum)

otu <- mydata2 %>% 
  select(starts_with("HAP"))

X <- mydata2 %>% 
  select(!starts_with("HAP"))
```

```
otu <- mvabund(otu)
# fit0 <- manyglm(otu ~ 1, family = "negative.binomial") # if no predictors
fit0 <- manyglm(otu ~ soup, family = "negative.binomial", data = X) # data = the env covariates. otu is the otu table
# fit0 <- manyglm(otu ~ soup, family = binomial("cloglog")) # for p/a data
offset0 <- log(rowSums(otu)) - log(rowSums(fitted(fit0))) 
mydata2$offset0 <- offset0
mydata2 <- mydata2 %>% 
  relocate(offset0, .after = pcrRep) %>% 
  pivot_longer(cols = starts_with("HAP"), 
               names_to = "OTUID", 
               values_to = "readNum") %>% 
  mutate(
    readNumOffsetCorr = readNum / exp(offset0) # correction
  )

mydata2 <- mydata %>% 
  select(OTUID, soup, soupRep, pcrRep, inputCOINum, inputGDNA, 
         readSpike, mtagSpike) %>% # physical spike-in read numbers
  left_join(mydata2) %>% 
  mutate(
    log.inputCOINum = log(inputCOINum + 1),
    log.inputGDNA = log(inputGDNA + 1),
    log.readNum = log(readNum + 1),
    log.readNumOffsetCorr = log(readNumOffsetCorr + 1),
    log.readSpike = log(readSpike + 1),
    log.mtagSpike = log(mtagSpike + 1)
  )
```

```
## Joining, by = c("OTUID", "soup", "soupRep", "pcrRep")
```

```
names(mydata2)
```

```
##  [1] "OTUID"                 "soup"                  "soupRep"              
##  [4] "pcrRep"                "inputCOINum"           "inputGDNA"            
##  [7] "readSpike"             "mtagSpike"             "offset0"              
## [10] "readNum"               "readNumOffsetCorr"     "log.inputCOINum"      
## [13] "log.inputGDNA"         "log.readNum"           "log.readNumOffsetCorr"
## [16] "log.readSpike"         "log.mtagSpike"
```

```
#  [1] "OTUID"                 "soup"                  "soupRep"              
#  [4] "pcrRep"                "inputCOINum"           "inputGDNA"            
#  [7] "readSpike"             "mtagSpike"             "offset0"              
# [10] "readNum"               "readNumOffsetCorr"     "log.inputCOINum"      
# [13] "log.inputGDNA"         "log.readNum"           "log.readNumOffsetCorr"
# [16] "log.readSpike"         "log.mtagSpike"  

# offset0 is the estimated spike-in
# log.readSpike is the log of the spike-in reads
# log.mtagSpike is the log of the UMIs, which is another way of doing spike-ins

# offset0 is correlated with 
ggplot(data = mydata2, aes(x = (offset0), y = (log.readSpike))) +
  geom_point(aes(colour = soup)) +
  geom_smooth(method = "lm", se = FALSE, size = 0.3, colour = "black", 
              alpha = 0.5, linetype = 1) +
  labs(x = "offset0", y = "ln(readSpike)") +
  theme_cowplot() +
  theme(legend.position = "right")
```

```
## `geom_smooth()` using formula 'y ~ x'
```

```
summary(lm(log.readSpike ~ offset0, data = mydata2))
```

```
## 
## Call:
## lm(formula = log.readSpike ~ offset0, data = mydata2)
## 
## Residuals:
##     Min      1Q  Median      3Q     Max 
## -1.7433 -0.8131  0.1267  0.7000  1.2753 
## 
## Coefficients:
##             Estimate Std. Error t value            Pr(>|t|)    
## (Intercept) 10.35032    0.01445  716.27 <0.0000000000000002 ***
## offset0      0.94670    0.03021   31.34 <0.0000000000000002 ***
## ---
## Signif. codes:  0 '***' 0.001 '**' 0.01 '*' 0.05 '.' 0.1 ' ' 1
## 
## Residual standard error: 0.8052 on 3274 degrees of freedom
## Multiple R-squared:  0.2307, Adjusted R-squared:  0.2305 
## F-statistic:   982 on 1 and 3274 DF,  p-value: < 0.00000000000000022
```

```
# R2 = 0.2307

ggplot(data = mydata2, aes(x = offset0, y = log.mtagSpike, group = OTUID)) +
  geom_point(aes(colour = soup)) +
  geom_smooth(method = "lm", se = FALSE, size = 0.3, colour = "black", 
              alpha = 0.5, linetype = 1) +
  labs(x = "offset0", y = "ln(mtagSpike)") +
  theme_cowplot() +
  theme(legend.position = "none")
```

```
## `geom_smooth()` using formula 'y ~ x'
```

```
summary(lm(log.mtagSpike ~ offset0, data = mydata2))
```

```
## 
## Call:
## lm(formula = log.mtagSpike ~ offset0, data = mydata2)
## 
## Residuals:
##      Min       1Q   Median       3Q      Max 
## -1.24665 -0.41439  0.04642  0.44989  1.08210 
## 
## Coefficients:
##             Estimate Std. Error t value            Pr(>|t|)    
## (Intercept)  9.55150    0.01023  933.25 <0.0000000000000002 ***
## offset0      0.58248    0.02140   27.22 <0.0000000000000002 ***
## ---
## Signif. codes:  0 '***' 0.001 '**' 0.01 '*' 0.05 '.' 0.1 ' ' 1
## 
## Residual standard error: 0.5703 on 3274 degrees of freedom
## Multiple R-squared:  0.1846, Adjusted R-squared:  0.1843 
## F-statistic:   741 on 1 and 3274 DF,  p-value: < 0.00000000000000022
```

```
# R2 = 0.1846
```

We now use these (inferior) model-estimated spike-ins to correct the
OTU sizes and then try to predict the input COI copy number and the
input genomic DNA, just as in Figure 4. The results are poorer than the
spike-in (no surprise), but better than no correction, as can be seen
via inspection of the full model regression lines (the thick black
lines)

prediction of input COI copy number

```
# test random effects
lmer_coi_readOffsetCorr1 <- lmer(log.inputCOINum ~ log.readNumOffsetCorr +
  (log.readNumOffsetCorr | OTUID) +
  (1 | soupRep / pcrRep), data = mydata2, REML = FALSE)
```

```
## boundary (singular) fit: see help('isSingular')
```

```
summary(lmer_coi_readOffsetCorr1)
```

```
## Linear mixed model fit by maximum likelihood  ['lmerMod']
## Formula: log.inputCOINum ~ log.readNumOffsetCorr + (log.readNumOffsetCorr |  
##     OTUID) + (1 | soupRep/pcrRep)
##    Data: mydata2
## 
##      AIC      BIC   logLik deviance df.resid 
##   3946.1   3994.5  -1965.0   3930.1     3142 
## 
## Scaled residuals: 
##      Min       1Q   Median       3Q      Max 
## -1.89709 -0.80779 -0.09339  0.90304  2.29981 
## 
## Random effects:
##  Groups         Name                  Variance Std.Dev. Corr 
##  OTUID          (Intercept)           2.25245  1.5008        
##                 log.readNumOffsetCorr 0.03391  0.1842   -0.77
##  pcrRep:soupRep (Intercept)           0.00000  0.0000        
##  soupRep        (Intercept)           0.00000  0.0000        
##  Residual                             0.18117  0.4256        
## Number of obs: 3150, groups:  OTUID, 50; pcrRep:soupRep, 9; soupRep, 3
## 
## Fixed effects:
##                       Estimate Std. Error t value
## (Intercept)           15.04422    0.23518  63.969
## log.readNumOffsetCorr  0.19337    0.02958   6.538
## 
## Correlation of Fixed Effects:
##             (Intr)
## lg.rdNmOffC -0.807
## optimizer (nloptwrap) convergence code: 0 (OK)
## boundary (singular) fit: see help('isSingular')
```

```
plot(lmer_coi_readOffsetCorr1)
```

```
# singular fit caused by zero variance in pcrRep:soupRep and soupRep, but these random factors must be retained because they are part of the experimental design
lmer_coi_readOffsetCorr2 <- lmer(log.inputCOINum ~ log.readNumOffsetCorr +
  (1 | OTUID) +
  (1 | soupRep / pcrRep), data = mydata2, REML = FALSE)
```

```
## boundary (singular) fit: see help('isSingular')
```

```
summary(lmer_coi_readOffsetCorr2)
```

```
## Linear mixed model fit by maximum likelihood  ['lmerMod']
## Formula: log.inputCOINum ~ log.readNumOffsetCorr + (1 | OTUID) + (1 |  
##     soupRep/pcrRep)
##    Data: mydata2
## 
##      AIC      BIC   logLik deviance df.resid 
##   4077.6   4114.0  -2032.8   4065.6     3144 
## 
## Scaled residuals: 
##      Min       1Q   Median       3Q      Max 
## -2.16666 -0.82056 -0.09142  0.95793  2.42283 
## 
## Random effects:
##  Groups         Name        Variance Std.Dev.
##  OTUID          (Intercept) 0.9943   0.9972  
##  pcrRep:soupRep (Intercept) 0.0000   0.0000  
##  soupRep        (Intercept) 0.0000   0.0000  
##  Residual                   0.1942   0.4407  
## Number of obs: 3150, groups:  OTUID, 50; pcrRep:soupRep, 9; soupRep, 3
## 
## Fixed effects:
##                       Estimate Std. Error t value
## (Intercept)           15.59068    0.15705   99.27
## log.readNumOffsetCorr  0.12065    0.01027   11.74
## 
## Correlation of Fixed Effects:
##             (Intr)
## lg.rdNmOffC -0.437
## optimizer (nloptwrap) convergence code: 0 (OK)
## boundary (singular) fit: see help('isSingular')
```

```
plot(lmer_coi_readOffsetCorr2)
```

```
# singular fit
lmer_coi_readOffsetCorr3 <- lmer(log.inputCOINum ~ log.readNumOffsetCorr +
   (1 | soupRep/pcrRep), data = mydata2, REML = FALSE)
```

```
## boundary (singular) fit: see help('isSingular')
```

```
# singular fit
summary(lmer_coi_readOffsetCorr3)
```

```
## Linear mixed model fit by maximum likelihood  ['lmerMod']
## Formula: log.inputCOINum ~ log.readNumOffsetCorr + (1 | soupRep/pcrRep)
##    Data: mydata2
## 
##      AIC      BIC   logLik deviance df.resid 
##   9187.3   9217.6  -4588.6   9177.3     3145 
## 
## Scaled residuals: 
##     Min      1Q  Median      3Q     Max 
## -2.7254 -0.6845  0.0851  0.6989  3.1712 
## 
## Random effects:
##  Groups         Name        Variance Std.Dev.
##  pcrRep:soupRep (Intercept) 0.000    0.000   
##  soupRep        (Intercept) 0.000    0.000   
##  Residual                   1.078    1.038   
## Number of obs: 3150, groups:  pcrRep:soupRep, 9; soupRep, 3
## 
## Fixed effects:
##                        Estimate Std. Error t value
## (Intercept)           14.563811   0.060189  241.97
## log.readNumOffsetCorr  0.274245   0.008567   32.01
## 
## Correlation of Fixed Effects:
##             (Intr)
## lg.rdNmOffC -0.952
## optimizer (nloptwrap) convergence code: 0 (OK)
## boundary (singular) fit: see help('isSingular')
```

```
plot(lmer_coi_readOffsetCorr3)
```

```
AIC(lmer_coi_readOffsetCorr1, lmer_coi_readOffsetCorr2, lmer_coi_readOffsetCorr3) # lmer_coi_readOffsetCorr1 has lowest AIC, keep random intercept and slope, keep OTUID, soupRep/pcrRep
```

```
##                          df      AIC
## lmer_coi_readOffsetCorr1  8 3946.061
## lmer_coi_readOffsetCorr2  6 4077.622
## lmer_coi_readOffsetCorr3  5 9187.293
```

```
# test fixed effects
lmer_coi_readOffsetCorr4 <- lmer(log.inputCOINum ~ 1 +
  (log.readNumOffsetCorr | OTUID) +
  (1 | soupRep / pcrRep), data = mydata2, REML = F)
```

```
## boundary (singular) fit: see help('isSingular')
```

```
# singular fit
AIC(lmer_coi_readOffsetCorr1, lmer_coi_readOffsetCorr4) # lmer_coi_readOffsetCorr1 lowest AIC
```

```
##                          df      AIC
## lmer_coi_readOffsetCorr1  8 3946.061
## lmer_coi_readOffsetCorr4  7 3975.921
```

```
anova(lmer_coi_readOffsetCorr1, lmer_coi_readOffsetCorr4) # significantly different. keep fixed effect
```

```
## Data: mydata2
## Models:
## lmer_coi_readOffsetCorr4: log.inputCOINum ~ 1 + (log.readNumOffsetCorr | OTUID) + (1 | soupRep/pcrRep)
## lmer_coi_readOffsetCorr1: log.inputCOINum ~ log.readNumOffsetCorr + (log.readNumOffsetCorr | OTUID) + (1 | soupRep/pcrRep)
##                          npar    AIC    BIC logLik deviance Chisq Df
## lmer_coi_readOffsetCorr4    7 3975.9 4018.3  -1981   3961.9         
## lmer_coi_readOffsetCorr1    8 3946.1 3994.5  -1965   3930.1 31.86  1
##                             Pr(>Chisq)    
## lmer_coi_readOffsetCorr4                  
## lmer_coi_readOffsetCorr1 0.00000001657 ***
## ---
## Signif. codes:  0 '***' 0.001 '**' 0.01 '*' 0.05 '.' 0.1 ' ' 1
```

```
# parameter estimates
lmer_coi_readOffsetCorr5 <- lmer(log.inputCOINum ~ log.readNumOffsetCorr +
  (log.readNumOffsetCorr | OTUID) +
  (1 | soupRep / pcrRep), data = mydata2, REML = TRUE)
```

```
## boundary (singular) fit: see help('isSingular')
```

```
# singular fit
summary(lmer_coi_readOffsetCorr5)
```

```
## Linear mixed model fit by REML ['lmerMod']
## Formula: log.inputCOINum ~ log.readNumOffsetCorr + (log.readNumOffsetCorr |  
##     OTUID) + (1 | soupRep/pcrRep)
##    Data: mydata2
## 
## REML criterion at convergence: 3937.4
## 
## Scaled residuals: 
##      Min       1Q   Median       3Q      Max 
## -1.89680 -0.80757 -0.09354  0.90164  2.30274 
## 
## Random effects:
##  Groups         Name                  Variance Std.Dev. Corr 
##  OTUID          (Intercept)           2.31253  1.5207        
##                 log.readNumOffsetCorr 0.03491  0.1868   -0.77
##  pcrRep:soupRep (Intercept)           0.00000  0.0000        
##  soupRep        (Intercept)           0.00000  0.0000        
##  Residual                             0.18116  0.4256        
## Number of obs: 3150, groups:  OTUID, 50; pcrRep:soupRep, 9; soupRep, 3
## 
## Fixed effects:
##                       Estimate Std. Error t value
## (Intercept)           15.04278    0.23778   63.26
## log.readNumOffsetCorr  0.19356    0.02992    6.47
## 
## Correlation of Fixed Effects:
##             (Intr)
## lg.rdNmOffC -0.807
## optimizer (nloptwrap) convergence code: 0 (OK)
## boundary (singular) fit: see help('isSingular')
```

```
r.squaredGLMM(lmer_coi_readOffsetCorr5) # 0.1182 0.8676
```

```
## Warning in pcrRep:soupRep: numerical expression has 3150 elements: only the
## first used

## Warning in pcrRep:soupRep: numerical expression has 3150 elements: only the
## first used
```

```
##            R2m       R2c
## [1,] 0.1181628 0.8775666
```

```
sresid <- resid(lmer_coi_readOffsetCorr5, type = "pearson")
hist(sresid)
```

```
plot(lmer_coi_readOffsetCorr5)
```

```
pred.mm1 <- ggpredict(lmer_coi_readOffsetCorr5, 
                      terms = c("log.readNumOffsetCorr[all]"))

(p9 <- ggplot() +
  geom_line(
    data = pred.mm1, aes(x = x, y = predicted),
    colour = "black", size = 1
  ) +
  geom_ribbon(
    data = pred.mm1, aes(
      x = x, ymin = predicted - std.error,
      ymax = predicted + std.error
    ),
    fill = "lightgrey", alpha = 0.3
  ) +
  geom_point(data = mydata2, aes(
    x = log.readNumOffsetCorr, y = log.inputCOINum,
    colour = OTUID
  ), size = 0.8) +
  geom_smooth(
    data = mydata2, aes(
      x = log.readNumOffsetCorr, y = log.inputCOINum,
      group = OTUID
    ),
    method = "lm", se = FALSE, size = 0.3, colour = "black", alpha = 0.5,
    linetype = 1
  ) +
  labs(x = "ln(Offset-corrected OTU size)", y = "ln(input COI copy number)") +
  theme_cowplot() +
  theme(legend.position = "none"))
```

```
## `geom_smooth()` using formula 'y ~ x'
```

```
## Warning: Removed 126 rows containing non-finite values (stat_smooth).
```

```
## Warning: Removed 126 rows containing missing values (geom_point).
```

should be exactly the same as “COI non-spike-corrected readNum”

```
lmer_coi_read1 <- lmer(log.inputCOINum ~ log.readNum +
  (log.readNum | OTUID) +
  (1 | soupRep / pcrRep), data = mydata2, REML = F)
```

```
## boundary (singular) fit: see help('isSingular')
```

```
# singular fit
lmer_coi_read2 <- lmer(log.inputCOINum ~ log.readNum +
  (1 | OTUID) +
  (1 | soupRep / pcrRep), data = mydata2, REML = F)
```

```
## boundary (singular) fit: see help('isSingular')
```

```
# singular fit
lmer_coi_read3 <- lmer(log.inputCOINum ~ log.readNum +
  (1 | soupRep / pcrRep), data = mydata2, REML = F)
```

```
## boundary (singular) fit: see help('isSingular')
```

```
# singular fit
AIC(lmer_coi_read1, lmer_coi_read2, lmer_coi_read3) # choose lmer_coi_read1
```

```
##                df      AIC
## lmer_coi_read1  8 4067.218
## lmer_coi_read2  6 4113.070
## lmer_coi_read3  5 9223.563
```

```
# test fixed effect
lmer_coi_read4 <- lmer(log.inputCOINum ~ 1 +
  (log.readNum | OTUID) +
  (1 | soupRep / pcrRep), data = mydata2, REML = F)
```

```
## boundary (singular) fit: see help('isSingular')
```

```
# singular fit
AIC(lmer_coi_read1, lmer_coi_read4)
```

```
##                df      AIC
## lmer_coi_read1  8 4067.218
## lmer_coi_read4  7 4095.451
```

```
anova(lmer_coi_read1, lmer_coi_read4) # keep fixed effect
```

```
## Data: mydata2
## Models:
## lmer_coi_read4: log.inputCOINum ~ 1 + (log.readNum | OTUID) + (1 | soupRep/pcrRep)
## lmer_coi_read1: log.inputCOINum ~ log.readNum + (log.readNum | OTUID) + (1 | soupRep/pcrRep)
##                npar    AIC    BIC  logLik deviance  Chisq Df    Pr(>Chisq)    
## lmer_coi_read4    7 4095.5 4137.8 -2040.7   4081.5                            
## lmer_coi_read1    8 4067.2 4115.7 -2025.6   4051.2 30.233  1 0.00000003832 ***
## ---
## Signif. codes:  0 '***' 0.001 '**' 0.01 '*' 0.05 '.' 0.1 ' ' 1
```

```
# estimate parameters with lmer_coi_read1
lmer_coi_read5 <- lmer(log.inputCOINum ~ log.readNum +
  (log.readNum | OTUID) +
  (1 | soupRep / pcrRep), data = mydata2, REML = TRUE)
```

```
## boundary (singular) fit: see help('isSingular')
```

```
# singular fit
summary(lmer_coi_read5)
```

```
## Linear mixed model fit by REML ['lmerMod']
## Formula: log.inputCOINum ~ log.readNum + (log.readNum | OTUID) + (1 |  
##     soupRep/pcrRep)
##    Data: mydata2
## 
## REML criterion at convergence: 4059.6
## 
## Scaled residuals: 
##      Min       1Q   Median       3Q      Max 
## -1.94255 -0.80640 -0.02837  0.92313  1.99929 
## 
## Random effects:
##  Groups         Name        Variance Std.Dev. Corr 
##  OTUID          (Intercept) 1.332730 1.15444       
##                 log.readNum 0.008726 0.09341  -0.50
##  pcrRep:soupRep (Intercept) 0.000000 0.00000       
##  soupRep        (Intercept) 0.000000 0.00000       
##  Residual                   0.190370 0.43631       
## Number of obs: 3150, groups:  OTUID, 50; pcrRep:soupRep, 9; soupRep, 3
## 
## Fixed effects:
##             Estimate Std. Error t value
## (Intercept) 15.67884    0.18015  87.034
## log.readNum  0.10655    0.01714   6.216
## 
## Correlation of Fixed Effects:
##             (Intr)
## log.readNum -0.609
## optimizer (nloptwrap) convergence code: 0 (OK)
## boundary (singular) fit: see help('isSingular')
```

```
r.squaredGLMM(lmer_coi_read5) # 0.042 0.85
```

```
## Warning in pcrRep:soupRep: numerical expression has 3150 elements: only the
## first used

## Warning in pcrRep:soupRep: numerical expression has 3150 elements: only the
## first used
```

```
##             R2m       R2c
## [1,] 0.04196802 0.8528916
```

```
# R2m = marginal R-squared (variance explained by fixed effects only)
# R2c = conditional R-squared (variance explained by random + fixed effects)

sresid <- resid(lmer_coi_read5, type = "pearson") # Extract the standardised residuals, check underline assumption of normally distibuted
hist(sresid)
```

```
plot(lmer_coi_read5) # gives a heteroscedasticity plot
```

```
pred.mm2 <- ggpredict(lmer_coi_read5, terms = c("log.readNum"))

# Plot the predictions
(p10 <- ggplot() +
  geom_line(
    data = pred.mm2, aes(x = x, y = predicted),
    colour = "black", size = 1
  ) + # slope
  geom_ribbon(
    data = pred.mm2, aes(
      x = x, ymin = predicted - std.error,
      ymax = predicted + std.error
    ),
    fill = "lightgrey", alpha = 0.5
  ) +
  geom_point(data = mydata2, aes(
    x = log.readNum, y = log.inputCOINum,
    colour = OTUID
  ), size = 0.8) +
  geom_smooth(
    data = mydata2, aes(
      x = log.readNum, y = log.inputCOINum,
      group = OTUID
    ),
    method = "lm", se = FALSE, size = 0.3, colour = "black", alpha = 0.5,
    linetype = 1
  ) +
  labs(x = "ln(non-Offset-corrected OTU size)", y = "ln(input COI copy number)") +
  theme_cowplot() +
  theme(legend.position = "none"))
```

```
## `geom_smooth()` using formula 'y ~ x'
```

```
## Warning: Removed 126 rows containing non-finite values (stat_smooth).
```

```
## Warning: Removed 126 rows containing missing values (geom_point).
```

```
p10 + p9
```

```
## `geom_smooth()` using formula 'y ~ x'
```

```
## Warning: Removed 126 rows containing non-finite values (stat_smooth).
## Removed 126 rows containing missing values (geom_point).
```

```
## `geom_smooth()` using formula 'y ~ x'
```

```
## Warning: Removed 126 rows containing non-finite values (stat_smooth).
## Removed 126 rows containing missing values (geom_point).
```

prediction of input genomic DNA

```
lmer_gDNA_readOffsetCorr1 <- lmer(log.inputGDNA ~ log.readNumOffsetCorr +
   (log.readNumOffsetCorr | OTUID) +
   (1 | soupRep / pcrRep), data = mydata2, REML = FALSE)
```

```
## boundary (singular) fit: see help('isSingular')
```

```
# singular fit
summary(lmer_gDNA_readOffsetCorr1)
```

```
## Linear mixed model fit by maximum likelihood  ['lmerMod']
## Formula: log.inputGDNA ~ log.readNumOffsetCorr + (log.readNumOffsetCorr |  
##     OTUID) + (1 | soupRep/pcrRep)
##    Data: mydata2
## 
##      AIC      BIC   logLik deviance df.resid 
##   3761.4   3810.1  -1872.7   3745.4     3268 
## 
## Scaled residuals: 
##      Min       1Q   Median       3Q      Max 
## -1.75640 -0.81964 -0.08629  0.94979  2.21610 
## 
## Random effects:
##  Groups         Name                  Variance Std.Dev. Corr 
##  OTUID          (Intercept)           0.83390  0.9132        
##                 log.readNumOffsetCorr 0.01659  0.1288   -0.97
##  pcrRep:soupRep (Intercept)           0.00000  0.0000        
##  soupRep        (Intercept)           0.00000  0.0000        
##  Residual                             0.17106  0.4136        
## Number of obs: 3276, groups:  OTUID, 52; pcrRep:soupRep, 9; soupRep, 3
## 
## Fixed effects:
##                       Estimate Std. Error t value
## (Intercept)             2.5487     0.1507  16.915
## log.readNumOffsetCorr   0.1343     0.0212   6.336
## 
## Correlation of Fixed Effects:
##             (Intr)
## lg.rdNmOffC -0.969
## optimizer (nloptwrap) convergence code: 0 (OK)
## boundary (singular) fit: see help('isSingular')
```

```
lmer_gDNA_readOffsetCorr2 <- lmer(log.inputGDNA ~ log.readNumOffsetCorr +
   (1 | OTUID) +
   (1 | soupRep / pcrRep), data = mydata2, REML = FALSE)
```

```
## boundary (singular) fit: see help('isSingular')
```

```
# singular fit
lmer_gDNA_readOffsetCorr3 <- lmer(log.inputGDNA ~ log.readNumOffsetCorr +
   (1 | soupRep / pcrRep), data = mydata2, REML = FALSE)
```

```
## boundary (singular) fit: see help('isSingular')
```

```
# singular fit

AIC(lmer_gDNA_readOffsetCorr1, lmer_gDNA_readOffsetCorr2, lmer_gDNA_readOffsetCorr3) # lmer_gDNA_readOffsetCorr1 has lower AIC, use random intercept and slope, keep OTUID, soupRep, pcrRep
```

```
##                           df      AIC
## lmer_gDNA_readOffsetCorr1  8 3761.374
## lmer_gDNA_readOffsetCorr2  6 3782.993
## lmer_gDNA_readOffsetCorr3  5 3780.993
```

```
# test fixed effects
lmer_gDNA_readOffsetCorr4 <- lmer(log.inputGDNA ~ 1 +
   (log.readNumOffsetCorr | OTUID) +
   (1 | soupRep / pcrRep), data = mydata2, REML = F)
```

```
## boundary (singular) fit: see help('isSingular')
```

```
# singular fit
AIC(lmer_gDNA_readOffsetCorr1, lmer_gDNA_readOffsetCorr4) # choose lmer_gDNA_readOffsetCorr1
```

```
##                           df      AIC
## lmer_gDNA_readOffsetCorr1  8 3761.374
## lmer_gDNA_readOffsetCorr4  7 3788.290
```

```
anova(lmer_gDNA_readOffsetCorr1, lmer_gDNA_readOffsetCorr4) # significantly different. keep fixed effect
```

```
## Data: mydata2
## Models:
## lmer_gDNA_readOffsetCorr4: log.inputGDNA ~ 1 + (log.readNumOffsetCorr | OTUID) + (1 | soupRep/pcrRep)
## lmer_gDNA_readOffsetCorr1: log.inputGDNA ~ log.readNumOffsetCorr + (log.readNumOffsetCorr | OTUID) + (1 | soupRep/pcrRep)
##                           npar    AIC    BIC  logLik deviance  Chisq Df
## lmer_gDNA_readOffsetCorr4    7 3788.3 3831.0 -1887.1   3774.3          
## lmer_gDNA_readOffsetCorr1    8 3761.4 3810.1 -1872.7   3745.4 28.916  1
##                              Pr(>Chisq)    
## lmer_gDNA_readOffsetCorr4                  
## lmer_gDNA_readOffsetCorr1 0.00000007561 ***
## ---
## Signif. codes:  0 '***' 0.001 '**' 0.01 '*' 0.05 '.' 0.1 ' ' 1
```

```
# fit model
lmer_gDNA_readOffsetCorr5 <- lmer(log.inputGDNA ~ log.readNumOffsetCorr +
   (log.readNumOffsetCorr | OTUID) +
   (1 | soupRep / pcrRep), data = mydata2, REML = TRUE)
```

```
## boundary (singular) fit: see help('isSingular')
```

```
# singular fit
summary(lmer_gDNA_readOffsetCorr5)
```

```
## Linear mixed model fit by REML ['lmerMod']
## Formula: log.inputGDNA ~ log.readNumOffsetCorr + (log.readNumOffsetCorr |  
##     OTUID) + (1 | soupRep/pcrRep)
##    Data: mydata2
## 
## REML criterion at convergence: 3755.9
## 
## Scaled residuals: 
##      Min       1Q   Median       3Q      Max 
## -1.76293 -0.82007 -0.08666  0.94689  2.22344 
## 
## Random effects:
##  Groups         Name                  Variance Std.Dev. Corr 
##  OTUID          (Intercept)           0.89243  0.9447        
##                 log.readNumOffsetCorr 0.01766  0.1329   -0.96
##  pcrRep:soupRep (Intercept)           0.00000  0.0000        
##  soupRep        (Intercept)           0.00000  0.0000        
##  Residual                             0.17087  0.4134        
## Number of obs: 3276, groups:  OTUID, 52; pcrRep:soupRep, 9; soupRep, 3
## 
## Fixed effects:
##                       Estimate Std. Error t value
## (Intercept)            2.53028    0.15483  16.342
## log.readNumOffsetCorr  0.13692    0.02174   6.299
## 
## Correlation of Fixed Effects:
##             (Intr)
## lg.rdNmOffC -0.968
## optimizer (nloptwrap) convergence code: 0 (OK)
## boundary (singular) fit: see help('isSingular')
```

```
r.squaredGLMM(lmer_gDNA_readOffsetCorr5) # 0.213 0.571
```

```
## Warning in pcrRep:soupRep: numerical expression has 3276 elements: only the
## first used

## Warning in pcrRep:soupRep: numerical expression has 3276 elements: only the
## first used
```

```
##            R2m       R2c
## [1,] 0.2133774 0.5712466
```

```
sresid <- resid(lmer_gDNA_readOffsetCorr5, type = "pearson")
hist(sresid)
```

```
plot(lmer_gDNA_readOffsetCorr5)
```

```
plot(sresid ~ mydata2$log.readNumOffsetCorr)
```

```
#
pred.mm3 <- ggpredict(lmer_gDNA_readOffsetCorr5, 
                      terms = c("log.readNumOffsetCorr[all]"))

(p11 <- ggplot() +
  geom_line(
    data = pred.mm3, aes(x = x, y = predicted),
    colour = "black", size = 1
  ) + 
  geom_ribbon(
    data = pred.mm3, aes(
      x = x, ymin = predicted - std.error,
      ymax = predicted + std.error
    ),
    fill = "lightgrey", alpha = 0.5
  ) +
  geom_point(data = mydata2, aes(
    x = log.readNumOffsetCorr, y = log.inputGDNA,
    colour = OTUID
  ), size = 0.8) +
  geom_smooth(
    data = mydata2, aes(
      x = log.readNumOffsetCorr, y = log.inputGDNA,
      group = OTUID
    ),
    method = "lm", se = FALSE, size = 0.3, colour = "black", alpha = 0.5,
    linetype = 1
  ) +
  ylim(2.3, 4.7) +
  labs(x = "ln(Offset-corrected OTU size)", y = "ln(input genomic DNA)") +
  theme_cowplot() +
  theme(legend.position = "none"))
```

```
## `geom_smooth()` using formula 'y ~ x'
```

should be exactly the same as “gDNA non-spike-corrected readNum”

```
lmer_gDNA_read1 <- lmer(log.inputGDNA ~ log.readNum +
  (log.readNum | OTUID) +
  (1 | soupRep / pcrRep), data = mydata2, REML = F)
```

```
## boundary (singular) fit: see help('isSingular')
```

```
# singular fit
summary(lmer_gDNA_read1)
```

```
## Linear mixed model fit by maximum likelihood  ['lmerMod']
## Formula: 
## log.inputGDNA ~ log.readNum + (log.readNum | OTUID) + (1 | soupRep/pcrRep)
##    Data: mydata2
## 
##      AIC      BIC   logLik deviance df.resid 
##   3806.4   3855.2  -1895.2   3790.4     3268 
## 
## Scaled residuals: 
##      Min       1Q   Median       3Q      Max 
## -1.62394 -0.91102 -0.03561  0.94808  1.69987 
## 
## Random effects:
##  Groups         Name        Variance  Std.Dev. Corr 
##  OTUID          (Intercept) 0.0258659 0.16083       
##                 log.readNum 0.0005691 0.02386  -1.00
##  pcrRep:soupRep (Intercept) 0.0000000 0.00000       
##  soupRep        (Intercept) 2.4244882 1.55708       
##  Residual                   0.1831074 0.42791       
## Number of obs: 3276, groups:  OTUID, 52; pcrRep:soupRep, 9; soupRep, 3
## 
## Fixed effects:
##             Estimate Std. Error t value
## (Intercept) 3.321853   0.899828   3.692
## log.readNum 0.022768   0.005632   4.042
## 
## Correlation of Fixed Effects:
##             (Intr)
## log.readNum -0.042
## optimizer (nloptwrap) convergence code: 0 (OK)
## boundary (singular) fit: see help('isSingular')
```

```
lmer_gDNA_read2 <- lmer(log.inputGDNA ~ log.readNum +
  (1 | OTUID) +
  (1 | soupRep / pcrRep), data = mydata2, REML = F)
```

```
## boundary (singular) fit: see help('isSingular')
```

```
# singular fit
lmer_gDNA_read3 <- lmer(log.inputGDNA ~ log.readNum +
  (1 | soupRep / pcrRep), data = mydata2, REML = F)
```

```
## boundary (singular) fit: see help('isSingular')
```

```
# singular fit

AIC(lmer_gDNA_read1, lmer_gDNA_read2, lmer_gDNA_read3) # lmer_gDNA_read3
```

```
##                 df      AIC
## lmer_gDNA_read1  8 3806.395
## lmer_gDNA_read2  6 3784.565
## lmer_gDNA_read3  5 3782.565
```

```
# fixed effect
lmer_gDNA_read4 <- lmer(log.inputGDNA ~ 1 +
  (1 | soupRep / pcrRep), data = mydata2, REML = F)
```

```
## boundary (singular) fit: see help('isSingular')
```

```
# singular fit
AIC(lmer_gDNA_read3, lmer_gDNA_read4) # keep fixed effect
```

```
##                 df      AIC
## lmer_gDNA_read3  5 3782.565
## lmer_gDNA_read4  4 3796.578
```

```
anova(lmer_gDNA_read3, lmer_gDNA_read4)
```

```
## Data: mydata2
## Models:
## lmer_gDNA_read4: log.inputGDNA ~ 1 + (1 | soupRep/pcrRep)
## lmer_gDNA_read3: log.inputGDNA ~ log.readNum + (1 | soupRep/pcrRep)
##                 npar    AIC  BIC  logLik deviance  Chisq Df Pr(>Chisq)    
## lmer_gDNA_read4    4 3796.6 3821 -1894.3   3788.6                         
## lmer_gDNA_read3    5 3782.6 3813 -1886.3   3772.6 16.013  1  0.0000629 ***
## ---
## Signif. codes:  0 '***' 0.001 '**' 0.01 '*' 0.05 '.' 0.1 ' ' 1
```

```
######## final model###########
lmer_gDNA_read5 <- lmer(log.inputGDNA ~ log.readNum +
  (1 | soupRep / pcrRep), data = mydata2, REML = T)
```

```
## boundary (singular) fit: see help('isSingular')
```

```
# singular fit
r.squaredGLMM(lmer_gDNA_read5) # 0.00487 0.005
```

```
## Warning in pcrRep:soupRep: numerical expression has 3276 elements: only the
## first used

## Warning in pcrRep:soupRep: numerical expression has 3276 elements: only the
## first used
```

```
##              R2m         R2c
## [1,] 0.004874685 0.004874685
```

```
# R2m = marginal R-squared (variance explained by fixed effects only)
# R2c = conditional R-squared (variance explained by random + fixed effects)

sresid <- resid(lmer_gDNA_read5, type = "pearson") # Extract the standardised residuals, check underline assumption of normally distibuted
hist(sresid)
```

```
plot(lmer_gDNA_read5) # gives a heteroscedasticity plot
```

```
plot(sresid ~ mydata2$log.readNum)
```

```
pred.mm4 <- ggpredict(lmer_gDNA_read5, terms = c("log.readNum[all]"))

# Plot the predictions
(p12 <- ggplot() +
  geom_line(
    data = pred.mm4, aes(x = x, y = predicted),
    colour = "black", size = 1
  ) + # slope
  geom_ribbon(
    data = pred.mm4, aes(
      x = x, ymin = predicted - std.error,
      ymax = predicted + std.error
    ),
    fill = "lightgrey", alpha = 0.5
  ) +
  geom_point(data = mydata2, aes(
    x = log.readNum, y = log.inputGDNA,
    colour = OTUID
  ), size = 0.8) +
  geom_smooth(
    data = mydata2, aes(
      x = log.readNum, y = log.inputGDNA,
      group = OTUID
    ),
    method = "lm", se = FALSE, size = 0.3, colour = "black", alpha = 0.5,
    linetype = 1
  ) +
  ylim(2.3, 4.7) +
  labs(x = "ln(non-Offset-corrected OTU size)", y = "ln(Input Genomic DNA)") +
  theme_cowplot() +
  theme(legend.position = "none"))
```

```
## `geom_smooth()` using formula 'y ~ x'
```

```
p11 + p12
```

```
## `geom_smooth()` using formula 'y ~ x'
```

```
## `geom_smooth()` using formula 'y ~ x'
```

There is only a small benefit of using the model-estimated spike-in
correctors on this dataset. The thick black regression lines are the
whole-model fits, and they are steeper in the model-estimated
“offset-corrected” models on the right. The thin lines are fit to each
OTU separately.

```
(p10 + p9) / (p12 + p11)
```

```
## `geom_smooth()` using formula 'y ~ x'
```

```
## Warning: Removed 126 rows containing non-finite values (stat_smooth).
```

```
## Warning: Removed 126 rows containing missing values (geom_point).
```

```
## `geom_smooth()` using formula 'y ~ x'
```

```
## Warning: Removed 126 rows containing non-finite values (stat_smooth).
## Removed 126 rows containing missing values (geom_point).
```

```
## `geom_smooth()` using formula 'y ~ x'
## `geom_smooth()` using formula 'y ~ x'
```
