## Supplementary material for "Extracting abundance information from DNA-based data": S5: Malaise-trap-mix-effects-model_20220731_OPEN_IN_CHROME.html

Malaise trap dilution gradient experiment, general linear mixed-effects model


### Malaise trap dilution gradient experiment, general linear mixed-effects model

###### Doug

###### 31 July 2022

```
dd <- read.csv(here('data','Malaise-trap_Samples_20211228.csv'), 
               header = T)
str(dd)
```

```
## 'data.frame':    4539 obs. of  17 variables:
##  $ OTU                   : chr  "OTU10" "OTU1042" "OTU106" "OTU108" ...
##  $ sampleName            : chr  "123545.M1.S1.A" "123545.M1.S1.A" "123545.M1.S1.A" "123545.M1.S1.A" ...
##  $ OTUsize               : int  102 10 2098 1207 3160 129 2488 1421 668 119 ...
##  $ dilution              : chr  "A" "A" "A" "A" ...
##  $ sample                : chr  "123545.M1.S1" "123545.M1.S1" "123545.M1.S1" "123545.M1.S1" ...
##  $ readSpikeSum          : num  9221 9221 9221 9221 9221 ...
##  $ inputDNA              : num  1 1 1 1 1 1 1 1 1 1 ...
##  $ added.lysis.buffer.ml.: num  13.5 13.5 13.5 13.5 13.5 13.5 13.5 13.5 13.5 13.5 ...
##  $ WEIGHT.g.             : num  2.31 2.31 2.31 2.31 2.31 2.31 2.31 2.31 2.31 2.31 ...
##  $ period                : int  1 1 1 1 1 1 1 1 1 1 ...
##  $ lysis.batch           : int  5 5 5 5 5 5 5 5 5 5 ...
##  $ aliquot               : num  0.6 0.6 0.6 0.6 0.6 0.6 0.6 0.6 0.6 0.6 ...
##  $ lysisBufferFrac       : num  0.0444 0.0444 0.0444 0.0444 0.0444 ...
##  $ FSL                   : num  252.3 24.7 5189.3 2985.4 7816.1 ...
##  $ sample2               : chr  "Sample 1" "Sample 1" "Sample 1" "Sample 1" ...
##  $ fittedOTUsize         : num  -0.352 -0.155 -0.38 -0.369 -0.36 ...
##  $ fittedFSL             : num  -0.4627 -0.0892 -0.1504 -0.3064 -0.0943 ...
```

```
dd$log.inputDNA <- log(dd$inputDNA)
dd$log.OTUsize <- log(dd$OTUsize)
dd$log.FSL <- log(dd$FSL)

plot_missing(dd)
```

```
plot_histogram(dd)
```

[1] “OTU” “sampleName” “OTUsize”  
[4] “dilution” “sample” “readSpikeSum” [7] “inputDNA”
“added.lysis.buffer.ml.” “WEIGHT.g.” [10] “period” “lysis.batch”
“aliquot” [13] “lysisBufferFrac” “FSL” “sample2” [16] “fittedOTUsize”
“fittedFSL” “log.inputDNA” [19] “log.OTUsize” “log.FSL”

relevant variable names

OTU: species

sampleName: Malaise trap *and* dilution

sample: Malaise trap name only

readSpikeSum: number of spike-in reads, summed over the two spike-in
species used in this experiment

OTUsize: number of reads per OTU in that sample and dilution level,
uncorrected for pipeline noise

FSL = OTUsize / (lysisBufferFrac \* readSpikeSum): number of reads per
OTU in that sample and dilution level, corrected for the fact that each
sample (Malaise trap) has a different amount of starting biomass. We
digest the DNA in “lysis buffer”, and we use a fixed aliquot of that
buffer, which therefore accounts for a different fraction of the total
lysis buffer (lysisBufferFrac). The spike-in correct for pipeline noise.
The product lysisBufferFrac\*readSpikeSum thus makes OTU size comparable
between samples. (FSL is from Ji et al. 2019 SPIKEPIPE, introduced by
Ovaskainen in his analysis of that dataset)

dilution: dilution level (A is undiluted, F is most diluted)
inputDNA: dilution level in quantitative terms (proportion of DNA left
in this sample after dilution, 1.00 == undiluted and 0.16 == most
diluted)

```
p1 <- ggplot(dd, 
             aes(x = sample, y = readSpikeSum, 
                 colour = dilution)) +
  geom_point()

p2 <- ggplot(dd, 
             aes(x = sample, y = lysisBufferFrac, 
                 colour = dilution))+
  geom_point()

p1 / p2
```

```
# one value of lysisBufferFrac per sample
# 6 values of readSpikeSum per sample (one per dilution level). The F points (pink) should be highest, which they are for 6 of 7 samples. So there is some error, as expected.
```

Checking that the FSL value was correctly calculated FSL =
1000(OTUsize / (lysisBufferFrac \* readSpikeSum))

```
dd2 <- dd %>% 
  mutate(FSL2 = 1000*(OTUsize / (lysisBufferFrac * readSpikeSum)))
plot(FSL ~ FSL2, data = dd2) # yes it was correctly calculated
```

```
rm(dd2) # delete dd2
```

```
# lmer1 <- lmer(log.inputDNA ~ log.FSL + (1 | sample), data = dd, 
#               REML = F)
# summary(lmer1)
# plot(lmer1)
# 
# lmer2 <-lmer(log.inputDNA ~ log.FSL + (1 | OTU), data = dd, REML = F)
# summary(lmer2)
# plot(lmer2)
# lmer1 and 2 are not useful models, given the experimental design of different sets of OTUs nested within each sample (= trap)

lmer3 <- lmer(log.inputDNA ~ log.FSL + (1 | sample/OTU), 
              data = dd, REML = F)
summary(lmer3)
```

```
## Linear mixed model fit by maximum likelihood  ['lmerMod']
## Formula: log.inputDNA ~ log.FSL + (1 | sample/OTU)
##    Data: dd
## 
##      AIC      BIC   logLik deviance df.resid 
##   4131.3   4163.4  -2060.6   4121.3     4534 
## 
## Scaled residuals: 
##     Min      1Q  Median      3Q     Max 
## -3.8275 -0.6082 -0.0023  0.5904  3.6220 
## 
## Random effects:
##  Groups     Name        Variance Std.Dev.
##  OTU:sample (Intercept) 1.10551  1.0514  
##  sample     (Intercept) 0.17927  0.4234  
##  Residual               0.06011  0.2452  
## Number of obs: 4539, groups:  OTU:sample, 892; sample, 7
## 
## Fixed effects:
##              Estimate Std. Error t value
## (Intercept) -3.903144   0.165448  -23.59
## log.FSL      0.526516   0.003517  149.69
## 
## Correlation of Fixed Effects:
##         (Intr)
## log.FSL -0.122
```

```
plot(lmer3)
```

```
lmer4<- lmer(log.inputDNA ~ log.FSL +(log.FSL | sample/OTU), 
             data = dd, REML = F)
```

```
## boundary (singular) fit: see help('isSingular')
```

```
summary(lmer4)
```

```
## Linear mixed model fit by maximum likelihood  ['lmerMod']
## Formula: log.inputDNA ~ log.FSL + (log.FSL | sample/OTU)
##    Data: dd
## 
##      AIC      BIC   logLik deviance df.resid 
##   4034.9   4092.6  -2008.4   4016.9     4530 
## 
## Scaled residuals: 
##     Min      1Q  Median      3Q     Max 
## -3.9028 -0.6136 -0.0017  0.5986  3.6801 
## 
## Random effects:
##  Groups     Name        Variance   Std.Dev. Corr 
##  OTU:sample (Intercept) 0.97734696 0.988609      
##             log.FSL     0.00008098 0.008999 1.00 
##  sample     (Intercept) 0.23045459 0.480057      
##             log.FSL     0.00128142 0.035797 -0.39
##  Residual               0.05841157 0.241685      
## Number of obs: 4539, groups:  OTU:sample, 892; sample, 7
## 
## Fixed effects:
##             Estimate Std. Error t value
## (Intercept) -3.89126    0.18589  -20.93
## log.FSL      0.52764    0.01401   37.67
## 
## Correlation of Fixed Effects:
##         (Intr)
## log.FSL -0.393
## optimizer (nloptwrap) convergence code: 0 (OK)
## boundary (singular) fit: see help('isSingular')
```

```
# singular fit
plot(lmer4)
```

```
# use lmer3 random factor structure since including log.FSL causes a singular fit error
# test if we can remove fixed effect log.FSL
lmer5 <- lmer(log.inputDNA ~ 1 + (1 | sample/OTU), 
              data = dd, REML = F)
```

```
## boundary (singular) fit: see help('isSingular')
```

```
summary(lmer5)
```

```
## Linear mixed model fit by maximum likelihood  ['lmerMod']
## Formula: log.inputDNA ~ 1 + (1 | sample/OTU)
##    Data: dd
## 
##      AIC      BIC   logLik deviance df.resid 
##   8369.1   8394.7  -4180.5   8361.1     4535 
## 
## Scaled residuals: 
##     Min      1Q  Median      3Q     Max 
## -1.5652 -0.8981  0.2763  0.8631  1.4500 
## 
## Random effects:
##  Groups     Name        Variance                  Std.Dev.       
##  OTU:sample (Intercept) 0.00000000000000000002545 0.0000000001595
##  sample     (Intercept) 0.00000000000000018110707 0.0000000134576
##  Residual               0.36941804442294123855817 0.6077977002449
## Number of obs: 4539, groups:  OTU:sample, 892; sample, 7
## 
## Fixed effects:
##              Estimate Std. Error t value
## (Intercept) -0.881277   0.009022  -97.69
## optimizer (nloptwrap) convergence code: 0 (OK)
## boundary (singular) fit: see help('isSingular')
```

```
# singular fit
plot(lmer5)
```

```
AIC (lmer3, lmer5) # lower AIC when log.FSL fixed effect is retained
```

```
##       df      AIC
## lmer3  5 4131.267
## lmer5  4 8369.068
```

```
anova(lmer4, lmer5) # keep log.FSL fixed effect
```

```
## Data: dd
## Models:
## lmer5: log.inputDNA ~ 1 + (1 | sample/OTU)
## lmer4: log.inputDNA ~ log.FSL + (log.FSL | sample/OTU)
##       npar    AIC    BIC  logLik deviance  Chisq Df            Pr(>Chisq)    
## lmer5    4 8369.1 8394.7 -4180.5   8361.1                                    
## lmer4    9 4034.9 4092.6 -2008.4   4016.9 4344.2  5 < 0.00000000000000022 ***
## ---
## Signif. codes:  0 '***' 0.001 '**' 0.01 '*' 0.05 '.' 0.1 ' ' 1
```

```
# estimate parameters
model.final <- lmer(log.inputDNA ~ log.FSL + (1 | sample/OTU),
                    data = dd, REML = T)
summary(model.final)
```

```
## Linear mixed model fit by REML ['lmerMod']
## Formula: log.inputDNA ~ log.FSL + (1 | sample/OTU)
##    Data: dd
## 
## REML criterion at convergence: 4132.4
## 
## Scaled residuals: 
##     Min      1Q  Median      3Q     Max 
## -3.8272 -0.6080 -0.0024  0.5902  3.6216 
## 
## Random effects:
##  Groups     Name        Variance Std.Dev.
##  OTU:sample (Intercept) 1.10567  1.0515  
##  sample     (Intercept) 0.21026  0.4585  
##  Residual               0.06013  0.2452  
## Number of obs: 4539, groups:  OTU:sample, 892; sample, 7
## 
## Fixed effects:
##              Estimate Std. Error t value
## (Intercept) -3.903354   0.178331  -21.89
## log.FSL      0.526538   0.003518  149.68
## 
## Correlation of Fixed Effects:
##         (Intr)
## log.FSL -0.113
```

```
AIC(model.final) # 4142.43
```

```
## [1] 4142.431
```

```
sresid <- resid(model.final, type = "pearson") # Extract the standardised residuals, check underline assumption of normally distibuted
hist(sresid) # normal residuals
```

```
plot(model.final)
```

```
plot(sresid ~ log.FSL, data = dd) # log.FSL is the spike-corrected OTU size, and the residuals show greater dispersion at lower log.FSL values, meaning that OTU size does a worse job of predicting input gDNA when the OTUs are small, which is not surprising.
```

```
r.squaredGLMM(model.final) #0.53, 0.98
```

```
## Warning: 'r.squaredGLMM' now calculates a revised statistic. See the help page.
```

```
##            R2m       R2c
## [1,] 0.5302842 0.9794754
```

```
pred.mm1 <- ggpredict(model.final, terms = c("log.FSL[all]"))

# for labelling facets
dd$sample2 <-  str_replace_all(dd$sample, c("123545.M1.S1" = "Sample 1", "124031.M1.S1" = "Sample 2", "286789.M1.S1" = "Sample 3", "357256.M2.S1" = "Sample 4", "700239.M1.S1" = "Sample 5", "HB.053.M1.S1" = "Sample 6", "HB.216.M1.S1" = "Sample 7"))

(p1 <- ggplot() +
  geom_line(data = pred.mm1, 
            aes(x = x, y = predicted), 
            colour = "black", size = 1) +
  geom_ribbon(data = pred.mm1, 
              aes(x = x, 
                  ymin = predicted - std.error, 
                  ymax = predicted + std.error), 
              fill = "lightgrey", alpha = 0.5) +
  geom_point(data = dd, aes(x = log.FSL, y = log.inputDNA, 
                             colour = OTU), 
             size = 0.8) +
  geom_smooth(data = dd, aes(x = log.FSL, y = log.inputDNA, 
                              group = OTU),
              method = "lm", se = FALSE, size = 0.1, 
              colour = "black", alpha = 0.5, linetype = 1) +
  labs(x = "log(spike-corrected OTU size)", 
       y = "log(input genomic DNA mass)") +
  theme_cowplot() +
  theme(legend.position = "none") + 
  facet_wrap(~sample2) +
  coord_cartesian(ylim = c(-2.0, 0.25)) +
  scale_y_continuous(breaks = seq(-2.5, 0.5, 0.5)))
```

```
## `geom_smooth()` using formula 'y ~ x'
```

```
# lmer_otusize1 <- lmer(log.inputDNA ~ log.OTUsize + (1| sample), 
#                       data = dd, REML = F )
lmer_otusize2 <- lmer(log.inputDNA ~ log.OTUsize + 
                        (1| sample/OTU),
                      data = dd, REML = F)
```

```
## boundary (singular) fit: see help('isSingular')
```

```
summary(lmer_otusize2) # singular fit
```

```
## Linear mixed model fit by maximum likelihood  ['lmerMod']
## Formula: log.inputDNA ~ log.OTUsize + (1 | sample/OTU)
##    Data: dd
## 
##      AIC      BIC   logLik deviance df.resid 
##   8370.1   8402.2  -4180.1   8360.1     4534 
## 
## Scaled residuals: 
##     Min      1Q  Median      3Q     Max 
## -1.5870 -0.8972  0.2585  0.8621  1.5082 
## 
## Random effects:
##  Groups     Name        Variance Std.Dev.
##  OTU:sample (Intercept) 0.0000   0.0000  
##  sample     (Intercept) 0.0000   0.0000  
##  Residual               0.3693   0.6077  
## Number of obs: 4539, groups:  OTU:sample, 892; sample, 7
## 
## Fixed effects:
##              Estimate Std. Error t value
## (Intercept) -0.859244   0.024433  -35.17
## log.OTUsize -0.004252   0.004382   -0.97
## 
## Correlation of Fixed Effects:
##             (Intr)
## log.OTUsize -0.929
## optimizer (nloptwrap) convergence code: 0 (OK)
## boundary (singular) fit: see help('isSingular')
```

```
lmer_otusize3 <- lmer(log.inputDNA ~ log.OTUsize + 
                        (log.OTUsize | sample/OTU), 
                      data =dd, REML = F)
```

```
## boundary (singular) fit: see help('isSingular')
```

```
summary(lmer_otusize3) # singular fit
```

```
## Linear mixed model fit by maximum likelihood  ['lmerMod']
## Formula: log.inputDNA ~ log.OTUsize + (log.OTUsize | sample/OTU)
##    Data: dd
## 
##      AIC      BIC   logLik deviance df.resid 
##   8375.0   8432.7  -4178.5   8357.0     4530 
## 
## Scaled residuals: 
##     Min      1Q  Median      3Q     Max 
## -1.7035 -0.8994  0.2381  0.8605  1.7488 
## 
## Random effects:
##  Groups     Name        Variance            Std.Dev.     Corr 
##  OTU:sample (Intercept) 0.00000000000000000 0.0000000000      
##             log.OTUsize 0.00000000000007144 0.0000002673  NaN 
##  sample     (Intercept) 0.00481833652171563 0.0694142386      
##             log.OTUsize 0.00016272280876906 0.0127562851 -1.00
##  Residual               0.36863288024147645 0.6071514475      
## Number of obs: 4539, groups:  OTU:sample, 892; sample, 7
## 
## Fixed effects:
##              Estimate Std. Error t value
## (Intercept) -0.852596   0.036120  -23.61
## log.OTUsize -0.004729   0.006572   -0.72
## 
## Correlation of Fixed Effects:
##             (Intr)
## log.OTUsize -0.966
## optimizer (nloptwrap) convergence code: 0 (OK)
## boundary (singular) fit: see help('isSingular')
```

```
ranova(lmer_otusize2) # remove random factor
```

```
## boundary (singular) fit: see help('isSingular')
## boundary (singular) fit: see help('isSingular')
```

```
## ANOVA-like table for random-effects: Single term deletions
## 
## Model:
## log.inputDNA ~ log.OTUsize + (1 | OTU:sample) + (1 | sample)
##                  npar  logLik    AIC LRT Df Pr(>Chisq)
## <none>              5 -4180.1 8370.1                  
## (1 | OTU:sample)    4 -4180.1 8368.1   0  1          1
## (1 | sample)        4 -4180.1 8368.1   0  1          1
```

```
ranova(lmer_otusize3) # remove random factor
```

```
## boundary (singular) fit: see help('isSingular')
## boundary (singular) fit: see help('isSingular')
```

```
## ANOVA-like table for random-effects: Single term deletions
## 
## Model:
## log.inputDNA ~ log.OTUsize + (log.OTUsize | OTU:sample) + (log.OTUsize | sample)
##                                           npar  logLik    AIC    LRT Df
## <none>                                       9 -4178.5 8375.0          
## log.OTUsize in (log.OTUsize | OTU:sample)    7 -4178.5 8371.0 0.0000  2
## log.OTUsize in (log.OTUsize | sample)        7 -4180.1 8374.1 3.1665  2
##                                           Pr(>Chisq)
## <none>                                              
## log.OTUsize in (log.OTUsize | OTU:sample)     1.0000
## log.OTUsize in (log.OTUsize | sample)         0.2053
```

```
# the random factors should be omitted from the model, but sample/OTU are part of the design, so i must keep those two

# test fixed effect 
lmer_otusize4 <- lmer(log.inputDNA ~ 1 + (1 | sample/OTU), 
                      data =dd, REML = F)
```

```
## boundary (singular) fit: see help('isSingular')
```

```
anova(lmer_otusize3, lmer_otusize4) # fixed effect log.OTUsize not sig, p = 0.534
```

```
## Data: dd
## Models:
## lmer_otusize4: log.inputDNA ~ 1 + (1 | sample/OTU)
## lmer_otusize3: log.inputDNA ~ log.OTUsize + (log.OTUsize | sample/OTU)
##               npar    AIC    BIC  logLik deviance  Chisq Df Pr(>Chisq)
## lmer_otusize4    4 8369.1 8394.7 -4180.5   8361.1                     
## lmer_otusize3    9 8375.0 8432.7 -4178.5   8357.0 4.1078  5      0.534
```

```
# parameter estimate REML=T and including the non-sig log.inputDNA to get an estimate of the low R2 value
lmer_otusize5 <- lmer(log.inputDNA ~ log.OTUsize + 
                        (1 | sample/OTU), 
                      data =dd, REML = T)
```

```
## boundary (singular) fit: see help('isSingular')
```

```
summary(lmer_otusize5)
```

```
## Linear mixed model fit by REML ['lmerMod']
## Formula: log.inputDNA ~ log.OTUsize + (1 | sample/OTU)
##    Data: dd
## 
## REML criterion at convergence: 8376.7
## 
## Scaled residuals: 
##     Min      1Q  Median      3Q     Max 
## -1.5867 -0.8970  0.2585  0.8619  1.5079 
## 
## Random effects:
##  Groups     Name        Variance Std.Dev.
##  OTU:sample (Intercept) 0.0000   0.0000  
##  sample     (Intercept) 0.0000   0.0000  
##  Residual               0.3695   0.6079  
## Number of obs: 4539, groups:  OTU:sample, 892; sample, 7
## 
## Fixed effects:
##              Estimate Std. Error t value
## (Intercept) -0.859244   0.024438  -35.16
## log.OTUsize -0.004252   0.004383   -0.97
## 
## Correlation of Fixed Effects:
##             (Intr)
## log.OTUsize -0.929
## optimizer (nloptwrap) convergence code: 0 (OK)
## boundary (singular) fit: see help('isSingular')
```

```
r.squaredGLMM(lmer_otusize5) # 0.0002 0.0002
```

```
##               R2m          R2c
## [1,] 0.0002073445 0.0002073445
```

```
# there is little reason to check residuals, as the model fit is so poor, but here they are
sresid <- resid(lmer_otusize5, type = "pearson") # Extract the standardised residuals
hist(sresid) # poor residuals
```

```
plot(lmer_otusize5)
```

```
plot(sresid ~ log.OTUsize, data = dd)
```

```
pred.mm2 <- ggpredict(lmer_otusize5, terms = c("log.OTUsize[all]"))

(p2 <- ggplot() +
  geom_line(data = pred.mm2, 
            aes(x = x, y = predicted), 
            colour = "black", size = 1) +
  geom_point(data = dd, 
             aes(x = log.OTUsize, y = log.inputDNA, 
                 colour = OTU), size = 0.8) +
  geom_smooth(data = dd, 
              aes(x = log.OTUsize, y = log.inputDNA, group = OTU),
              method = "lm", se = FALSE, size = 0.1, 
              colour = "black", alpha = 0.5, linetype = 1) +
  labs(x = "log(non-spike-corrected OTU size)", 
       y = "log(input genomic DNA mass)") +
  theme_cowplot() + 
  theme(legend.position = "none") + 
  coord_cartesian(ylim = c(-2.0, 0)) +
  facet_wrap(~sample2))
```

```
## `geom_smooth()` using formula 'y ~ x'
```

```
p2 | p1 + plot_annotation(tag_levels = 'A')
```

```
## `geom_smooth()` using formula 'y ~ x'
## `geom_smooth()` using formula 'y ~ x'
```

###### Model-based pipeline-noise estimator: offset0 (ã from Eq. 2)

To remove a source of terminological ambiguity, i change the varname
sample to trap, because from the point of view of the estimator model,
the *samples* are the 6 dilutions made from each trap. The 6
dilutions are scaled in the var inputDNA (0.16 to 1.00), where 1.00
means undiluted and 0.16 is most diluted.

```
dd2 <- dd %>% 
  select(OTUsize, sample, dilution, inputDNA, OTU, lysisBufferFrac,
         readSpikeSum) %>% 
  pivot_wider(names_from = OTU, values_from = OTUsize, 
              values_fill = 0) %>% 
  rename(trap = sample)

# readSpikeSum is greater in samples with low inputDNA (i.e. those that are diluted more). However, a complicating factor here is that we change the amount of spike-in added, and this dataset has three dilution series with lots of spike-in added and 4 series with lower amounts of spike-in added. Also, the decline in readSpikeSum is not smooth, which we attribute to experimental error.
ggplot(data = dd2, aes(x = inputDNA, y = readSpikeSum, 
                       colour = trap, shape = trap)) +
  geom_point() +
  geom_smooth()
```

```
## `geom_smooth()` using method = 'loess' and formula 'y ~ x'
```

```
## Warning in simpleLoess(y, x, w, span, degree = degree, parametric =
## parametric, : Chernobyl! trL>n 6

## Warning in simpleLoess(y, x, w, span, degree = degree, parametric =
## parametric, : Chernobyl! trL>n 6
```

```
## Warning in sqrt(sum.squares/one.delta): NaNs produced
```

```
## Warning in stats::qt(level/2 + 0.5, pred$df): NaNs produced
```

```
## Warning in simpleLoess(y, x, w, span, degree = degree, parametric =
## parametric, : Chernobyl! trL>n 6

## Warning in simpleLoess(y, x, w, span, degree = degree, parametric =
## parametric, : Chernobyl! trL>n 6
```

```
## Warning in sqrt(sum.squares/one.delta): NaNs produced
```

```
## Warning in stats::qt(level/2 + 0.5, pred$df): NaNs produced
```

```
## Warning in simpleLoess(y, x, w, span, degree = degree, parametric =
## parametric, : Chernobyl! trL>n 6

## Warning in simpleLoess(y, x, w, span, degree = degree, parametric =
## parametric, : Chernobyl! trL>n 6
```

```
## Warning in sqrt(sum.squares/one.delta): NaNs produced
```

```
## Warning in stats::qt(level/2 + 0.5, pred$df): NaNs produced
```

```
## Warning in simpleLoess(y, x, w, span, degree = degree, parametric =
## parametric, : Chernobyl! trL>n 6

## Warning in simpleLoess(y, x, w, span, degree = degree, parametric =
## parametric, : Chernobyl! trL>n 6
```

```
## Warning in sqrt(sum.squares/one.delta): NaNs produced
```

```
## Warning in stats::qt(level/2 + 0.5, pred$df): NaNs produced
```

```
## Warning in simpleLoess(y, x, w, span, degree = degree, parametric =
## parametric, : Chernobyl! trL>n 6

## Warning in simpleLoess(y, x, w, span, degree = degree, parametric =
## parametric, : Chernobyl! trL>n 6
```

```
## Warning in sqrt(sum.squares/one.delta): NaNs produced
```

```
## Warning in stats::qt(level/2 + 0.5, pred$df): NaNs produced
```

```
## Warning in simpleLoess(y, x, w, span, degree = degree, parametric =
## parametric, : Chernobyl! trL>n 6

## Warning in simpleLoess(y, x, w, span, degree = degree, parametric =
## parametric, : Chernobyl! trL>n 6
```

```
## Warning in sqrt(sum.squares/one.delta): NaNs produced
```

```
## Warning in stats::qt(level/2 + 0.5, pred$df): NaNs produced
```

```
## Warning in simpleLoess(y, x, w, span, degree = degree, parametric =
## parametric, : Chernobyl! trL>n 6

## Warning in simpleLoess(y, x, w, span, degree = degree, parametric =
## parametric, : Chernobyl! trL>n 6
```

```
## Warning in sqrt(sum.squares/one.delta): NaNs produced
```

```
## Warning in stats::qt(level/2 + 0.5, pred$df): NaNs produced
```

```
## Warning: The shape palette can deal with a maximum of 6 discrete values because
## more than 6 becomes difficult to discriminate; you have 7. Consider
## specifying shapes manually if you must have them.
```

```
## Warning: Removed 6 rows containing missing values (geom_point).
```

```
## Warning in max(ids, na.rm = TRUE): no non-missing arguments to max; returning
## -Inf

## Warning in max(ids, na.rm = TRUE): no non-missing arguments to max; returning
## -Inf

## Warning in max(ids, na.rm = TRUE): no non-missing arguments to max; returning
## -Inf

## Warning in max(ids, na.rm = TRUE): no non-missing arguments to max; returning
## -Inf

## Warning in max(ids, na.rm = TRUE): no non-missing arguments to max; returning
## -Inf

## Warning in max(ids, na.rm = TRUE): no non-missing arguments to max; returning
## -Inf

## Warning in max(ids, na.rm = TRUE): no non-missing arguments to max; returning
## -Inf
```

```
# option to filter the dataset by subsets of trap
dd3 <- dd2 %>% filter(trap %in% 
                        c("700239.M1.S1", "357256.M2.S1", "124031.M1.S1", "286789.M1.S1", "123545.M1.S1", "HB.053.M1.S1", "HB.216.M1.S1"))
# "700239.M1.S1", "357256.M2.S1", "124031.M1.S1", "286789.M1.S1", "123545.M1.S1", "HB.053.M1.S1", "HB.216.M1.S1"

# "123545.M1.S1", "HB.053.M1.S1", "HB.216.M1.S1" are the samples with high numbers of spike-in reads

# if you want to see only a few traps
ggplot(data = dd3, aes(x = inputDNA, y = readSpikeSum, 
                       colour = trap, shape = trap)) +
  geom_point() +
  geom_smooth()
```

```
## `geom_smooth()` using method = 'loess' and formula 'y ~ x'
```

```
## Warning in simpleLoess(y, x, w, span, degree = degree, parametric =
## parametric, : Chernobyl! trL>n 6
```

```
## Warning in simpleLoess(y, x, w, span, degree = degree, parametric =
## parametric, : Chernobyl! trL>n 6
```

```
## Warning in sqrt(sum.squares/one.delta): NaNs produced
```

```
## Warning in stats::qt(level/2 + 0.5, pred$df): NaNs produced
```

```
## Warning in simpleLoess(y, x, w, span, degree = degree, parametric =
## parametric, : Chernobyl! trL>n 6

## Warning in simpleLoess(y, x, w, span, degree = degree, parametric =
## parametric, : Chernobyl! trL>n 6
```

```
## Warning in sqrt(sum.squares/one.delta): NaNs produced
```

```
## Warning in stats::qt(level/2 + 0.5, pred$df): NaNs produced
```

```
## Warning in simpleLoess(y, x, w, span, degree = degree, parametric =
## parametric, : Chernobyl! trL>n 6

## Warning in simpleLoess(y, x, w, span, degree = degree, parametric =
## parametric, : Chernobyl! trL>n 6
```

```
## Warning in sqrt(sum.squares/one.delta): NaNs produced
```

```
## Warning in stats::qt(level/2 + 0.5, pred$df): NaNs produced
```

```
## Warning in simpleLoess(y, x, w, span, degree = degree, parametric =
## parametric, : Chernobyl! trL>n 6

## Warning in simpleLoess(y, x, w, span, degree = degree, parametric =
## parametric, : Chernobyl! trL>n 6
```

```
## Warning in sqrt(sum.squares/one.delta): NaNs produced
```

```
## Warning in stats::qt(level/2 + 0.5, pred$df): NaNs produced
```

```
## Warning in simpleLoess(y, x, w, span, degree = degree, parametric =
## parametric, : Chernobyl! trL>n 6

## Warning in simpleLoess(y, x, w, span, degree = degree, parametric =
## parametric, : Chernobyl! trL>n 6
```

```
## Warning in sqrt(sum.squares/one.delta): NaNs produced
```

```
## Warning in stats::qt(level/2 + 0.5, pred$df): NaNs produced
```

```
## Warning in simpleLoess(y, x, w, span, degree = degree, parametric =
## parametric, : Chernobyl! trL>n 6

## Warning in simpleLoess(y, x, w, span, degree = degree, parametric =
## parametric, : Chernobyl! trL>n 6
```

```
## Warning in sqrt(sum.squares/one.delta): NaNs produced
```

```
## Warning in stats::qt(level/2 + 0.5, pred$df): NaNs produced
```

```
## Warning in simpleLoess(y, x, w, span, degree = degree, parametric =
## parametric, : Chernobyl! trL>n 6

## Warning in simpleLoess(y, x, w, span, degree = degree, parametric =
## parametric, : Chernobyl! trL>n 6
```

```
## Warning in sqrt(sum.squares/one.delta): NaNs produced
```

```
## Warning in stats::qt(level/2 + 0.5, pred$df): NaNs produced
```

```
## Warning: The shape palette can deal with a maximum of 6 discrete values because
## more than 6 becomes difficult to discriminate; you have 7. Consider
## specifying shapes manually if you must have them.
```

```
## Warning: Removed 6 rows containing missing values (geom_point).
```

```
## Warning in max(ids, na.rm = TRUE): no non-missing arguments to max; returning
## -Inf

## Warning in max(ids, na.rm = TRUE): no non-missing arguments to max; returning
## -Inf

## Warning in max(ids, na.rm = TRUE): no non-missing arguments to max; returning
## -Inf

## Warning in max(ids, na.rm = TRUE): no non-missing arguments to max; returning
## -Inf

## Warning in max(ids, na.rm = TRUE): no non-missing arguments to max; returning
## -Inf

## Warning in max(ids, na.rm = TRUE): no non-missing arguments to max; returning
## -Inf

## Warning in max(ids, na.rm = TRUE): no non-missing arguments to max; returning
## -Inf
```

```
otu <- dd3 %>% 
  select(starts_with("OTU"))

X <- dd3 %>% 
  select(!starts_with("OTU"))
```

```
otu <- mvabund(otu)

# fit0 <- manyglm(otu ~ 1, family = "negative.binomial") # if no predictors
# fit0 <- manyglm(otu ~ soup, family = binomial("cloglog")) # for p/a data
fit0 <- manyglm(otu ~ inputDNA + trap, # including trap to 
                family = "negative.binomial", 
                data = X) 
# offset(log(lysisBufferFrac)) is to correct  
plot(fit0)
```

```
offset0 <- log(rowSums(otu)) - log(rowSums(fitted(fit0))) 
rowsum <- rowSums(otu)
dd3$offset0 <- offset0
dd3$rowsum <- rowsum
dd3 <- dd3 %>% 
  relocate(c(rowsum, offset0), .after = readSpikeSum)

p3 <- ggplot(data = dd3, aes(x = offset0, y = log(readSpikeSum), 
                       group = trap, 
                       colour = trap,
                       shape = trap)) +
  geom_point() +
  geom_smooth(method = "lm", se = FALSE, size = 0.6, 
              alpha = 0.5, linetype = 1) +
  labs(x = "offset0", y = "ln(readSpike)") +
  theme_cowplot() +
  theme(legend.position = "right")

p4 <- ggplot(data = dd3, aes(x = inputDNA, y = rowsum, 
                       group = trap, 
                       colour = trap,
                       shape = trap)) +
  geom_point() +
  geom_smooth(method = "lm", se = FALSE, size = 0.6, 
              alpha = 0.5, linetype = 1) +
  # labs(x = "inputDNA", y = "ln(readSpike)") +
  theme_cowplot() +
  theme(legend.position = "right")

p4 / p3
```

```
## `geom_smooth()` using formula 'y ~ x'
```

```
## Warning: The shape palette can deal with a maximum of 6 discrete values because
## more than 6 becomes difficult to discriminate; you have 7. Consider
## specifying shapes manually if you must have them.
```

```
## Warning: Removed 6 rows containing missing values (geom_point).
```

```
## `geom_smooth()` using formula 'y ~ x'
```

```
## Warning: The shape palette can deal with a maximum of 6 discrete values because
## more than 6 becomes difficult to discriminate; you have 7. Consider
## specifying shapes manually if you must have them.

## Warning: Removed 6 rows containing missing values (geom_point).
```

```
summary(lm(log(readSpikeSum) ~ offset0, data = dd3))
```

```
## 
## Call:
## lm(formula = log(readSpikeSum) ~ offset0, data = dd3)
## 
## Residuals:
##     Min      1Q  Median      3Q     Max 
## -2.4357 -1.1276 -0.1006  0.9715  2.6733 
## 
## Coefficients:
##             Estimate Std. Error t value            Pr(>|t|)    
## (Intercept)   9.2748     0.2293  40.457 <0.0000000000000002 ***
## offset0       0.4507     0.6894   0.654               0.517    
## ---
## Signif. codes:  0 '***' 0.001 '**' 0.01 '*' 0.05 '.' 0.1 ' ' 1
## 
## Residual standard error: 1.483 on 40 degrees of freedom
## Multiple R-squared:  0.01057,    Adjusted R-squared:  -0.01416 
## F-statistic: 0.4274 on 1 and 40 DF,  p-value: 0.517
```

```
# R2 = 0.03062
```
